# Hotspot propensity across mutational processes

**DOI:** 10.1101/2022.09.14.507952

**Authors:** Claudia Arnedo-Pac, Ferran Muiños, Abel Gonzalez-Perez, Nuria Lopez-Bigas

## Abstract

The ability to study mutation rate variability at nucleotide resolution is impaired by the sparsity of observed mutational events across the genome. To circumvent this problem, here we investigated the propensity of 14 different mutational processes to form recurrently mutated sites across tumour samples (hotspots). We found that mutational signatures 1 (SBS1) and 17 (SBS17a and SBS17b) have the highest propensity to form hotspots, generating 5-78 times more than other common somatic mutational processes. After accounting for trinucleotide mutational probabilities, sequence composition and heterogeneity of mutation rates at 10 Kbp, the majority (89-95%) of SBS17a and b hotspots remain unexplained. This suggests that local genomic features play a significant role in SBS17a and b hotspot propensity, among which we identify CTCF binding as a minor contributor. In the case of SBS1, we demonstrate that including genome-wide distribution of methylated CpGs sites into our models can explain most (80-100%) of its hotspot propensity. We also observe an increased hotspot propensity of SBS1 in normal tissues from mammals, as well as in *de novo* germline mutations. We demonstrate that hotspot propensity is a useful readout to assess the accuracy of mutation rate models at nucleotide resolution. This new approach and the findings derived from it open up new avenues for a range of somatic and germline studies investigating and modelling mutagenesis.

## Introduction

Mutations are heterogeneously distributed along genomes, following the interplay between DNA damage and DNA repair. Their specific interaction defines what we know as mutational processes, with heterogeneous activity along the nucleotide sequence. Understanding these mechanisms underlying somatic and germline mutagenesis, in humans and other species, has profound implications for the biological and biomedical sciences.

At the large (e.g., megabase) scale, the variability of mutation rates of different mutational processes is known to be associated with chromatin features such as DNA accessibility, transcriptional activity and replication timing^1–4^. Such covariates have been successfully used to model neutral mutagenesis at this “low” resolution^4^. At a more local scale, the mutation rate depends upon nucleotide sequence composition^5–7^, which is also included in such models^4, 8^, as well as upon local ––up to a few Kbp–– scale chromatin features including nucleosome occupancy^9^, transcription factor binding^10–14^, and non-canonical DNA secondary structures^15, 16^. A few of these local chromatin features are known to interact with particular mutational processes to increase mutation rates within specific nucleotides. This is the case of UV-light damage at the binding sites of ETS-family of transcription factors^17–19^ and APOBEC3A mutations at ssDNA loops within DNA hairpins^20–22^. Despite the knowledge accumulated to date on the genomic features that influence the mutation rate, the ultimate goal of determining how much and why the rate of mutation contributed by different processes vary at single base resolution (i.e., at each unique position in the genome) remains a major challenge^20^.

The current lack of accurate estimates of the mutation rate at base resolution is primarily caused by the sparsity of observed mutations across the 3 billion base pairs (bp) of the human genome, despite pooling the thousands of whole genomes currently available in the public domain. For the same reason, we have not yet been able to comprehensively characterise the determinants of the variability of the mutation rate at this scale^23, 24^. We are thus in need of novel ways to study mutagenesis at single base resolution.

Here, we propose to study mutation rate variability at single base resolution through the analysis of mutational hotspots, that is, recurrent mutations affecting the exact same genomic position across samples. This idea is based on the hypothesis that hotspots will reflect, apart from random sampling effects due to the finite size of the genome, genuine differences in the underlying mutation probability per unique genomic site for a given mutational process. We thus posit that tracking the rate of formation of hotspots will allow us to explore, firstly, which mutational processes have a higher mutation rate variability at single base resolution. And, secondly, to quantify how much of this variability can be explained by large and small scale determinants of mutagenesis.

In this study, we leveraged the somatic mutations of more than 7,500 whole genome sequences of tumours from 49 cancer types, and systematically detected and quantified the mutational processes creating passenger hotspots. We discovered that mutational processes active across tumours exhibit very different propensities to form hotspots, with signatures 1 (SBS1) and 17 (SBS17a and SBS17b) generating 5-78 times more hotspots than other common mutational processes when controlling for differences in their activities across tumours. We found that trinucleotide mutational probabilities, sequence composition and heterogeneity of mutation rates at 10 Kbp only explain a fraction of hotspot propensity among mutational signatures, ranging from 5% in SBS17 to 59% in SBS4. Conversely, by including genome-wide distribution of methylated CpGs sites into SBS1 models, we get to explain 80-100% of SBS1 hotspot propensity. The high hotspot propensity of SBS1 is also observed in normal tissue samples from mammals, as well as across *de novo* germline mutations in the human population. Our work provides a new metric to study mutation rate variability at nucleotide resolution, highlighting the current difficulty to accurately estimate and model mutation rate variability across most mutational processes, with the exception of SBS1. These findings have strong implications for studies of basic mutagenesis as well as for those measuring positive or negative selection in cancer and evolution.

## Results

### Mutational hotspots across cancers

We developed a new method, named HotspotFinder, to identify and annotate unique genomic positions that are recurrently mutated (two or more times; i.e., hotspots) to the same alternate (e.g., two C>T transitions) across the whole-genomes of 7,507 sequenced primary and metastatic tumours from 49 cancer types (Fig. 1a-d; Supplementary Table S1 and S2; Methods; Supplementary Note 1). To avoid false positive hotspots due to sequencing, mapping or somatic mutation calling errors^25^, careful filters of problematic genomic areas and positions containing population variants were applied (Methods; Supplementary Note 1). Similarly, we excluded potential hotspots caused by positive selection by filtering out mutations overlapping the coding –and surrounding non-coding– sequence of known cancer driver genes^26, 27^ (Fig. 1e; Supplementary Table S3; Methods).

**Fig. 1.**
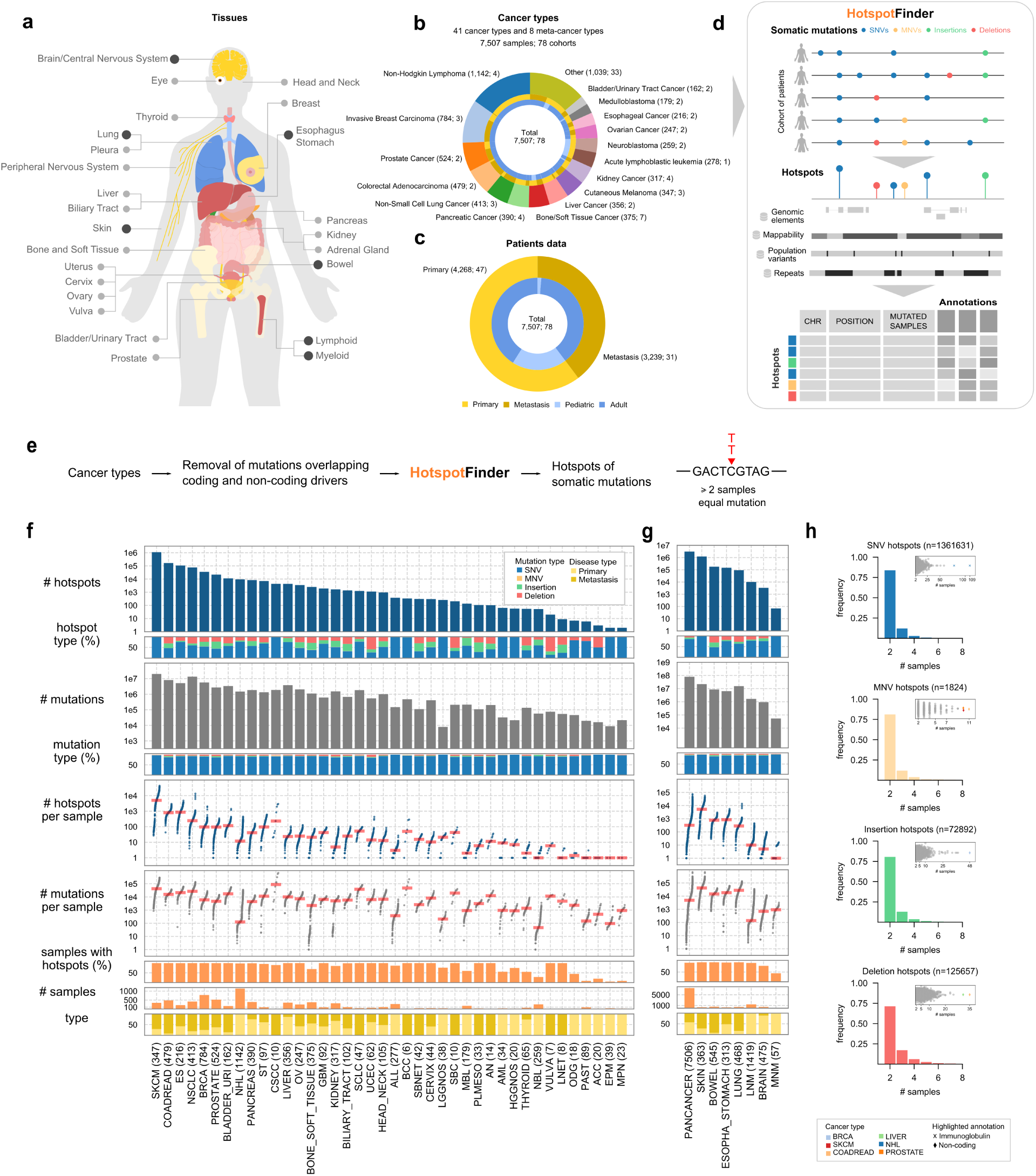
**Identification of hotspots across cancers. a**) Cancer types analysed depicting specific and meta-cancer types in light and dark grey, respectively. **b**) Number of patients and sequencing cohorts among specific cancer types. **c**) Detail of the number of primary, metastatic, adult and paediatric tumours analysed. **d**) Schematic overview of HotspotFinder, a new algorithm to identify hotspots of somatic mutations (Methods; Supplementary Note 1). **e**) Overview of the steps for hotspots identification. **f**) Summary of total hotspots identified across specific cancer types and **g**) meta-cancer types with at least one hotspot. Number of samples per cancer type after filtering out driver–overlapping mutations are shown in parenthesis. Note that 1 sample belonging to ALL (and PANCANCER) was excluded from the analysis since no mutations outside driver regions were left after filtering. **h**) Histograms of hotspot size (number of mutated samples per hotspot) considering only hotspots from specific cancer types. Embedded dotplots show hotspot sizes per individual hotspot, where the shape and the colour represents the overlapping genomic element and the cancer type where the hotspot was identified, respectively. Cancer types are listed as follows: Acute Lymphoblastic Leukemia (ALL), Acute Myeloid Leukemia (AML), Adrenocortical Carcinoma (ACC), Anal Cancer (AN), Basal Cell Carcinoma (BCC), Biliary Tract (BILIARY_TRACT), Bladder/Urinary Tract (BLADDER_URI), Bone/Soft Tissue (BONE_SOFT_TISSUE), Bowel (BOWEL), CNS/Brain (BRAIN), Cervix (CERVIX), Colorectal Adenocarcinoma (COADREAD), Cutaneous Melanoma (SKCM), Cutaneous Squamous Cell Carcinoma (CSCC), Endometrial Carcinoma (UCEC), Ependymoma (EPM), Esophageal cancer (ES), Esophagus/Stomach cancers (ESOPHA_STOMACH), Glioblastoma Multiforme (GBM), Head and Neck (HEAD_NECK), High-Grade Glioma NOS (HGGNOS), Invasive Breast Carcinoma (BRCA), Kidney (KIDNEY), Liver (LIVER), Low-Grade Glioma NOS (LGGNOS), Lung (LUNG), Lung Neuroendocrine Tumor (LNET), Lymphoid Neoplasm (LNM), Medulloblastoma (MBL), Myeloid Neoplasm (MNM), Myeloproliferative Neoplasms (MPN), Neuroblastoma (NBL), Non-Hodgkin Lymphoma (NHL), Non-Small Cell Lung Cancer (NSCLC), Oligodendroglioma (ODG), Ovarian Cancer (OV), Pan-cancer (PANCANCER), Pancreas (PANCREAS), Pilocytic Astrocytoma (PAST), Pleural Mesothelioma (PLMESO), Prostate (PROSTATE), Retinoblastoma (RBL), Skin (SKIN), Small Bowel Cancer (SBC), Small Bowel Neuroendocrine Tumor (SBNET), Small Cell Lung Cancer (SCLC), Stomach cancer (ST), Thyroid (THYROID), Vulva (VULVA).

A total of 1,562,004 alternate-specific hotspots of four different types of mutations were identified across individual cancer types (3,106,161 across the pan-cancer cohort): 1,361,631 corresponded to SNVs (87.2%), 125,657 to deletions (8.0%), 72,892 to insertions (4.7%), and 1,824 to MNVs (0.1%) (Fig. 1f,g; Supplementary Table S4). Hotspots covered roughly 0.13% of the mappable hg38 reference genome (approximately 2,439 Mbp after excluding driver-associated regions; see Methods) and the vast majority (99.35%) were located in non-coding regions. In all cancer types except retinoblastomas at least one hotspot was observed. The majority of hotspots were small, comprising 2 or 3 mutated samples (Fig. 1h, Supp. Fig. 1), with few exceptions (Fig. 1h, Supp. Fig. 1-2). Hotspots of insertions and deletions were particularly abundant (at similar or greater rates than SNVs hotspots) across cancer types with active indel mutational processes (i.e., 19.3% and 30.1% insertion and deletion hotspot frequency in colorectal tumours, respectively) (Fig. 1f,g; Supp. Fig. 3). High rates of hotspots of insertions and deletions across other tumour types with few samples may be due to specific mutational processes such as mismatch repair deficiency, or sequencing and/or calling errors and biases across cohorts as shown in other studies^28^. Given that the vast majority of hotspots identified were composed of SNVs, and because errors in indels mapping and calling could lead to higher rates of false positive insertion and deletion hotspots^28^, we decided to focus on the study of SNV hotspots formation. Henceforth, we use hotspot as synonymous with SNV hotspot, unless otherwise specified.

**Fig. 2.**
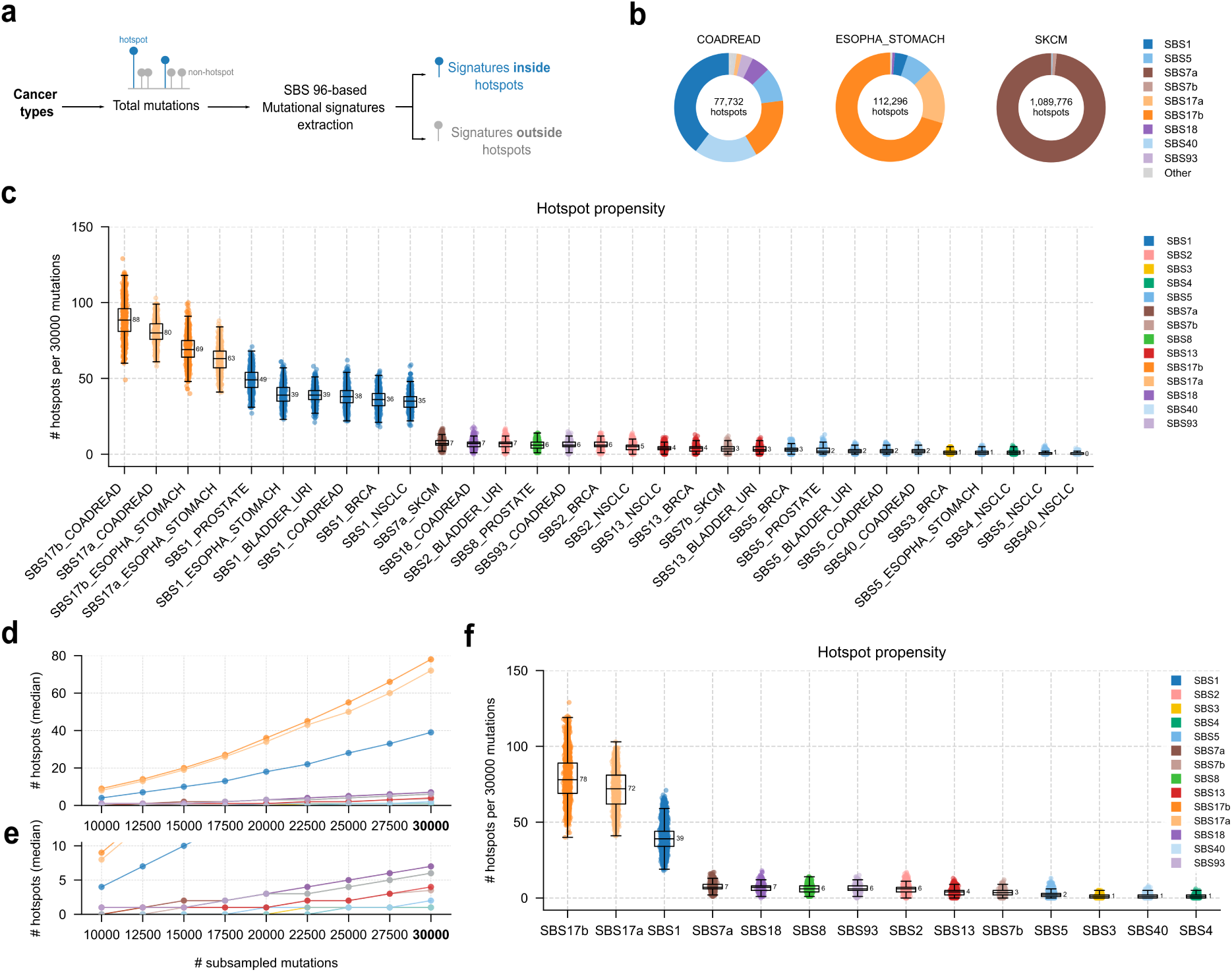
Mutational processes with increased propensity to form hotspots. a. ) Graphical definition of the analysis of mutational signatures in the sets of mutations inside and outside hotspots. **b**) Pie charts depicting the relative number of hotspots observed per signature in the cancer type. **c**) Number of hotspots per signature and cancer type observed by subsampling 30,000 total mutations (300 mutations/sample, 100 samples) within each group in the set of mappable megabases (Methods). **d**) Median number of observed hotspots per signature across subsamples at different mutation burden (100-300 mutations/sample, 100 samples) (Methods). In order to obtain signature-level estimates, subsamples across different cancer types were merged as listed in Methods. **e**) Zoom into 0-10 hotspots from d. **f**) Number of hotspots per signature observed within 30,000 subsampled mutations (300 mutations/sample, 100 samples) across cancer types merging data shown in c. Error bars in c and f show 1.5 times the IQR below and above 1st and 3rd quartiles, respectively. Signatures are sorted in descending order according to the mean number of observed hotspots.

**Fig. 3.**
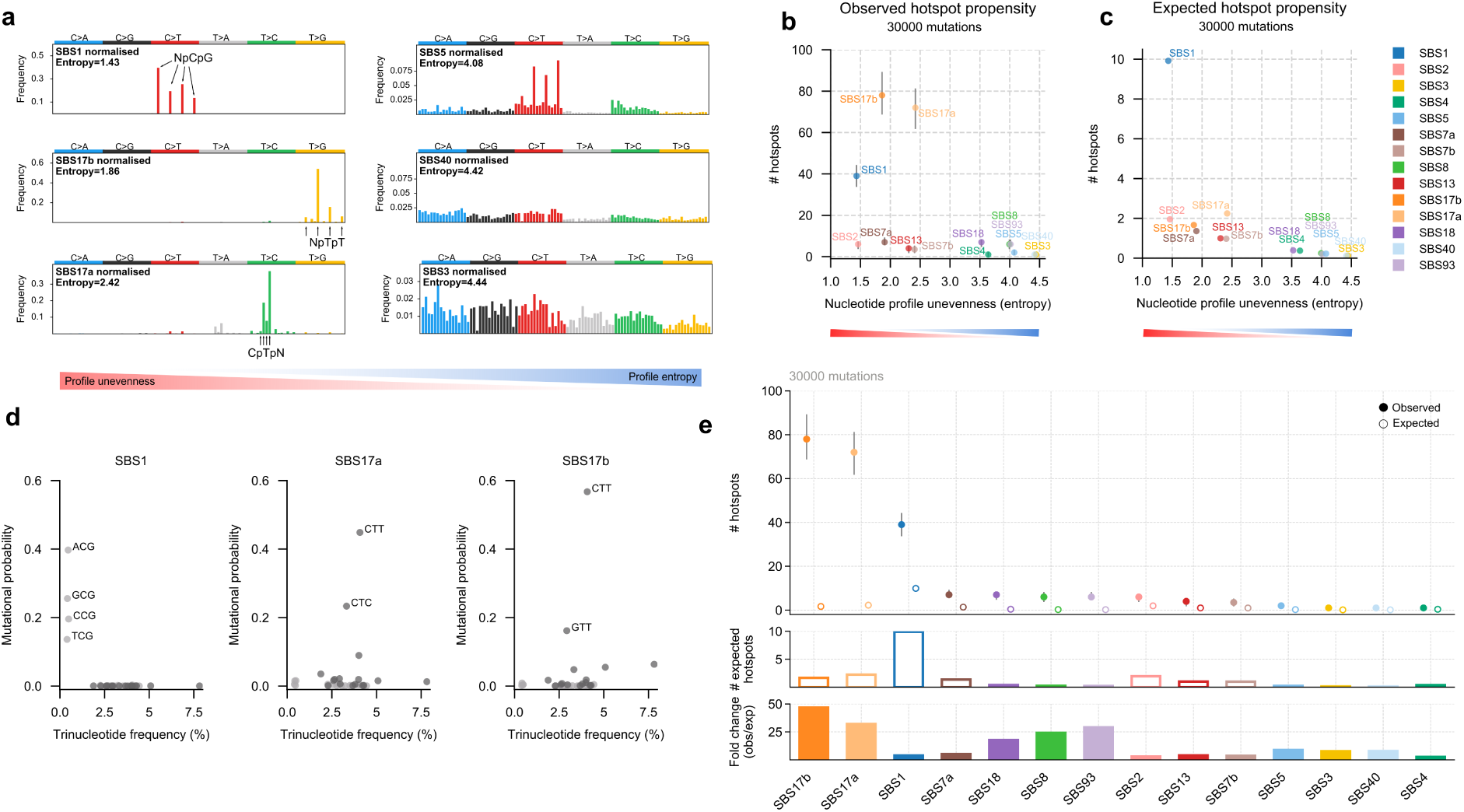
**Contribution of trinucleotide mutational probabilities and trinucleotide abundance to hotspots formation across signatures. a**) Normalised trinucleotide profiles of SBS1, SBS17a, SBS17b, SBS40, SBS5 and SBS3; additional signatures are shown in Supp. Fig. 19a. **b**) Number of observed hotspots (y axis) versus the entropy of the normalised signature profile (x axis). Observed hotspot propensity was computed by subsampling 30,000 total mutations (300 mutations/sample, 100 samples) within mappable megabases (Methods). Error bars show the interquartile ranges (IQR) of the number of hotspots observed across subsamples. **c**) Number of expected hotspot propensity (y axis) versus the entropy of the normalised signature profile (x axis). Expected data was generated using the model of homogeneous distribution of trinucleotide-specific mutation rates across the genome for 300 mutations/sample and 100 samples across mappable megabases, therefore it is comparable to the data in b. **d**) Mutational probability of trinucleotides per signature versus their frequency within mappable megabases. The mutational probability for each trinucleotide was obtained by merging those from the respective three alternates given by the normalised signature profile. **e**) Comparison of observed versus expected hotspot propensity per signature (top-middle) and the fold change of observed versus expected number of hotspots (bottom). Observed dots show median hotspots; error bars show the IQR. Observed and expected data correspond to that shown in b and c.

While the number of hotspots per cancer type increased with sample size and mutation burden (Spearman’s R=0.93, p=9e-18 and R=0.72, p=2e-7, respectively; Supp. Fig. 4a,b), similarly as predicted by theoretical models of homogeneous mutation rates across trinucleotides (Supp. Fig. 4c,d), hotspots appeared at different rates across tumour types (Supp. Fig. 5-6). Computing the number of SNVs required to generate one hotspot (hotspot conversion rates) showed large variability across cancer types, ranging from 21 mutations in melanomas to 3,036 mutations in medulloblastomas (Supp. Fig. 7). This variability in conversion rates, together with the small fraction of hotspots shared between cancer types (Supp. Fig. 8), suggested that hotspots could capture differences in mutation rate heterogeneity at single base resolution across mutational processes active in distinct tissues.

**Fig. 4.**
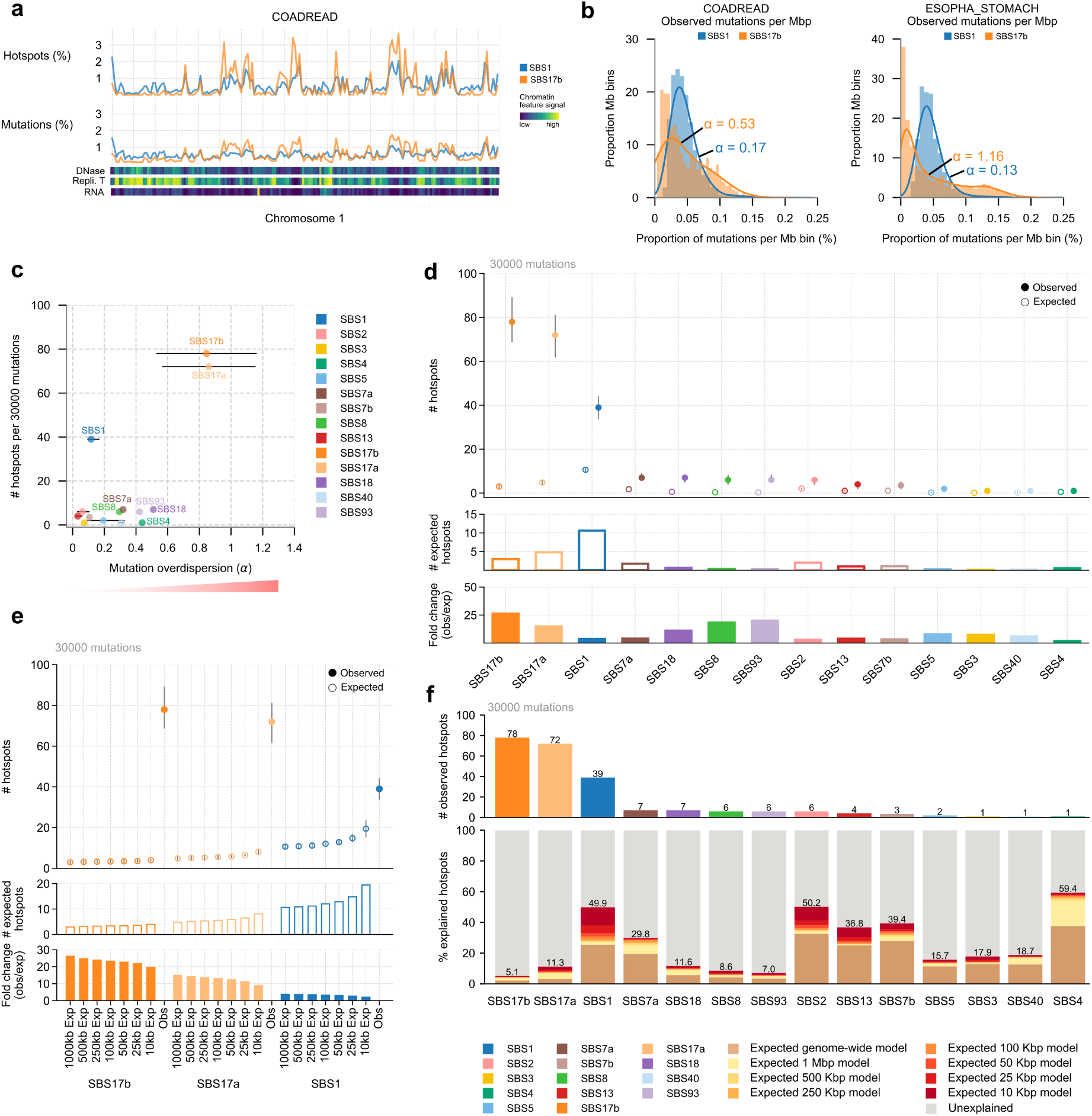
**Contribution of large scale unevenness to hotspots formation across signatures. a**) Proportion of observed hotspots (top) and mutations (bottom) in colorectal cancers attributable to SBS1 and SBS17b across mappable megabases of chromosome 1. Normalised epigenomic signals of chromatin accessibility, replication timing and expression per megabase are shown below. **b**) Distribution of the observed proportion of mutations across mappable megabases. Alpha values show the overdispersion of the negative binomial distribution fitted with the mutation counts per megabase (Methods). **c**) Number of observed hotspots versus the overdispersion (unevenness) of mutation counts within genomic megabases. Observed hotspots were computed by subsampling 30,000 total mutations (300 mutations/sample, 100 samples) within mappable megabases as shown in Fig. 4b. **d**) Comparison of observed versus the expected theoretical number of hotspots per signature (top-middle) calculated with the megabase model accounting for trinucleotide composition and large-scale mutation rate variability. Expected data was generated for 300 mutations/sample and 100 samples across mappable megabases. Dots show the median number of hotspots across cancers. Error bars correspond to the IQR. The fold change of observed versus expected number of hotspots is shown below. **e**) Comparison of observed versus expected hotspot propensity per signature calculated across different bin sizes ranging from 1 Mbp to 10 Kbp long. Observed and expected dots show median hotspots; error bars show the IQR. **f**) Median observed hotspot propensity using 30,000 subsampled mutations (top). Fraction of the median observed hotspot propensity that is accounted for by the different expected hotspot propensity models (bottom). The total percentage of hotspot propensity explained by the 10 Kbp-based model is shown for each signature.

**Fig. 5.**
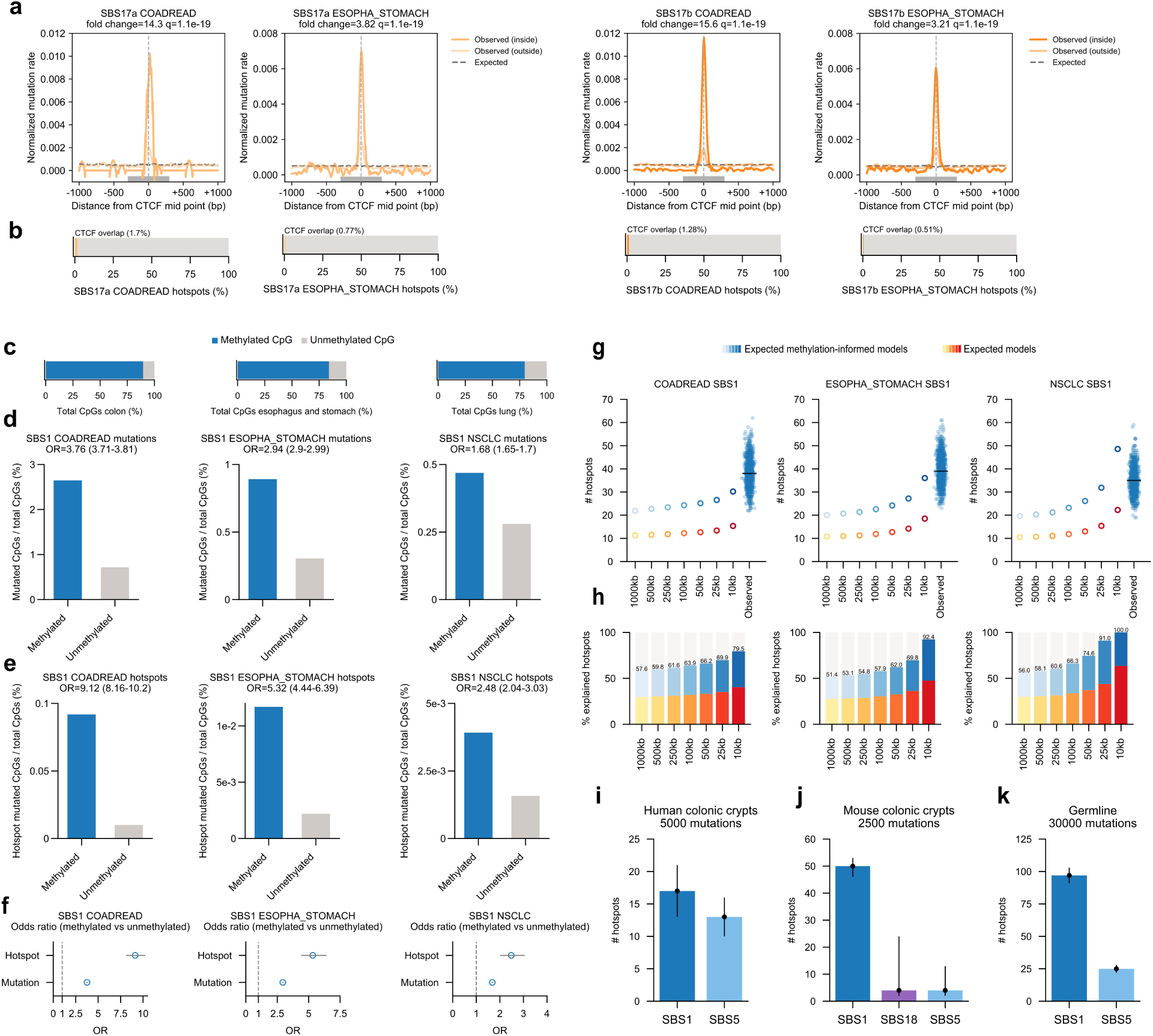
**Underlying mechanisms of hotspot formation. a**) Piled-up normalised mutation rate of SBS17a and b across CTCF binding sites (600 bp; grey rectangle) and their flanking 5’ and 3’ regions (700 bp each). Mutations are split as overlapping hotspots (observed inside) and non-overlapping hotspots (observed outside). The fold change of inside mutation rates in CTCF binding sites versus flanks are shown on top. Significance of the fold change is computed by randomly distributing inside mutations across CTCF and flanking sequences considering the signature’s trinucleotide mutational probability and the trinucleotide sequence composition (Methods). **b**) Proportion of SBS17a and b hotspots in the cancer type overlapping CTCF binding sites. **c**) Frequency of methylated and unmethylated CpGs in normal tissue epigenomes. **d**) Percentage of SBS1 mutated CpG sites among total methylated CpGs (blue) and total unmethylated CpGs (grey) in a tissue-matched epigenome. The odds ratio and its 95% confidence interval of mutations in methylated versus unmethylated CpGs is shown on top. **e**) Percentage of CpGs overlapped by SBS1 hotspots in methylated and unmethylated CpGs, computed as in panel d for mutations. **f**) Odds ratio for SBS1 hotspots and mutations in methylated versus unmethylated CpGs sites per cancer type. Error bars represent the 95% confidence interval. **g**) Comparison of observed versus expected SBS1 hotspot propensity. Orange dots depict the expected hotspot propensity using 1 Mbp to 10 Kbp models. Blue dots depict the expected hotspot propensity using methylation-informed 1 Mbp to 10 Kbp models. Observed hotspot propensity is shown as filled blue dots, where each dot represents a subsample (n=1,000). The median hotspot propensity is shown in black. Note that the expected hotspot propensity within 10 Kb methylation-informed models is included in the range of the distribution of observed hotspot propensities for each cancer type. **h**) Percentage of explained hotspot propensity resulting from the expected models shown in g compared to the median observed hotspot propensity in the cancer type. Percentage of explained hotspots by methylation-informed models is shown on top. **i**) Hotspot propensity in human colonic crypts computed using a total of 5,000 mutations per signature (25 samples and 250 mutations/sample). **j**) Hotspot propensity in mouse colonic crypts computed using a total of 2,500 mutations per signature (25 samples and 100 mutations/sample). **k**) Hotspot propensity in human *de novo* germline mutations computed using a total of 30,000 mutations per signature (30,000 random mutations from the merged datasets per subsample). Black dots in panels i, j and k show the median hotspot propensity across 1,000 subsamples; error bars represent the IQR.

### Propensity of mutational processes to form hotspots

To study the variability of the mutation rate at single nucleotide resolution for different mutational processes, we first analysed the enrichment of hotspots across different sequence contexts. We observed that, across cancer types, hotspots were differentially enriched for different types of nucleotide changes, and within particular trinucleotide sequences (Supp. Fig. 9-10). A similar observation was found for indel hotspots (Supp. Fig. 11).

We then estimated the number of hotspots generated by single base substitution (SBS) mutational signatures in each tumour type by conducting a *de novo* extraction and decomposition into the COSMIC (v3.2 GRCh38) signatures to facilitate comparisons across cancers (Fig. 2a,b; Supp. Fig. 12, 13; Methods; Supplementary Note 2, 3). SBS1 appeared as an important contributor of hotspots across all cancer types, particularly in tumours of the brain (69.6% of hotspots), pancreas (60.9%; Fig. 2b), prostate (42.3%) and colorectum (39.7%; Fig. 2b), in agreement with the enrichment of hotspots in C>T transitions in the NpCpG context (Supp. Fig. 9-10). SBS17a and b contributed a large proportion of hotspots in esophageal cancers (16.8% and 72.3% of hotspots, respectively), stomach tumours (12.1% and 67.4%), and, to a lesser degree, of those in colorectal cancers (1.51% and 18.6%) (Fig. 2b; Supp. Fig. 9), in accordance with hotspot enrichment for T>G transversions and T>C transitions, especially within CpTpT contexts in gastrointestinal tumours (Supp. Fig. 9-10). SBS5, an age-related signature of unknown aetiology ubiquitous across cancer types, SBS2 and SBS13 (APOBEC-related), SBS4 (related to tobacco smoking), and SBS7a (caused by UV light damage) were responsible for a high proportion of hotspots of specific cancer types (Fig. 2b; Supp. Fig. 13). The analysis of the extended context (up to 21 bp) of hotspots per signature showed that despite small preferences for specific nucleotides outside the trinucleotide context (e.g., SBS7a and b^7^, SBS17a and b^29, 30^, and SBS93), the contribution of the trinucleotide sequence clearly dominates all of them, suggesting it plays the preponderant role in the formation of hotspots as is the case with mutations^7^ (Supp. Fig. 14).

The burden of hotspots contributed by each mutational signature showed, as expected, a good correlation with its activity and the proportion of samples at which it was found active (Supp. Fig. 15). In order to account for this in our analysis, we set out to implement a new metric, the propensity of a signature to form hotspots, which informs about its intrinsic inclination to generate hotspots independently of their overall cohort activity and sample-wise contribution to the mutation burden. The larger the hotspot propensity of a mutational process, the higher the variability of its mutation rates at single base resolution.

To test this new metric, we selected 14 mutational signatures with high activity in one or more cancer types (detailed criteria in Methods) and re-ran hotspot identification upon subsampling a fixed number of mutations contributed by each of them. Across a range of 10,000-30,000 mutations sampled from 100 tumours (at equal number of mutations per sample), we observed approximately 1 to 2 orders of magnitude more hotspots contributed by SBS17b, SBS17a and SBS1 than by the other eleven mutational signatures studied across tumour types (Fig. 2c-f). Specifically, for 30,000 mutations across 100 samples, a median of 78, 72 and 39 hotspots were observed for SBS17b, SBS17a and SBS1, respectively (Fig. 2f). Conversely, SBS7a, SBS18, SBS2, SBS93, SBS8, SBS7b, and SBS13 contributed 3-7 hotspots, and SBS5, SBS40, SBS3 and SBS4 generated 1-2 hotspots (Fig. 2f). That is, SBS17b, SBS17a and SBS1 contributed 5-78 times more hotspots than the other signatures under the same conditions. Equivalent results were observed with larger sampling (Supp. Fig. 16). In an orthogonal ––subsampling independent–– calculation of the propensity to form hotspots employing the fold change of mutations contributed by each signature across tumours inside and outside hotspots, we obtained very similar results (Supp. Fig. 17-18; Supplementary Note 4). In summary, SBS17b, SBS17a and SBS1 show the highest propensity to form hotspots among mutational processes commonly active in human tissues.

### Signature profile unevenness and trinucleotide abundance affect hotspot formation

We hypothesised that one possible reason for the disparity in the propensity to form hotspots of different mutational processes could be the number of genomic positions available to each of them. This is determined, in the first place, by the particular trinucleotide mutational probabilities of the signature that give rise to a particular shape (i.e., skewness or unevenness vs uniformity or evenness) of its trinucleotide profile. We measured the unevenness of the activity of signatures across trinucleotides as the entropy of their mutational profile normalised by their abundance in the human genome, which informs about the mutational probability of the signature across trinucleotides irrespective of the genome composition (Supplementary table S5). The more uneven the profile of a signature, the lower its entropy (Fig. 3a; Supp. Fig. 19a). The three signatures with the highest propensity to form hotspots across tissues (SBS1, SBS17a and SBS17b) possess low entropy profiles (Fig. 3b; Methods). Conversely, signatures with even profiles, such as SBS3 and SBS5, did not display a high propensity to form hotspots. These results suggest that the fewer active trinucleotides in a mutational signature (the more uneven its profile), the more likely it is that two mutations contributed by the process map to the same genomic position (Fig. 3b).

The availability of the 96 trinucleotides in the human genome must also influence the propensity of different mutational signatures to form hotspots. To investigate the combined effect of genomic trinucleotide abundance and signatures prolife unevenness, we determined the expected (based solely on the trinucleotide substitution probabilities) number of hotspots formed by mutations contributed by several processes (see details in Methods and Supplementary Note 5). Following this assumption, 4.4-78.4 times more hotspots were expected to be contributed by SBS1 than by any other signature (Fig. 3c; Supp. Fig. 19b). The explanation for this is that the four NpCpG trinucleotides targeted by SBS1 are comparatively depleted in the human genome (0.4-0.5 % NpCpG vs 1.9-7.9 % non-NpCpG in mappable bins; Fig. 3d; Supp. Fig. 19c). In contrast, the NpTpT and CpTpN trinucleotides, which respectively concentrate the activity of SBS17b and SBS17a, show average (or above average) representation in the human genome (Fig. 3d). Interestingly, for all mutational signatures analysed, we observed more hotspots than expected, with the highest differences for SBS17b and SBS17a (Fig. 3e; Methods). Since the number of hotspots computed via the theoretical model only accounts for the unevenness of the mutational profile of the signature and the abundance of trinucleotides in the genome, we sought to quantify how other variables contribute to hotspot propensities.

### Hotspot propensity is differentially impacted by large-scale chromatin features across signatures

We reasoned that another factor underlying the differences in hotspot propensity across signatures could be the uneven distribution of their mutations along the human genome (Supp. Fig. 20). Several megabase scale chromatin features, such as replication time and chromatin compaction are known to play a role in this variability of the mutation rate along the genome^2–4, 31–34^. To compute if the distribution of mutations at large scales influences the propensity of different signatures to form hotspots, we counted the number of mutations and hotspots contributed by each signature in autosomal mappable bins of length 1 Mbp. We found that the density of hotspots was positively correlated with that of mutations at the megabase scale (Supp. Fig. 21), as illustrated along chromosome 1 for SBS1 and SBS17b in colorectal cancers (Fig. 4a). As expected, hotspot density per megabase correlated with chromatin accessibility, replication time and level of transcription (Fig. 4a; Supp. Fig. 22-24).

In order to quantify the potential effect of the large-scale genomic variability of a signature’s activity in the formation of hotspots, we first computed the overdispersion of the distribution of its mutation rates across megabase bins (Fig. 4b; Methods). SBS1 mutations exhibited low overdispersion across megabase genomic bins. Others, like SBS17b, showed large variability in mutation counts at the megabase scale (Fig. 4b; Supp. Fig. 25). Actually, SBS1, SBS17a and SBS17b, the mutational processes with the highest propensity to form hotspots, appeared at opposite ends of the spectrum of megabase mutation overdispersion across all tested signatures (Fig. 4c). SBS17a and b exhibit the highest megabase mutation rate unevenness, followed by SBS18 and SBS4 (Fig. 4c). While differences in the interplay of mutational processes with chromatin features may underlie the dissimilar unevenness observed across signatures, it is worth noticing that the NpTpT trinucleotides targeted by SBS17a and SBS17b show a greater inter-megabase variability than the NpCpG targeted by SBS1 (Supp. Fig. 19d).

Next, we compared the number of hotspots observed across 1 Mbp segments of the genome with that expected after accounting for the megabase distribution of mutations and the trinucleotide composition of each segment (Fig. 4d; Methods). While the expected number of hotspots across signatures (in particular for SBS4) increased compared to that computed using only the signature mutational profile and the trinucleotide genome composition (Fig. 3e), the observed-to-expected hotspot fold-change was still greater than 1 for all signatures. In other words, the megabase-scale distribution of mutations contributed by different signatures has a partial influence on their propensity to form hotspots.

We thus asked whether genomic features that affect the rate of mutations at scales smaller than 1 Mbp could explain part of the unaccounted hotspot propensity. To that end, we repeated the calculation of expected hotspots at sub-megabase bins of lengths 500, 250, 100, 50, 25 and 10 Kbp. Interestingly, we observed a monotonic increase of the expected number of hotspots as the bin size decreased across signatures (Fig. 4e, Supp. Fig. 26), which is in line with the increase in the overdispersion of mutations (Supplementary table S7). We next estimated the increasing contribution of large-scale covariates, on top of mutational probabilities of a signature and the trinucleotide composition of the genome, to the observed hotspot propensity. Nearly 50% of SBS1 hotspot propensity can be explained by the 10 Kbp models (Fig. 4f). The corresponding fraction is between 30 and 60% for SBS4, SBS2, SBS13, SBS7a and b. For other mutational processes, conversely, reducing the scale of the genomic bins at which the mutation rate variability is assessed does not result in appreciable increase in the fraction of hotspots explained by the models. Specifically, 88.7% and 94.9% of SBS17a and b hotspots, respectively, still remain unexplained, a similar rate as for SBS8, SBS18 and SBS93. Extending the subsampling experiment to 60,000 mutations showed equivalent explainability gaps of hotspot propensities (Supp. Fig. 27). In summary, our results suggest that large-scale chromatin features, down to 10 Kbp, play distinct roles on hotspot formation across mutational processes. Although known mutation rate determinants can predict up to 60% of hotspot rates for some mutational processes, the knowledge gap is still noticeable across signatures (40.6-94.9%).

### Determinants of signature 17 hotspot propensity are largely unknown

Signatures SBS17a and b showed the greatest hotspot propensity, which remains unexplained, for the most part, by well-known determinants of mutation rate variability. We hypothesised that additional chromatin features at the local ––below the 10 Kbp–– scale^23^ may have an important influence on their hotspot propensity. We found that SBS17a and b hotspots appeared increased in colorectal (14.3 and 15.6 times) and esophageal-stomach tumours (3.82 and 3.21 times) at CTCF binding sites with respect to their flanking sequences, significantly beyond the expected given their sequence composition (Fig. 5a). These results are consistent with prior reports of CTCF binding sites bearing clusters of SBS17 mutations^13, 14, 34^. Nevertheless, only 1.7% and 0.8% of SBS17a hotspots, and 1.3% and 0.5% of SBS17b hotspots in colorectal and esophageal-stomach cancers, respectively, overlap CTCF binding sites (Fig. 5b), although the majority of CTCF-overlapping hotspots in these two tumour types (86% and 53.5%) are attributed to the combined activity of these two signatures. Therefore, despite the contribution of CTCF binding sites to the expected hotspot rate, other still unidentified small scale genomic features must also bear increased rate of SBS17a and b hotspot propensity.

**Signature 1 hotspot propensity can be explained by tissue-matched methylation data** Similarly, we sought to identify additional factors to improve our estimates of SBS1 mutation rate variability at nucleotide resolution. Since the aetiology of SBS1 involves the spontaneous deamination of 5-methylcytosines^6^, we reasoned that the differential methylation of CpG sites across the genome could be the reason behind the remaining unexplained hotspot propensity (Fig. 4f). Indeed, SBS1 hotspots, as well as mutations, observed across colorectal, esophageal-stomach and non-small cell lung cancers are enriched for CpG sites that appear methylated in the respective tissues of origin (Fig. 5c-e). Motivated by this result, we next computed the expected hotspot propensity across these three tumour types adding the methylation status of CpGs to the features already considered in the models. We found that, as the size of the genomic bins decreased, these methylation-aware models produced numbers of expected hotspots that approached the distribution of the observations for 30,000 random sampled mutations (Fig. 5g). Between 80% and 100% of the observed SBS1 hotspot propensity is ultimately explained by methylation-aware models at 10 Kbp bins (Fig. 5h).

### Signatures 1 and 17 high hotspot propensity in non-malignant somatic and germline tissues

SBS1 is ubiquitously active across normal somatic tissues of humans and other mammals^35, 36^, and it has been shown to also contribute *de novo* mutations to the human germ line^35, 37^. We thus reasoned that its increased hotspot propensity should be detectable across these normal tissues as it is across human tumours. To test this, we collected somatic mutations identified across normal colonic crypts from human and mouse^36^, and *de novo* germline mutations identified across 7 datasets of family pedigrees^37–42^. Employing the subsampling strategy described above (between 2,500 and 30,000 mutations depending on the mutation rate and sample size in each dataset; Methods), we found a median of 18 observed hotspots of SBS1 across human colonic crypts and 50 across mouse colonic crypts (Fig. 5i,j), and 97 across *de novo* human germline mutations (Fig. 5k). Similarly, we found high hotspot propensity of SBS17a and b in Barrett’s oesophagus pre-malignant lesions^43^ (Supp. Fig. 28). Thus, the mechanisms driving the clustering of mutations contributed by SBS17a/b and SBS1 along the genome are not restricted to cancer cells, but operate also in non-malignant tissues, including germ cells in the case of SBS1.

In summary, SBS17a/b and SBS1 are the signatures with the largest hotspot propensity. In the case of SBS17a and b, only 5-11% of their hotspot propensity can be explained by their mutational profile and the distribution of its mutations across 10 Kbp, suggesting that small-scale genomic features, including CTCF binding and others that remain to be identified, play a key role in their hotspot propensity. On the contrary, we demonstrate that 80-100% of SBS1 observed hotspot propensity is driven by the relative scarcity and unequal distribution of methylated CpGs along the genome at 10 Kbp scale. Bearing in mind that the rate of hotspots is a readout of the variability of the mutation rate at base pair resolution, these findings are relevant for the correct modelling of SBS1 mutation rate to assess its contribution to the mutations in normal tissues of humans and mammals, and *de novo* mutations entering the germline.

## Discussion

To study the variability of the mutation rate at single nucleotide resolution, here we introduced a new metric, the hotspot propensity, which tracks the intrinsic tendency of mutational processes to form hotspots after correcting for differences in activity across cohorts and single samples. Measuring the hotspot propensity of 14 mutational signatures commonly active across tissues showed striking variability (up to 78 times fold) between them, with SBS1, SBS17a and SBS17b exhibiting the largest propensity to form mutational hotspots. The mutations of SBS1, the ubiquitous clock-like mutational process attributed to 5-methylcytosine deamination^5, 6^, show a high propensity to form hotspots across most tissues analysed. SBS17a and b, of unknown aetiology, stand out as highly hotspot-prone both in primary esophageal, stomach and colorectal tumours^29^. In our dataset, SBS17b hotspots are also observed in metastases of patients exposed to capecitabine/5-fluorouracil as part of the treatment of their primary tumours^30, 44^ (Supplementary Note 2). We corroborated the increased propensity to form hotspots –albeit at smaller rates than SBS1 and SBS17a/b– of the mutations of other signatures, including the UV-light caused SBS7a^10, 11, 17–19^ and the APOBEC related SBS2 ^20–22^. We envisage that the increasing availability of cancer genome sequences, coupled with improvements in mutation calling and filtering of false positive calls, will pave the way for the exploration of hotspot propensity of other types of mutations not covered within our study (e.g., indels).

We have also leveraged the measurement of hotspot propensity to explore how much currently known determinants of mutagenesis actually contribute to the variability of the mutation rate at single nucleotide resolution. To do this, we built models of the expected hotspot propensity with increasing complexity –i.e., including additional known or suspected determinants of the mutation rate– across signatures and then compared them with the observed propensity. We were thus able to compute the accumulated contribution of these known or suspected large and small scale determinants of hotspot formation. The unevenness of the mutational profile of a signature, and the abundance of different trinucleotides in the human genome show a high contribution to the hotspot propensity of some mutational signatures. Notably, the high hotspot propensity of SBS1 is underpinned by its high activity at NpCpG trinucleotides, which are relatively depleted in the human genome (due to SBS1 action throughout evolution^9, 45^). Determinants of the mutation rate at relatively large scale ––between 10 Kbp and 1 Mbp–– show a differential contribution to hotspot propensity across signatures. For some signatures, like SBS4, SBS2 and SBS1, large-scale seems to have a larger contribution than for others, such as SBS17a and SBS17b. Nevertheless, the sequence composition of the genome, the mutational profile of a signature, and the large scale determinants of the mutation rate only explain a fraction, ranging from 5.1% in SBS17b to 59.4% in SBS4, of the observed hotspot propensity. This finding quantifies our knowledge gap of the determinants of the rate of mutations for different processes at nucleotide base resolution.

In order to help closing this gap, we explored the contribution of additional features to the high hotspot propensity of SBS17 and SBS1. We corroborated that the mutational hotspots of SBS17a/b are enriched for CTCF binding sites, although only a small fraction of these hotspots (0.5-1.7%) overlap CTCF binding sites. Thus indicating that other unknown factors, below the 10 Kbp scale, play a major role in the propensity of SBS17 mutations to form hotspots. For example, it has recently been suggested that nucleotide excision repair could play a role in SBS17 mutagenesis^46^. The situation presents in a totally different light for the hotspot propensity of SBS1 mutations. Careful calculation of the number of SBS1 expected hotspots accounting for its mutational profile, its mutation rate at 10 Kbp and the uneven distribution of methylation at CpG sites along the genome explain the vast majority (80%-100%) of hotspots observed across three different tissues. In other words, our analysis allowed us to close the gap of explainability in the variability of the mutation rate of SBS1 at single nucleotide resolution. We foresee that the methodology described in the manuscript to compute the hotspot propensity of the mutations of different signatures will be applied to explore other potential new determinants of mutagenesis for signatures beyond SBS1, in particular, the elusive SBS17a/b.

While here we have focused on the study of the variability of the mutation rate at single nucleotide resolution in tumours, some of the mutational processes studied are also active in somatic healthy tissues and in the germline. A salient example is the spontaneous deamination of 5-methylcytosine, underlying SBS1^5, 6^, a universal age-related process affecting not only human somatic cells, but also human germ cells^35^, and the tissues of other species^36, 47^. We show that, as is the case in colorectal tumours, a high propensity of hotspots may be observed for SBS1 mutations occurring in the normal colonic crypts of human and mouse. Likewise, SBS1 hotspots are found across *de novo* germline mutations. This suggests that the findings shown for SBS1 constitute a universal feature of somatic and germline tissues.

The driving motivation of this work was the systematic exploration of the variability of the mutation rate at single nucleotide resolution and its causes, which we undertook through the calculation of the propensity of the mutations of different processes to form hotspots. While the fine-grained variability of the mutation rate exploiting coding neutral hotspots has been previously explored across cancer genomes^20, 25, 48^, to the best of our knowledge, this constitutes the first time hotspot propensity is exploited as a readout of mutation rate variability at nucleotide resolution. This high-resolution understanding of the mutation rate variability has different implications. First of all, it provides a new approach to study potential determinants of mutagenesis, as we have showcased with CpG methylation for SBS1. In the field of cancer genomics, statistical tools to identify signals of positive selection across the genome rely on the accuracy of background (i.e., neutral) mutagenesis. While state-of-the-art background models based on large and small scale features have been successful in the identification of protein coding cancer drivers genes^4, 8^, this is still challenging for non-coding regions^49, 50^ and for individual cancer driver mutations^51^. The findings shown here could contribute to measuring the uncertainty of current background models at nucleotide resolution. Similarly, the analysis of evolutionary trajectories in cancer, normal tissues, and across evolution rely on the modelling of mutation rates at single nucleotide resolution. These analyses could also benefit from improving our estimates of the propensity of the different mutational processes to create hotspots as well as from studying new determinants of mutagenesis. We envision that our work constitutes a step forward in the pursuit of these long-term goals.

## Methods

### Cohorts

We collected somatic mutations from 78 cohorts of whole genome sequenced cancer patients included in IntOGen^27^ (release 1 February 2020). Cohorts contained primary and metastatic tumours from adult and paediatric individuals, encompassing a total of 7,507 samples and 83,410,018 somatic mutations. Detailed information about each sequencing cohort and information on how to download them can be found at Supplementary Table S1 and www.intogen.org.

### Pre-processing of cohorts

In order to homogenise the datasets for our analysis and minimise the number of false mutation calls, we conducted the following pre-processing on individual cohorts as follows:

– Liftover of somatic mutations to hg38 reference genome. Mutations in those cohorts that used hg19 as reference genome were lifted over to hg38 using pyliftover package version 0.3 (pypi.org/project/pyliftover/) as described in ^27^. Only mutations that mapped to hg38 were kept for analysis.
– Filtering of somatic mutations: we removed mutations that a) fell outside of autosomal or sexual chromosomes; b) had the same reference and alternate nucleotides; c) had a reference nucleotide that did not match the annotated hg38 reference nucleotide; d) had an unknown nucleotide ––a nucleotide not corresponding to A, C, G, T–– in their trinucleotide or pentanucleotide reference sequence, as stated by their start position; e) were classified as complex indels ––indels that are a mixture of insertions and deletions such as GTG>GAAA.
– Filtering of germline variants: our analysis aimed for the identification of hotspots of somatic mutations. In order to decrease contamination of somatic calls by unfiltered germline mutations, we removed mutations overlapping population variants. Briefly, we removed mutations overlapping genomic positions with one or more polymorphic variants (i.e., allele frequency equal or greater than 1%) (see “Mappable genome and high mappability genomic bins” section for complete details).
– Filtering of low mappability sequences: non-mappable regions (i.e., repetitive or non-unique sequences in the genome) are prone to sequencing artefacts. To control such errors, we discarded mutations that 1) fell outside high mappability regions and/or 2) overlapped blacklisted regions of low mappability (see “Mappable genome and high mappability genomic bins” section for complete details).
– Filtering of hypermutated samples: from each cohort, we filtered out hypermutated samples, this is, samples that carried more than 10,000 mutations and exceeded 1.5 times the interquartile range over the 75^th^ percentile, as described in ^27^.

### Cancer types classification

WGS samples were merged into 49 cancer types comprising one or more individual cohorts. Cancer type classification was based on the Memorial Sloan Kettering Cancer Center (MSKCC) OncoTree^52^ (2021-11-02 release, available at oncotree.mskcc.org). Assignment of each cohort to the different cancer type levels in the OncoTree hierarchy was carried out using the available clinical information of the cohort and can be found in Supplementary Table S1. Ad-hoc cancer types were added in those cases where the OncoTree classification did not fulfil the cohort definition. In order to avoid redundant cancer types ––entities containing the same or very similar set of samples–– within our analysis, we simplified the resulting hierarchy into two different levels A (specific) and B (meta-cancer type): level A entities were the most specific annotation available for a group of samples (e.g., melanomas); when two or more level A entities could be merged together according to the hierarchy, a level B annotation was added (e.g., melanomas, basal cell carcinomas, and cutaneous squamous cell carcinoma were grouped in skin cancers). Finally, all samples were merged into the Pancancer level. Cancer types included in the analysis are listed in Supplementary Table S2.

### Pre-processing of cancer types

In order to identify the presence of multiple samples originating from the same donor, we conducted a systematic analysis of shared mutations among samples in each cohort and cancer type. Briefly, for every sample in a dataset, we computed the number of equal mutations with any other sample in the group and divided it over the total number of mutations of both samples in the comparison. Samples with more than 10% of shared mutations with any other sample were flagged for manual review. Two samples from M_OS were found to have primary tumour samples sequenced in D_OS and were subsequently removed from the metastatic cohort M_OS.

### Driver gene annotations

Cancer driver genes were collected from two sources: the Compendium of Cancer Genes from the driver discovery pipeline IntOGen^27^ (release 1 February 2020) and the COSMIC Cancer Gene Census^26^ (CGC) (downloaded on 24-08-2021). The Compendium of Cancer Genes is composed of genes with experimental and/or *in silico* protein-coding driver evidence (n=568 genes). CGC list contains expert-curated genes with experimental driver evidence from sporadic and familial cancers. Only those genes annotated as somatic and having a cancer role different from fusion partners were included (n=589 genes). Our final set of driver genes consisted of 782 genes as listed in Supplementary table S3. The genomic coordinates of the coding and surrounding non-coding sequences of driver genes were obtained from Gencode^53^ v35 comprehensive gene annotation file at ftp.ebi.ac.uk/pub/databases/gencode/Gencode_human/release_35/gencode.v35.annotation.gtf. gz (downloaded on 30-08-2020) (Supplementary Note 1).

### Mappable genome and high mappability genomic bins

We defined the mappable genome as the fraction of the reference hg38 genome included in our analysis (total nucleotides = 2,439,219,900 bp; nucleotides with known trinucleotide context = 2,439,219,170 bp). The mappable genome consisted of regions of high mappability that did not overlap with 1) blacklisted sequences of low mappability, 2) genomic positions containing population variants, and 3) coding or non-coding regions of driver genes (see “Driver gene annotations” and Supplementary Note 1 for further details). Regions of high mappability (≥ 0.9) based on 100-mer pileup mappability were computed for hg38 reference genome using The GEnomic Multi-tool^54^ (GEM) mappability software version 2013-04-06. BED files containing hg38 blacklisted regions of low mappability were obtained from the ENCODE Unified GRCh38 Blacklist (downloaded from encodeproject.org/files/ENCFF356LFX on 16-06-2020). Positions containing population variants were defined as those overlapping any substitution or short indel with total variant allele frequency above 1% as identified by gnomAD^55^ version 3.0 (downloaded from gnomad.broadinstitute.org on 25-06-2020). Next, we defined a set of high mappable megabases (1 Mbp) based on their overlap to the mappable genome. We first obtained hg38 genomic bins by partitioning chromosome coordinates in consecutive non–overlapping chunks of 1 Mb length. For each bin, we computed the sequence overlap with the mappable genome using the Python library pybedtools^56, 57^ and kept those bins within autosomes whose sequence overlap with the mappable genome was above the first quartile of the distribution of fractional sequence overlap across megabase bins (Q1 = 0.80 fractional overlap; n = 2,196 bins) (total nucleotides = 2,012,091,302 bp; nucleotides with known trinucleotide context = 2,012,091,115). For the set of mappable 1 Mbp bins, we obtained sets of submegabase bins of lengths 500 Kbp (n = 4,392 bins), 250 Kbp (n = 8,784 bins), 100 Kbp (n = 21,960 bins), 50 Kbp (n = 43,920 bins), 25 Kbp (n = 87,840 bins), and 10 Kbp (n = 219,600 bins) bins by partitioning individual 1 Mbp bins into consecutive chunks. All hg38 nucleotide sequences were retrieved from the Python package bgreference version 0.6.

### Identification of hotspots of somatic mutations

Genome-wide recurrently mutated positions from independent samples were identified using the new algorithm HotspotFinder version 1.0.0 (Supplementary Note 1), freely available at bitbucket.org/bbglab/hotspotfinder. Hotspots of the four mutation types (SNVs, MNVs, insertions and deletions) were analysed separately. For each cancer type, HotspotFinder was run over the set of filtered mutations after excluding those overlapping coding or non-coding sequences (5’UTR, 3’UTR, splice sites, introns, proximal and distal promoters; Supplementary Note 1) of driver elements. Hotspots were identified as single positions in the genome that contained i) 2 or more mutations of equal alternates (e.g., two C>T transitions) or ii) 2 or more mutations of different alternates (e.g., C>T and C>G). All the analyses included in the present work were carried out using hotspots of equal alternate. Hotspots were annotated with the default mappability, population variants and genomic regions provided within the method (Supplementary Note 1) and those non-overlapping genomic elements were kept. All other parameters were set as default.

### Hotspot burden modelling

We modelled the relationship between hotspot burden and mutation burden per sample for each cancer type with univariate ordinary least squares (OLS) regression models using the Python package statsmodels^58^.

### Estimation of conversion rates

Conversion rates or the number of mutations to observe 1 hotspot were calculated for cancer types with more than 100 individuals through a subsampling experiment. For 1,000 times, we selected 100 random individuals (without replacement) and pooled their SNVs to identify hotspots of equal alternate as previously explained. We then modelled the number of hotspots per individual against their observed mutation burden using OLS regression models^58^. Conversion rates were computed as the inverse of the regression slope of significant models (p<0.05) for those cancer types with at least 750 significant linear models across random replicates.

### Enrichment of substitution types in hotspots

Mutations overlapping and non-overlapping hotspots (mutations inside and outside hotspots, respectively) were classified into 6 and 96 pyrimidine-based substitution types based on GRCh38 reference genome using SigProfilerMatrixGenerator^59^ version 1.1.26. Hotspot enrichments for each substitution in a cancer type were computed as the ratio (fold change) of the substitution frequency in the set of mutations inside versus the substitution frequency outside. Clustering of cancer types according to enrichments of 6-class based substitutions were computed using the hierarchical clustering function cluster.hierarchy from the Python scipy library^60^ with the linkage function “complete”.

### Mutational signatures extraction

*De novo* trinucleotide based SBS mutational signatures (96-mutation types using pyrimidines as reference) were extracted using SigProfiler framework^59, 61, 62^ for the 31 cancer types bearing at least 30 samples and 100,000 total SNVs (Supplementary Note 2). Input GRCh38 96-mutational catalogues were calculated using SigProfilerMatrixGenerator^59^ version 1.1.26 and mutational signatures were extracted with SigProfilerExtractor^61^ version 1.1.0 (Supplementary Note 2). *De novo* signatures were decomposed into COSMIC v3.2 GRCh38 reference signatures to allow comparisons across cancer types (Supplementary Note 2). All SBS signature names used in the manuscript correspond to this reference set. Signatures that were present in at least 5% of mutations in a sample were considered active in the sample. At the cancer type level, signatures that were active in at least 5% of samples were considered active in the cancer type.

### Assignment of mutational signatures to mutations and hotspots

The probability of each SNV ––considering its sample of origin and trinucleotide context–– to arise from each of the decomposed COSMIC signatures in the cancer type was obtained from SigProfilerExtractor (“Decomposed_Mutation_Probabilities.txt” table for the best extracted solution). As a result, a vector of mutational probabilities was generated for each SNV. In those cases where the 1 to 1 attribution of mutations to signatures was required, mutations were credited to the signature showing highest mutational probability (maximum likelihood^32^). Hotspots were assigned to mutational signatures by computing, first, the average mutational probability vector among the mutations contributing to the hotspot, and then selecting the signature with the maximum average probability (check Supplementary Note 3 for further details).

### Estimation of hotspot propensity

We set to estimate the propensity of commonly active mutational signatures to form hotspots across cancer types independently of their number of exposed samples and mutation burden contributed to each of them. First, we selected the 7 cancer types with the largest sample size (5,000 or more observed hotspots and prioritising non-meta-cancer types when possible), including: bladder-urinary tract cancers (BLADDER_URI), breast cancers (BRCA), colorectal cancers (COADREAD), oesophagus-stomach cancers (ESOPHA_STOMACH), non-small cell lung cancers (NSCLC), prostate cancers (PROSTATE) and skin melanomas (SKCM). Then, we selected the 14 signatures that fulfilled the following criteria: i) they showed 450 or more attributed hotspots (resulting in at least 1% of the total hotspots in the cancer type) and ii) contributed more than 300 high confidence mutations per sample attributed to the signature via highest probability (maximum likelihood p > 0.5) in at least 101 samples within the cancer type. The signature-cancer type pairs that passed these thresholds included: SBS1 (BLADDER_URI, BRCA, COADREAD, ESOPHA_STOMACH, NSCLC, PROSTATE), SBS13 (BLADDER_URI, BRCA, NSCLC), SBS17a (COADREAD, ESOPHA_STOMACH), SBS17b (COADREAD, ESOPHA_STOMACH), SBS18 (COADREAD), SBS2 (BLADDER_URI, BRCA, NSCLC), SBS3 (BRCA), SBS4 (NSCLC), SBS40 (COADREAD, NSCLC), SBS5 (BLADDER_URI, BRCA, COADREAD, ESOPHA_STOMACH, NSCLC, PROSTATE), SBS7a (SKCM), SBS7b (SKCM), SBS8 (PROSTATE), SBS93 (COADREAD). To estimate the propensity of a signature to form hotspots in a cancer type while accounting for differences in sample size and activity, we subsampled groups of 100 tumours of a given cancer type with a fixed number *n* (between 100 and 300 mutations/sample or ∼ 0.05 and 0.15 mutations/sample·Mbp), of randomly selected high confidence mutations (maximum likelihood p > 0.5) without replacement contributed by the signature under analysis. For each of the 1,000 subsamples, we then counted the number of observed hotspots (allowing a single hotspot per position) among the 100 *· n* subsampled mutations of the signature under analysis across cancer types. This analysis was carried out over the set of high mappable megabase bins (n = 2,196 bins; 2,012,091,302 bp). To estimate hotspot propensity at a larger mutation rate, we subsampled 60,000 total mutations (100 samples and 600 mutations/sample or 0.3 mutations/sample·Mbp) for those signatures where at least 101 samples had 600 high confidence mutations in the cancer type. Due to the limited sample size, the following signature and cancer type pairs were not included in this analysis: SBS1-BLADDER_URI, SBS1-BRCA, SBS1-NSCLC, SBS5-BLADDER_URI, and SBS17a-COADREAD.

### Signatures enrichment in hotspots

For each sample containing hotspots, we first computed the frequency of its active signatures in the sets of hotspot and non-hotspot mutations. These frequencies were obtained by aggregating the signature mutational probability vectors (conveying the relative contribution of all possible signatures to a mutation) across all mutations in the specified set, which were subsequently normalised to 1 (see Assignment of mutational signatures to mutations and hotspots). Then, a signature fold change (*FC*) in a given sample was computed as the ratio of the normalised frequency of the signature *S* inside hotspots versus the normalised frequency of the signature outside hotspots:

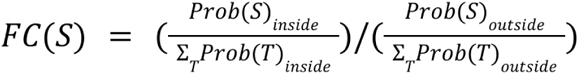

To obtain the fold change per active signature we calculated the median fold change among active samples. For each signature, we tested whether the magnitude of the frequency inside hotspots deferred from that of non-hotspot mutations across sample-paired observations. We applied a two-sided Wilcoxon rank-sum test using the “wilcoxon” function from the Python module scipy.stats^60^. The obtained p-values were adjusted for multiple testing using the Benjamini-Hochberg method from statsmodels.sandbox.stats.multicomp function^58^ in Python with *α* =0.01.

### Entropy of mutational signatures profiles

The entropy of the 96-channel profiles of SBS mutational signatures (COSMIC v3.2 GRCh38) was calculated after correcting by the trinucleotide content of the mappable genome, i.e. we only account for the relative mutability of each context. This is done by taking the frequency profiles from COSMIC, dividing each trinucleotide frequency by the genome-wide abundance of the corresponding reference triplet and normalising the resulting profile so that the probabilities add up to 1 (Supplementary table S5). The entropy, *H*, of a mutational signature profile given by a vector of frequencies (*p_k_* | *k*=1, …, 96) was calculated using the scipy.stats.entropy function^60^ in Python as follows:

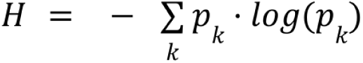

### Theoretical models of hotspots formation

We devised a method to compute the expected number of hotspots generated i) by a given mutational process and ii) in a given DNA region. The method assumes a simplifying theoretical scenario whereby i) all samples in the cohort have identical mutation rates and ii) positions mutate independently from one other. Briefly, if the regional mutation rate is homogeneous across the genome, we can estimate the probability that each position in the region undergoes each possible type of mutation (alternate allele) consistently with the relative mutability of each trinucleotide context dictated by the mutational process. The expected presence of a hotspot at a given position can then be calculated as the probability of at least two samples getting the same mutation at a given position. Because of statistical independence across positions, the regional expectation can be computed as the sum of expectations across positions.

Equipped with this method, we provide expected hotspot rate estimates associated with the 14 different mutational signatures from which we previously calculated hotspot propensity. Expected hotspot rates can be computed in two different scenarios: i) genome wide: assuming that mutability is homogeneous along the genome, hence this is based on the trinucleotide composition of the genome alone, and ii) per chunk: assuming cancer-type-specific variation in the distribution of mutation rate per signature asserted across genomic chunks. The analysis per chunks was conducted for different partitions of the mappable genome, using the following chunk sizes: 500, 250, 100, 50, 25, 10 Kbps and 1Mbp (Supplementary Note 5). The relative mutation rates per chunk, signature and cancer type were calculated from the total number of mutations attributed to the signature in the cancer type cohort estimated via maximum likelihood, i.e., mapping mutations to the signature with highest probability to have generated the mutation in the corresponding sample (Supplementary Note 3).

The same signature-cancer type pairs used to calculate observed hotspot propensity were analysed. For both models, comparisons between observed and expected hotspot propensity were carried out using 30,000 total mutations (300 muts/sample across mappable 1 Mbp bins), which was equivalent to approximately 0.15 muts/Mbp. To match the definition of hotspots with that of the expected calculation, only 1 observed hotspot per genomic position was considered (e.g., 2 C>A mutations and 2 C>T within the same position resulted in 1 hotspot).

We also implemented genome-wide and per-chunk theoretical hotspot rate models that used non-standard profiles to represent mutational processes. For the SBS1-induced hotspot analysis we implemented models that account for the differential SBS1 mutability across NpCpG>T contexts with and without accounting for the mutation rate bias due to methylated cytosines. For the models using methylated and unmethylated cytosine channels, methylated-unmethylated mutation rate fold-change was inferred empirically for each of the cancer types: COADREAD, ESOPHA_STOMACH, NSCLC (see section “Analysis of methylated CpG sites” in Methods and Supplementary Note 5).

### Analysis of large scale chromatin features

Chromatin accessibility and gene expression data for different tissues and cell lines matching the cancer types under analysis were obtained from the Epigenome Roadmap Project^63^ at egg2.wustl.edu/roadmap/web_portal (see Supplementary Table 6 for complete details and urls). Replication timing data for 7 cell lines from solid tissues was obtained from ENCODE^64^ at hgdownload.cse.ucsc.edu/goldenPath/hg19/encodeDCC/wgEncodeUwRepliSeq. For each of these three chromatin features (mapped to hg19 reference genome), we obtained the average signal across hg19 mappable megabases ––liftovered from the hg38 mappable megabase coordinates–– as follows. For chromatin accessibility, we first computed the average counts across megabases from genome-wide fold-enrichment DNase counts tracks (BIGWIG format) per epigenome. Then, for each megabase, we computed the average DNase-seq signal across the different epigenomes linked to a cancer type. Similarly, megabase gene expression signals were computed from normalised coverage genome tracks (BIGWIG format). In this case, if stranded libraries were available for an epigenome, we first added up the absolute RNA-seq signals from the negative and positive strands and then computed the average signal per epigenome and megabase. The cancer type signal per megabase was obtained by calculating the mean megabase RNA-seq signal from the different epigenomes linked to the cancer type. For replication timing data, we used the percentage-normalised Repli-seq signal tracks (BIGWIG format). Following the same approach, we obtained the average Repli-seq signal across cell lines per megabase. Signals extracted from BIGWIG files were handled using the Python package pyBigWig^65^.

In order to investigate the relationship between the number of hotspots and non-hotspot mutations with chromatin accessibility, gene expression and replication timing per signature and cancer type, we first intersected mutations inside and outside hotspots with mappable megabase bins. Next, we added up the vector of mutational probabilities of each mutation to arise from a signature in the cancer type (see Assignment of mutational signatures to mutations and hotspots). For each signature, we normalised its signal across megabases for mutations inside and outside hotspots. We then categorised mappable megabases into 10 percentiles (deciles) according to the distribution of the chromatin feature signal across megabases and plotted the signature activity inside and outside hotspots on each decile.

### Overdispersion of mutational signatures across the genome

For each signature, we fitted the distribution of their attributed mutation counts across mappable 1 Mbp bins by fitting a negative binomial regression model, which yields an overdispersion parameter. Briefly, the overdispersion parameter, referred to as *α* throughout, measures the excess variance over the mean, i.e., excess variance over the variance that we would expect if the mutation counts were Poisson distributed. Specifically, if *μ* denotes the mean, in our negative binomial regression setting the relationship between the variance *v* and the mean *μ* is given by the equation:

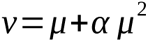

The Python function statsmodels.discrete.discrete_model.NegativeBinomial was used^58^.

### Analysis of CTCF binding sites

CTCF binding sites were defined as CTCF ChIP peaks in a tissue or cell line matching the cancer type. hg38 ChIP peak coordinates were downloaded from ReMap2022^66^ at remap.univ-amu.fr on 08-03-2022 (Supplementary Table 6). Only CTCF peaks within autosomes, 200-600 bp long, and showing 90% or more overlap to the mappable genome were kept for analysis (sigmoid colon n = 27,626; epithelial esophagus n = 38,740). Intersections between annotations were carried out using pybedtools^56, 57^. The enrichment of hotspots within CTCF binding sites was computed as follows. For each individual CTCF feature, we first constructed a window of length *L* where the feature of length *L_feature_*was positioned in the centre surrounded by two flanking sites of equal size (*L−L_feature_*) /2. Window length *L* was defined as 2,000 bp and feature length *L_feature_* as 600 bp (the maximum size encompassing all individual features under analysis). For each cancer type, we intersected each CTCF window with inside and outside hotspot mutations attributed to SBS17a and b by maximum likelihood. We obtained the expected distribution of mutations per signature by randomising 1,000 times the observed number of mutations inside hotspots across the window according to the signature trinucleotide probabilities and the window sequence composition. To compute the hotspot enrichment in CTCF binding sites (fold change), we piled-up mutations in equal positions across windows and calculated the ratio of mutations inside the feature versus its flanks. The significance of the observed fold changes was estimated by fitting simulated fold changes to a gaussian kernel density estimate distribution and deriving the upper quantile of the observed fold change. Resulting p-values were adjusted for multiple testing using the statsmodels Benjamini-Hochberg function^58^ with α=0.01. To visualise the results across piled-up CTCF binding sites, observed and expected mutation counts per position were normalised to the respective total number of mutations in the set across the window. A smoothing Savitzky-Golay filter of length 101 bp was applied using the scipy.signal.savgol_filter function^60^.

### Analysis of methylated CpG sites

Fractional methylation from whole genome bisulfite sequencing (WGBS) data for colon, esophagus/stomach and lung tissues was downloaded from Roadmap^63^ on 12-11-2022 (Supplementary Table 6). These tissues were selected based on the availability of uniformly processed WGBS, the gold-standard assay to quantify genome-wide methylation, within the Roadmap project. Methylation calls per CpG site were liftovered to hg38 and those failing to liftover or falling in a non-CpG site were removed. Only CpGs within autosomes and overlapping mappable 1 Mbp bins were kept. For COADREAD, ESOPHA_STOMACH and NSCLC cancer types, we annotated each mutation and hotspot with the average fractional methylation across the epigenomes matching the tissue of origin of the cancer type. Those CpG sites with missing methylation data in one or more epigenomes were skipped from analyses. CpGs with average fractional methylation above 0.5 were classified as methylated.

### Estimation of hotspot propensity in normal and germline tissues

We obtained somatic single base substitutions from different datasets to estimate SBS1 and SBS17a and b hotspot propensity in non-cancerous tissues. Somatic mutations from normal colonic crypts from human and mouse^36^ were downloaded from github.com/baezortega/CrossSpecies2021 and preprocessed as stated by the authors (steps 0 and 1 from the code repository). Only samples passing the quality criteria were used, resulting in 28 and 43 human and mouse samples, respectively. Somatic mutations from Barrett’s esophagus (BE) were obtained from ^43^. This dataset contains samples from 80 BE patients collected at different time points. In order to analyse a single sample per patient, we randomly selected 1 sample from the first sampling time point (T1) in each patient, resulting in 80 BE samples. Germline *de novo* mutations were obtained from 7 different datasets^37–42^, comprising a total of 7,796 families. Of note, *de novo* mutations in these datasets were not mapped to individuals/samples of origin. All datasets containing human mutations mapped to hg19 were liftovered to the hg38 reference genome. Mutations failing to map to hg38 or being mapped to a different reference nucleotide in hg38 were discarded. Mutations in sexual chromosomes were removed in all datasets. Finally, we filtered all human datasets with our mappable genome 1 Mbp bins. After filtering, we obtained a total of 52,233, 37,691 and 911,922 somatic mutations from human colonic crypts, mouse colonic crypts, and BE samples, respectively. Filtering of *de novo* germline mutations resulted in a total of 581,978 mutations for analysis.

We identified the signatures active in human and mouse colonic crypts, as well as in the germline, by signature fitting. Unlike the *de novo* signature deconstruction methods where the mutational signatures are inferred from scratch, signature fitting assumes that a collection of mutational signatures is given to then infer an exposure per signature and sample. We implemented signature fitting as a non-negative least squares regression, whereby a linear model is fit with the least squares criterion and the estimated model parameters (exposures per signature per sample) are constrained to be non-negative. Concretely, if *S* is a 96 *× k* matrix with the *k* given signature profiles arranged as columns, and *M* is a 96 *× n* matrix of mutation counts from *n* samples, signature fitting solves for:

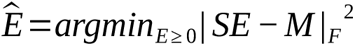

For human and mouse colonic crypts, we based our fitting on the three mutational signatures most active across the original study^36^, SBS1, SBS5 and SBS18. COSMIC v3.2 hg38 and mm10 reference signatures were used in human and mouse, respectively. In the case of the germline datasets, we tested the 92,170 possible combinations of 1-4 signatures from selected COSMIC signatures (SBS1, SBS2, SBS3, SBS4, SBS5, SBS6, SBS7a, SBS7b, SBS7c, SBS7d, SBS8, SBS9, SBS10a, SBS10b, SBS10c, SBS10d, SBS11, SBS12, SBS13, SBS14, SBS15, SBS16, SBS17a, SBS17b, SBS18, SBS19, SBS20, SBS21, SBS22, SBS23, SBS24, SBS25, SBS26, SBS27, SBS28, SBS29, SBS30, SBS40, SBS44). We conducted signature fitting with each combination and we thereupon computed the Akaike Information Criterion (AIC) of the resulting models. We selected the solution containing the signatures that were most frequently found across the top best solutions (SBS1 and SBS5). In Barrett’s esophagus samples, we used the mutational probabilities of individual mutations to arise from the different signatures active in the dataset as provided by the authors^43^. We focused our analysis on the top 5 active signatures: SBS1, SBS5, SBS17b, SBS17a, and SBS18.

Finally, we estimated hotspot propensity in each dataset by randomly sampling a number of mutations and/or samples for 1,000 times. A total of 5,000 mutations per signature (25 samples and 250 mutations/sample) and 2,500 mutations per signature (25 samples and 100 mutations/sample) were used in the human and mouse colonic crypts, respectively. For Barrett’s esophagus samples, we leveraged 15,000 mutations per signature (50 samples and 300 mutations/sample). To calculate germline hotspot propensity, where *de novo* mutations could not be mapped to individuals/samples of origin, we merged *de novo* mutations in the 7 datasets and randomly sampled 30,000 mutations.

### Sequence logos

We computed the frequency of the nucleotide sequence composition around hotspots attributed to a signature across cancer types. The same signature-cancer type groups as those used to calculate hotspot propensity were used. For each hotspot in the signature, we retrieved the 10 bp 5’ and 3’ flanking sequences (considering the strand containing a pyrimidine in the hotspot position) from bgreference package and built a 21 bp window centred at the hotspot. We then computed the information content over the nucleotide frequency with respect to the nucleotide hg38 mappable genome frequency at each position across the window. Logo plots and information content were generated using the Python package logomaker^67^ version 0.8.

### Additional software used

The following Python packages were used across different analyses, including: matplotlib^68^, numpy^69^, and pandas^70^.

## Supplementary material

**Supplementary Table S1**. Sequencing cohorts used in the analysis.

**Supplementary Table S2**. Cancer types used in the analysis.

**Supplementary Table S3**. List of cancer driver genes.

**Supplementary Table S4**. Total observed hotspots across cancer types using filtered mutations.

**Supplementary Table S5**. Normalised mutational profiles of signatures used in the analysis.

**Supplementary Table S6**. Annotations of large and small scale chromatin covariates.

**Supplementary Table S7**. Overdispersion of somatic mutations across megabase bins.

**Supplementary Note 1**. HotspotFinder.

**Supplementary Note 2**. Mutational signatures extraction.

**Supplementary Note 3**. Assignment of mutational signatures to mutations and hotspots.

**Supplementary Note 4**. Comment on hotspot propensity.

**Supplementary Note 5**. Theoretical models of hotspot propensity.

## Data availability

Somatic mutations were retrieved from several sources as listed in Supplementary Table S1. Large and small scale features (DNase, Repli-seq, RNA-seq, CTCF ChIP, and fractional methylation) are listed in Supplementary Table S6. Tables containing the hotspots identified across the cancer types analysed in this study will be made available at a Zenodo repository upon publication.

## Code availability

The code used within this work is freely available from external sources (see Methods) or has been developed in-house. HotspotFinder algorithm is available for download at bitbucket.org/bbglab/hotspotfinder. The in-house code containing the analyses and figures will be available at the time of publication.

## Acknowledgements

The authors wish to acknowledge the contribution of patients, families and biomedical researchers who shared, processed and sequenced the data used within the study. The results published here are in part based on the data generated by the Pan-Cancer Analysis of Whole Genomes, St Jude Children’s Research Hospital, PedcBioPortal, Therapeutically Applicable Research to Generate Effective Treatments program. This publication and the underlying study have been made possible partly on the basis of the data that Hartwig Medical Foundation has made available to the study. We acknowledge the technical contributions of Iker Reyes-Salazar and Loris Mularoni to HotspotFinder algorithm and annotations. N.L.-B. acknowledges funding from the European Research Council (consolidator grant 682398) and ERDF/Spanish Ministry of Science, Innovation and Universities – Spanish State Research Agency/DamReMap Project (RTI2018-094095-B-I00) and Asociación Española Contra el Cáncer (AECC) (GC16173697BIGA). IRB Barcelona is a recipient of a Severo Ochoa Centre of Excellence Award from the Spanish Ministry of Economy and Competitiveness (MINECO; Government of Spain) and is supported by CERCA (Generalitat de Catalunya). C.A.-P. is supported by “la Caixa” Foundation (ID 100010434) fellowship (LCF/BQ/ES18/11670011).

## Author contributions

C.A.-P, A.G.-P. and N.L.-B. conceptualised the project. C.A.-P curated the sequencing data for the analysis and implemented HotspotFinder. F.M. implemented the theoretical models of hotspots formation and generated the expected hotspot counts. C.A.-P carried out all the remaining analyses. C.A.-P prepared the figures. All authors participated in the design of the analyses and figures and in the interpretation of the results. C.A.-P, A.G.-P. and F.M. drafted the manuscript. All authors reviewed and edited the final manuscript. A.G.-P. and N.L.-B. supervised the project.

## References

1. Polak, P. et al. Cell-of-origin chromatin organization shapes the mutational landscape of cancer. Nature 518, 360–364 (2015).

2. Stamatoyannopoulos, J. A. et al. Human mutation rate associated with DNA replication timing. Nat. Genet. 41, 393–395 (2009).

3. Schuster-Böckler, B. & Lehner, B. Chromatin organization is a major influence on regional mutation rates in human cancer cells. Nature 488, 504–507 (2012).

4. Lawrence, M. S. et al. Mutational heterogeneity in cancer and the search for new cancer-associated genes. Nature 499, 214–218 (2013).

5. Nik-Zainal, S. et al. Mutational processes molding the genomes of 21 breast cancers. Cell 149, 979– 993 (2012).

6. Alexandrov, L. B. et al. Signatures of mutational processes in human cancer. Nature 500, 415–421 (2013).

7. Alexandrov, L. B. et al. The repertoire of mutational signatures in human cancer. Nature 578, 94–101 (2020).

8. Martincorena, I. et al. Universal patterns of selection in cancer and somatic tissues. Cell 171, 1029–1041.e21 (2017).

9. Pich, O. et al. Somatic and Germline Mutation Periodicity Follow the Orientation of the DNA Minor Groove around Nucleosomes. Cell 175, 1074–1087.e18 (2018).

10. Sabarinathan, R., Mularoni, L., Deu-Pons, J., Gonzalez-Perez, A. & López-Bigas, N. Nucleotide excision repair is impaired by binding of transcription factors to DNA. Nature 532, 264–267 (2016).

11. Perera, D. et al. Differential DNA repair underlies mutation hotspots at active promoters in cancer genomes. Nature 532, 259–263 (2016).

12. Kaiser, V. B., Taylor, M. S. & Semple, C. A. Mutational Biases Drive Elevated Rates of Substitution at Regulatory Sites across Cancer Types. PLoS Genet. 12, e1006207 (2016).

13. Katainen, R. et al. CTCF/cohesin-binding sites are frequently mutated in cancer. Nat. Genet. 47, 818–821 (2015).

14. Guo, Y. A. et al. Mutation hotspots at CTCF binding sites coupled to chromosomal instability in gastrointestinal cancers. Nat. Commun. 9, 1520 (2018).

15. Zou, X. et al. Short inverted repeats contribute to localized mutability in human somatic cells. Nucleic Acids Res. 45, 11213–11221 (2017).

16. Georgakopoulos-Soares, I., Morganella, S., Jain, N., Hemberg, M. & Nik-Zainal, S. Noncanonical secondary structures arising from non-B DNA motifs are determinants of mutagenesis. Genome Res. 28, 1264–1271 (2018).

17. Fredriksson, N. J. et al. Recurrent promoter mutations in melanoma are defined by an extended context-specific mutational signature. PLoS Genet. 13, e1006773 (2017).

18. Elliott, K. et al. Elevated pyrimidine dimer formation at distinct genomic bases underlies promoter mutation hotspots in UV-exposed cancers. PLoS Genet. 14, e1007849 (2018).

19. Mao, P. et al. ETS transcription factors induce a unique UV damage signature that drives recurrent mutagenesis in melanoma. Nat. Commun. 9, 2626 (2018).

20. Hess, J. M. et al. Passenger hotspot mutations in cancer. Cancer Cell 36, 288–301.e14 (2019).

21. Buisson, R. et al. Passenger hotspot mutations in cancer driven by APOBEC3A and mesoscale genomic features. Science 364, (2019).

22. Shi, M.-J. et al. Identification of new driver and passenger mutations within APOBEC-induced hotspot mutations in bladder cancer. Genome Med. 12, 85 (2020).

23. Gonzalez-Perez, A., Sabarinathan, R. & Lopez-Bigas, N. Local determinants of the mutational landscape of the human genome. Cell 177, 101–114 (2019).

24. Supek, F. & Lehner, B. Scales and mechanisms of somatic mutation rate variation across the human genome. DNA Repair (Amst) 81, 102647 (2019).

25. Smith, T. C. A., Carr, A. M. & Eyre-Walker, A. C. Are sites with multiple single nucleotide variants in cancer genomes a consequence of drivers, hypermutable sites or sequencing errors? PeerJ 4, e2391 (2016).

26. Sondka, Z. et al. The COSMIC Cancer Gene Census: describing genetic dysfunction across all human cancers. Nat. Rev. Cancer 18, 696–705 (2018).

27. Martínez-Jiménez, F. et al. A compendium of mutational cancer driver genes. Nat. Rev. Cancer 20, 555–572 (2020).

28. Priestley, P. et al. Pan-cancer whole-genome analyses of metastatic solid tumours. Nature 575, 210– 216 (2019).

29. Secrier, M. et al. Mutational signatures in esophageal adenocarcinoma define etiologically distinct subgroups with therapeutic relevance. Nat. Genet. 48, 1131–1141 (2016).

30. Christensen, S. et al. 5-Fluorouracil treatment induces characteristic T>G mutations in human cancer. Nat. Commun. 10, 4571 (2019).

31. Polak, P. et al. Reduced local mutation density in regulatory DNA of cancer genomes is linked to DNA repair. Nat. Biotechnol. 32, 71–75 (2014).

32. Morganella, S. et al. The topography of mutational processes in breast cancer genomes. Nat. Commun. 7, 11383 (2016).

33. Tomkova, M., Tomek, J., Kriaucionis, S. & Schuster-Böckler, B. Mutational signature distribution varies with DNA replication timing and strand asymmetry. Genome Biol. 19, 129 (2018).

34. Otlu, B., et al. Topography of mutational signatures in human cancer. BioRxiv (2022) doi:10.1101/2022.05.29.493921.

35. Moore, L. et al. The mutational landscape of human somatic and germline cells. Nature 597, 381– 386 (2021).

36. Cagan, A. et al. Somatic mutation rates scale with lifespan across mammals. Nature 604, 517–524 (2022).

37. Rahbari, R. et al. Timing, rates and spectra of human germline mutation. Nat. Genet. 48, 126–133 (2016).

38. An, J.-Y. et al. Genome-wide de novo risk score implicates promoter variation in autism spectrum disorder. Science 362, (2018).

39. Halldorsson, B. V. et al. Characterizing mutagenic effects of recombination through a sequence-level genetic map. Science 363, (2019).

40. CYuen, R. K., et al. Whole genome sequencing resource identifies 18 new candidate genes for autism spectrum disorder. Nat. Neurosci. 20, 602–611 (2017).

41. Sasani, T. A. et al. Large, three-generation human families reveal post-zygotic mosaicism and variability in germline mutation accumulation. eLife 8, (2019).

42. Goldmann, J. M. et al. Parent-of-origin-specific signatures of de novo mutations. Nat. Genet. 48, 935–939 (2016).

43. Paulson, T. G. et al. Somatic whole genome dynamics of precancer in Barrett’s esophagus reveals features associated with disease progression. Nat. Commun. 13, 2300 (2022).

44. Pich, O. et al. The mutational footprints of cancer therapies. Nat. Genet. 51, 1732–1740 (2019).

45. Long, H. et al. Evolutionary determinants of genome-wide nucleotide composition. *Nat*. Ecol. Evol. 2, 237–240 (2018).

46. Barbour, J. A., et al. XPD protects CTCF-Cohesin binding sites from somatic mutagenesis. BioRxiv (2022) doi:10.1101/2022.01.21.477237.

47. Fryxell, K. J. & Zuckerkandl, E. Cytosine deamination plays a primary role in the evolution of mammalian isochores. Mol. Biol. Evol. 17, 1371–1383 (2000).

48. Stobbe, M. D. et al. Recurrent somatic mutations reveal new insights into consequences of mutagenic processes in cancer. PLoS Comput. Biol. 15, e1007496 (2019).

49. Rheinbay, E. et al. Analyses of non-coding somatic drivers in 2,658 cancer whole genomes. Nature 578, 102–111 (2020).

50. Sherman, M. A. et al. Genome-wide mapping of somatic mutation rates uncovers drivers of cancer. Nat. Biotechnol. 40, 1634–1643 (2022).

51. Muiños, F., Martínez-Jiménez, F., Pich, O., Gonzalez-Perez, A. & Lopez-Bigas, N. In silico saturation mutagenesis of cancer genes. Nature 596, 428–432 (2021).

52. Kundra, R. et al. Oncotree: A cancer classification system for precision oncology. *JCO Clin*. Cancer Inform. 5, 221–230 (2021).

53. Frankish, A. et al. GENCODE reference annotation for the human and mouse genomes. Nucleic Acids Res. 47, D766–D773 (2019).

54. Marco-Sola, S., Sammeth, M., Guigó, R. & Ribeca, P. The GEM mapper: fast, accurate and versatile alignment by filtration. Nat. Methods 9, 1185–1188 (2012).

55. Karczewski, K. J. et al. The mutational constraint spectrum quantified from variation in 141,456 humans. Nature 581, 434–443 (2020).

56. Quinlan, A. R. & Hall, I. M. BEDTools: a flexible suite of utilities for comparing genomic features. Bioinformatics 26, 841–842 (2010).

57. Dale, R. K., Pedersen, B. S. & Quinlan, A. R. Pybedtools: a flexible Python library for manipulating genomic datasets and annotations. Bioinformatics 27, 3423–3424 (2011).

58. Seabold, S. & Perktold, J. Statsmodels: Econometric and Statistical Modeling with Python. in Proceedings of the 9th Python in Science Conference 92–96 (SciPy, 2010). doi:10.25080/Majora-92bf1922-011.

59. Bergstrom, E. N. et al. SigProfilerMatrixGenerator: a tool for visualizing and exploring patterns of small mutational events. BMC Genomics 20, 685 (2019).

60. Virtanen, P. et al. SciPy 1.0: fundamental algorithms for scientific computing in Python. Nat. Methods 17, 261–272 (2020).

61. Islam, S. M. A. et al. Uncovering novel mutational signatures by de novo extraction with SigProfilerExtractor. Cell Genomics 2, None (2022).

62. Alexandrov, L. B., Nik-Zainal, S., Wedge, D. C., Campbell, P. J. & Stratton, M. R. Deciphering signatures of mutational processes operative in human cancer. Cell Rep. 3, 246–259 (2013).

63. Roadmap Epigenomics Consortium et al. Integrative analysis of 111 reference human epigenomes. Nature 518, 317–330 (2015).

64. ENCODE Project Consortium. An integrated encyclopedia of DNA elements in the human genome. Nature 489, 57–74 (2012).

65. Ryan, D., Grüning, B. & Ramirez, F. Pybigwig 0.2.4. Zenodo (2016) doi:10.5281/zenodo.45238.

66. Hammal, F., de Langen, P., Bergon, A., Lopez, F. & Ballester, B. ReMap 2022: a database of Human, Mouse, Drosophila and Arabidopsis regulatory regions from an integrative analysis of DNA-binding sequencing experiments. Nucleic Acids Res. 50, D316–D325 (2022).

67. Tareen, A. & Kinney, J. B. Logomaker: beautiful sequence logos in Python. Bioinformatics 36, 2272– 2274 (2020).

68. Hunter, J. D. Matplotlib: A 2D Graphics Environment. Comput. Sci. Eng. 9, 90–95 (2007).

69. Harris, C. R. et al. Array programming with NumPy. Nature 585, 357–362 (2020).

70. Reback, J. et al. pandas-dev/pandas: Pandas 1.4.4. Zenodo (2022) doi:10.5281/zenodo.7037953.

