## Supplementary Figures for "Hotspot propensity across mutational processes"

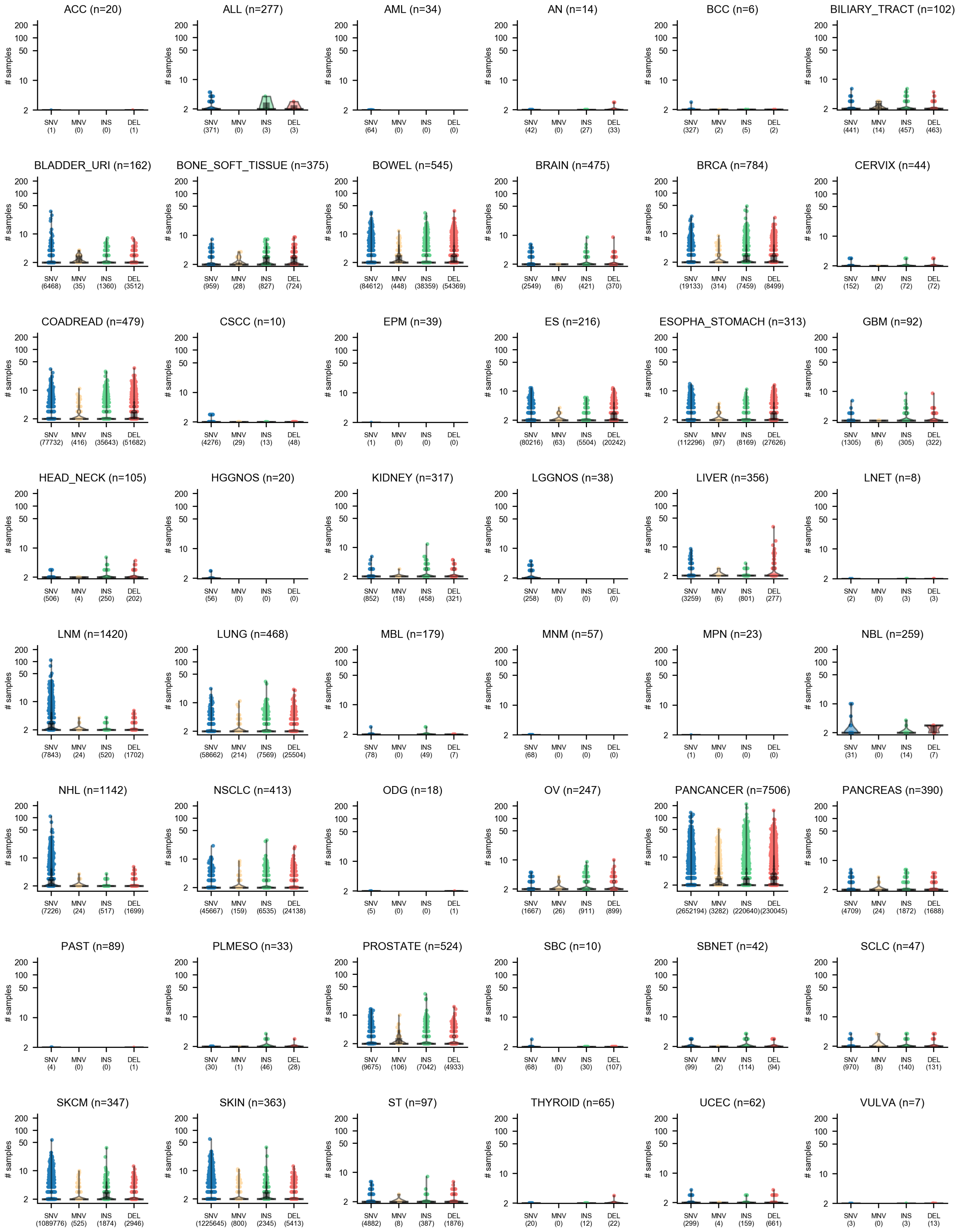

### samples
200 100 50 10 2
ACC (n=20) SNV (1) MNV (0) INS (0) DEL (1)
ALL (n=277) SNV (371) MNV (0) INS (3) DEL (3)
AML (n=34) SNV (64) MNV (0) INS (0) DEL (0)
AN (n=14) SNV (42) MNV (0) INS (27) DEL (33)
BCC (n=6) SNV (327) MNV (2) INS (5) DEL (2)
BILIARY_TRACT (n=102) SNV (441) MNV (14) INS (457) DEL (463)
BLADDER_URI (n=162) SNV (6468) MNV (35) INS (1360) DEL (3512)
BONE_SOFT_TISSUE (n=375) SNV (959) MNV (28) INS (827) DEL (724)
BOWEL (n=545) SNV (84612) MNV (448) INS (38359) DEL (54369)
BRAIN (n=475) SNV (2549) MNV (6) INS (421) DEL (370)
BRCA (n=784) SNV (19133) MNV (314) INS (7459) DEL (8499)
CERVIX (n=44) SNV (152) MNV (2) INS (72) DEL (72)
COADREAD (n=479) SNV (77732) MNV (416) INS (35643) DEL (51682)
CSCC (n=10) SNV (4276) MNV (29) INS (13) DEL (48)
EPM (n=39) SNV (1) MNV (0) INS (0) DEL (0)
ES (n=216) SNV (80216) MNV (63) INS (5504) DEL (20242)
ESOPHA_STOMACH (n=313) SNV (112296) MNV (97) INS (8169) DEL (27626)
GBM (n=92) SNV (1305) MNV (6) INS (305) DEL (322)
HEAD_NECK (n=105) SNV (506) MNV (4) INS (250) DEL (202)
HGGNOS (n=20) SNV (56) MNV (0) INS (0) DEL (0)
KIDNEY (n=317) SNV (852) MNV (18) INS (458) DEL (321)
LGGNOS (n=38) SNV (258) MNV (0) INS (0) DEL (0)
LIVER (n=356) SNV (3259) MNV (6) INS (801) DEL (277)
LNET (n=8) SNV (2) MNV (0) INS (3) DEL (3)
LNM (n=1420) SNV (7843) MNV (24) INS (520) DEL (1702)
LUNG (n=468) SNV (58662) MNV (214) INS (7569) DEL (25504)
MBL (n=179) SNV (78) MNV (0) INS (49) DEL (7)
MNM (n=57) SNV (68) MNV (0) INS (0) DEL (0)
MPN (n=23) SNV (1) MNV (0) INS (0) DEL (0)
NBL (n=259) SNV (31) MNV (0) INS (14) DEL (7)
NHL (n=1142) SNV (7226) MNV (24) INS (517) DEL (1699)
NSCLC (n=413) SNV (45667) MNV (159) INS (6535) DEL (24138)
ODG (n=18) SNV (5) MNV (0) INS (0) DEL (1)
OV (n=247) SNV (1667) MNV (26) INS (911) DEL (899)
PANCANCER (n=7506) SNV (2652194) MNV (3282) INS (220640) DEL (230045)
PANCREAS (n=390) SNV (4709) MNV (24) INS (1872) DEL (1688)
PAST (n=89) SNV (4) MNV (0) INS (0) DEL (1)
PLMESO (n=33) SNV (30) MNV (1) INS (46) DEL (28)
PROSTATE (n=524) SNV (9675) MNV (106) INS (7042) DEL (4933)
SBC (n=10) SNV (68) MNV (0) INS (30) DEL (107)
SBNET (n=42) SNV (99) MNV (2) INS (114) DEL (94)
SCLC (n=47) SNV (970) MNV (8) INS (140) DEL (131)
SKCM (n=347) SNV (1089776) MNV (525) INS (1874) DEL (2946)
SKIN (n=363) SNV (1225645) MNV (800) INS (2345) DEL (5413)
ST (n=97) SNV (4882) MNV (8) INS (387) DEL (1876)
THYROID (n=65) SNV (20) MNV (0) INS (12) DEL (22)
UCEC (n=62) SNV (299) MNV (4) INS (159) DEL (661)
VULVA (n=7) SNV (3) MNV (0) INS (3) DEL (13)

**Supp. Fig. 1. Hotspot size across cancer types.** Distribution of hotspot sizes across the four mutation types (SNVs, MNVs, insertions and deletions) per cancer type. Dots represent individual hotspots per mutation type (total number is shown in parenthesis). Horizontal black lines show median hotspot size.

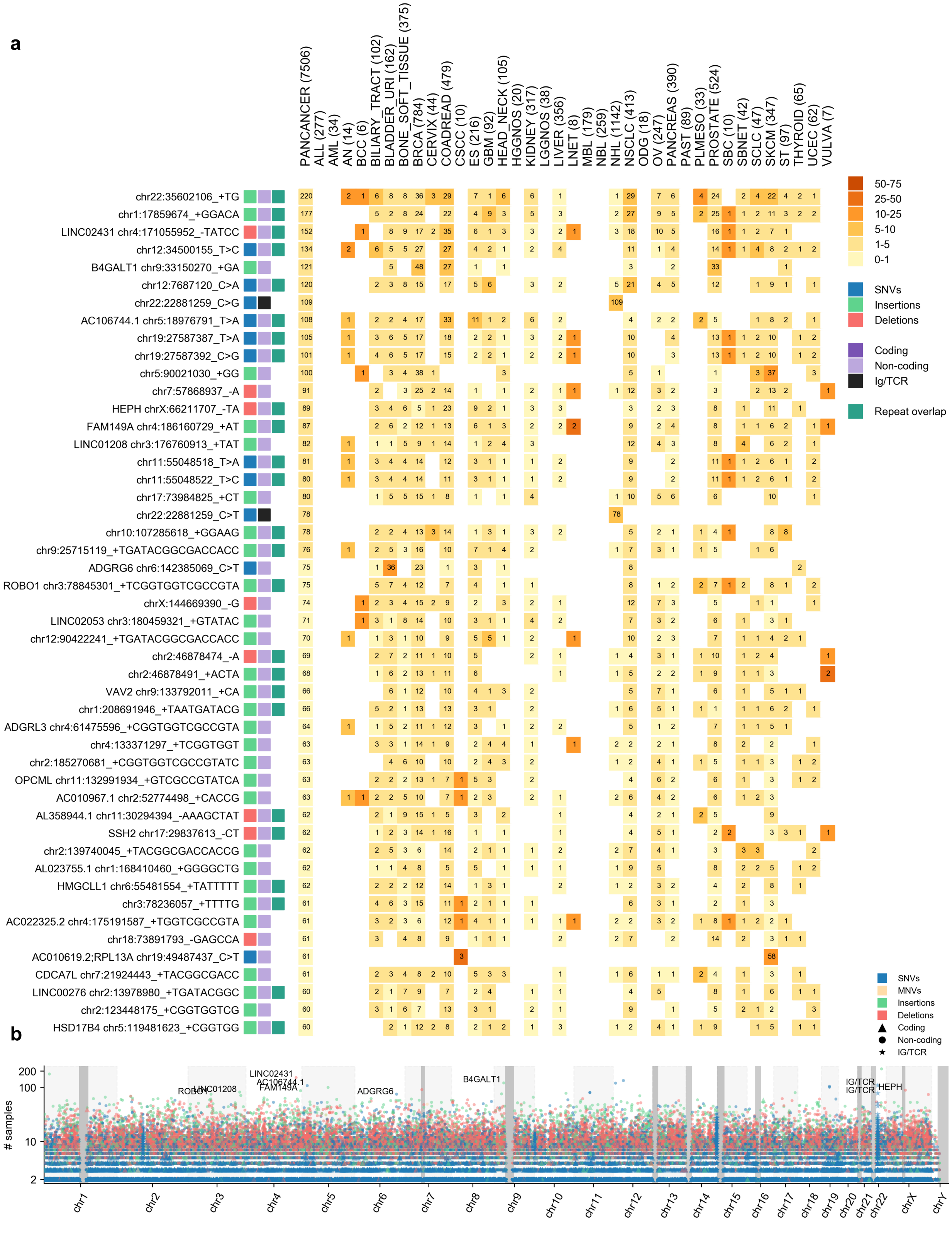
a
PANCANCER (7506)
ALL (277)
AML (34)
AN (14)
BCC (6)
BILIARY_TRACT (102)
BLADDER_URI (162)
BONE_SOFT_TISSUE (375)
BRCA (784)
CERVIX (44)
COADREAD (479)
CSCC (10)
ES (216)
GBM (92)
HEAD_NECK (105)
HGGNOS (20)
KIDNEY (317)
LGGNOS (38)
LIVER (356)
LNET (8)
MBL (179)
NBL (259)
NHL (1142)
NSCLC (413)
ODG (18)
OV (247)
PANCREAS (390)
PAST (89)
PLMESO (33)
PROSTATE (524)
SBC (10)
SBNET (42)
SCLC (47)
SKCM (347)
ST (97)
THYROID (65)
UCEC (62)
VULVA (7)
chr22:35602106_+TG
chr1:17859674_+GGACA
LINC02431 chr4:171055952_-TATCC
chr12:34500155_T>C
B4GALT1 chr9:33150270_+GA
chr12:7687120_C>A
chr22:22881259_C>G
AC106744.1 chr5:18976791_T>A
chr19:27587387_T>A
chr19:27587392_C>G
chr5:90021030_+GG
chr7:57868937_-A
HEPH chrX:66211707_-TA
FAM149A chr4:186160729_+AT
LINC01208 chr3:176760913_+TAT
chr11:55048518_T>A
chr11:55048522_T>C
chr17:73984825_+CT
chr22:22881259_C>T
chr10:107285618_+GGAAG
chr9:25715119_+TGATACGGCGACCACC
ADGRG6 chr6:142385069_C>T
ROBO1 chr3:78845301_+TCGGTGGTCGCCGTA
chrX:144669390_-G
LINC02053 chr3:180459321_+GTATAC
chr12:90422241_+TGATACGGCGACCACC
chr2:46878474_-A
chr2:46878491_+ACTA
VAV2 chr9:133792011_+CA
chr1:208691946_+TAATGATACG
ADGRL3 chr4:61475596_+CGGTGGTCGCCGTA
chr4:133371297_+TCGGTGGT
chr2:185270681_+CGGTGGTCGCCGTATC
OPCML chr11:132991934_+GTCGCCGTATCA
AC010967.1 chr2:52774498_+CACCG
AL358944.1 chr11:30294394_-AAAGCTAT
SSH2 chr17:29837613_-CT
chr2:139740045_+TACGGCGACCACCG
AL023755.1 chr1:168410460_+GGGGCTG
HMGCLL1 chr6:55481554_+TATTTTT
chr3:78236057_+TTTTG
AC022325.2 chr4:175191587_+TGGTCGCCGTA
chr18:73891793_-GAGCCA
AC010619.2;RPL13A chr19:49487437_C>T
CDCA7L chr7:21924443_+TACGGCGACC
LINC00276 chr2:13978980_+TGATACGGC
chr2:123448175_+CGGTGGTCG
HSD17B4 chr5:119481623_+CGGTGG
50-75
25-50
10-25
5-10
1-5
0-1
SNVs
Insertions
Deletions
Coding
Non-coding
Ig/TCR
Repeat overlap
b
### samples
200
100
10
2
ROBO1
LINC01208
LINC02431
AC106744.1
FAM149A
ADGRG6
B4GALT1
IG/TCR
IG/TCR
HEPH
chr1
chr2
chr3
chr4
chr5
chr6
chr7
chr8
chr9
chr10
chr11
chr12
chr13
chr14
chr15
chr16
chr17
chr18
chr19
chr20
chr21
chr22
chrX
chrY
SNVs
MNVs
Insertions
Deletions
Coding
Non-coding
IG/TCR

**Supp. Fig. 2. Top recurrent hotspots and their distribution across the genome. a**) Heatmap showing mutational frequencies of the top recurrent hotspots bearing 60 or more mutated samples. Rows show hotspots (and genomic elements overlapping them, if any) found by HotspotFinder. Hotspots are denoted by their genomic coordinates, and mutation alternate. Columns show cancer types (sample size in parenthesis). The number of mutated samples and the corresponding mutational frequency in the cancer type are shown in each square. Leftmost columns represent annotations of the mutation type, region annotation class, and overlap of a repetitive sequence of hotspots as identified by HotspotFinder. **b**) Distribution of hotspots for each mutation type across the mappable genome. Hotspot shape illustrates whether the hotspot overlaps coding, non-coding or immunoglobulin and T-cell receptor regions. Regions of 10 or more consecutive low mappable megabases are shown in grey.

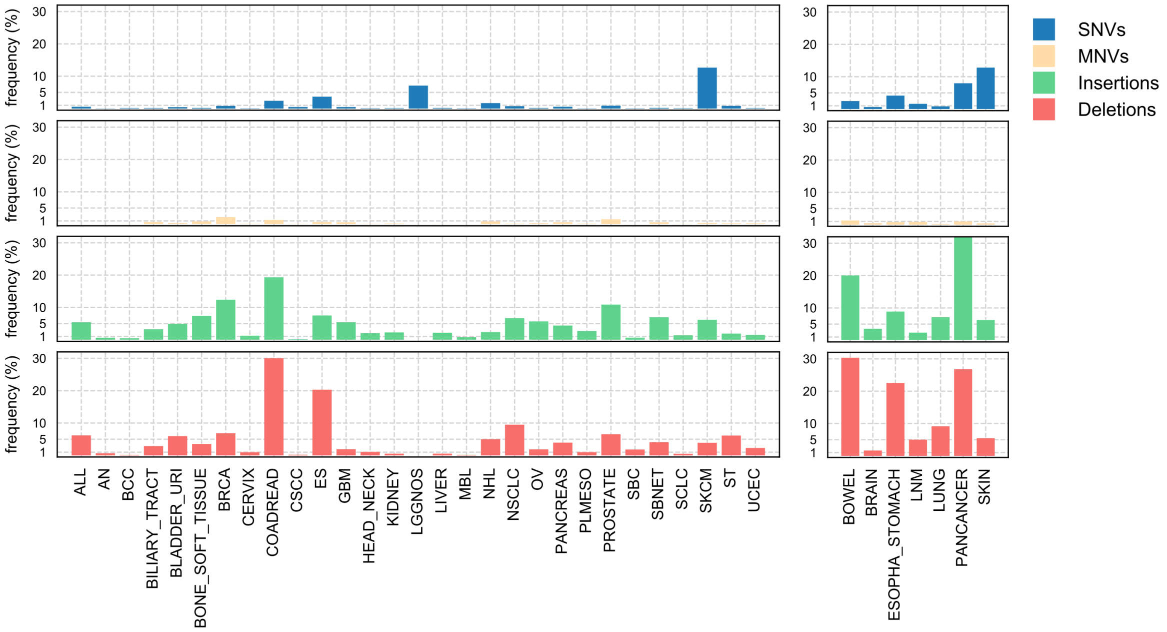


**Supp. Fig. 3. Frequency of hotspots formation across mutation types.** Bars showing the percentage of mutations that overlap a hotspot across cancers and mutation types.

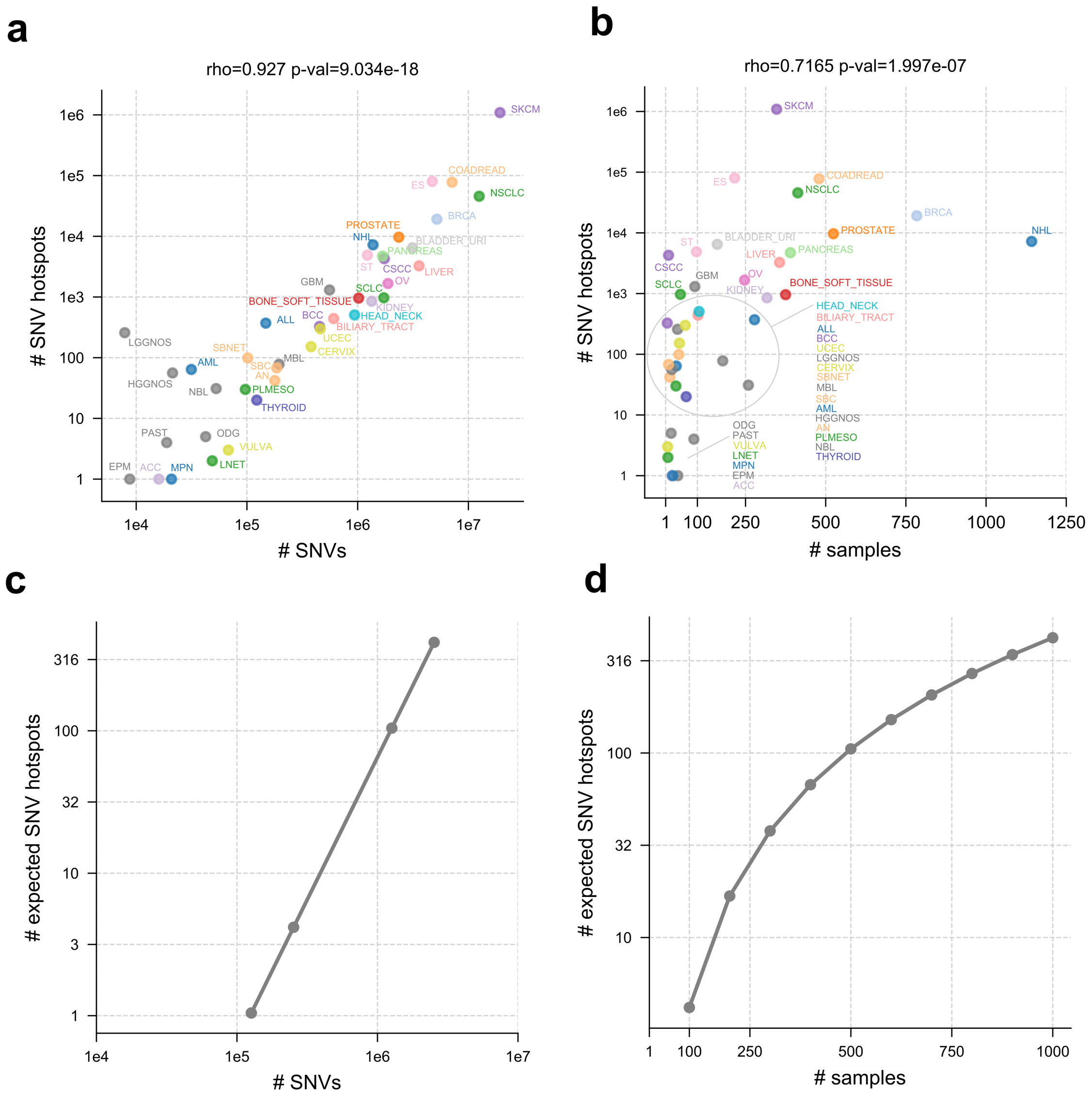


**Supp. Fig. 4. Relationship between the increase of hotspot burden, mutation burden and sample size.** **a**) Relationship between number of observed hotspots and mutations across cancer types. Spearman's rho correlation coefficient among both variables is shown on top. **b**) Correlation among observed SNV hotspots and sample size across cancer types. Spearman's rho correlation coefficient among both variables is shown on top. **c**) Expected number of hotspots generated under a theoretical model using 100 samples with equal mutation burden and homogeneous distribution of trinucleotide-specific mutation rates across the genome (Supplementary Note 5). **d**) Expected number of hotspots under the same theoretical model shown in c across different sample sizes with equal mutation burden (1 SNV / Mbp per sample).

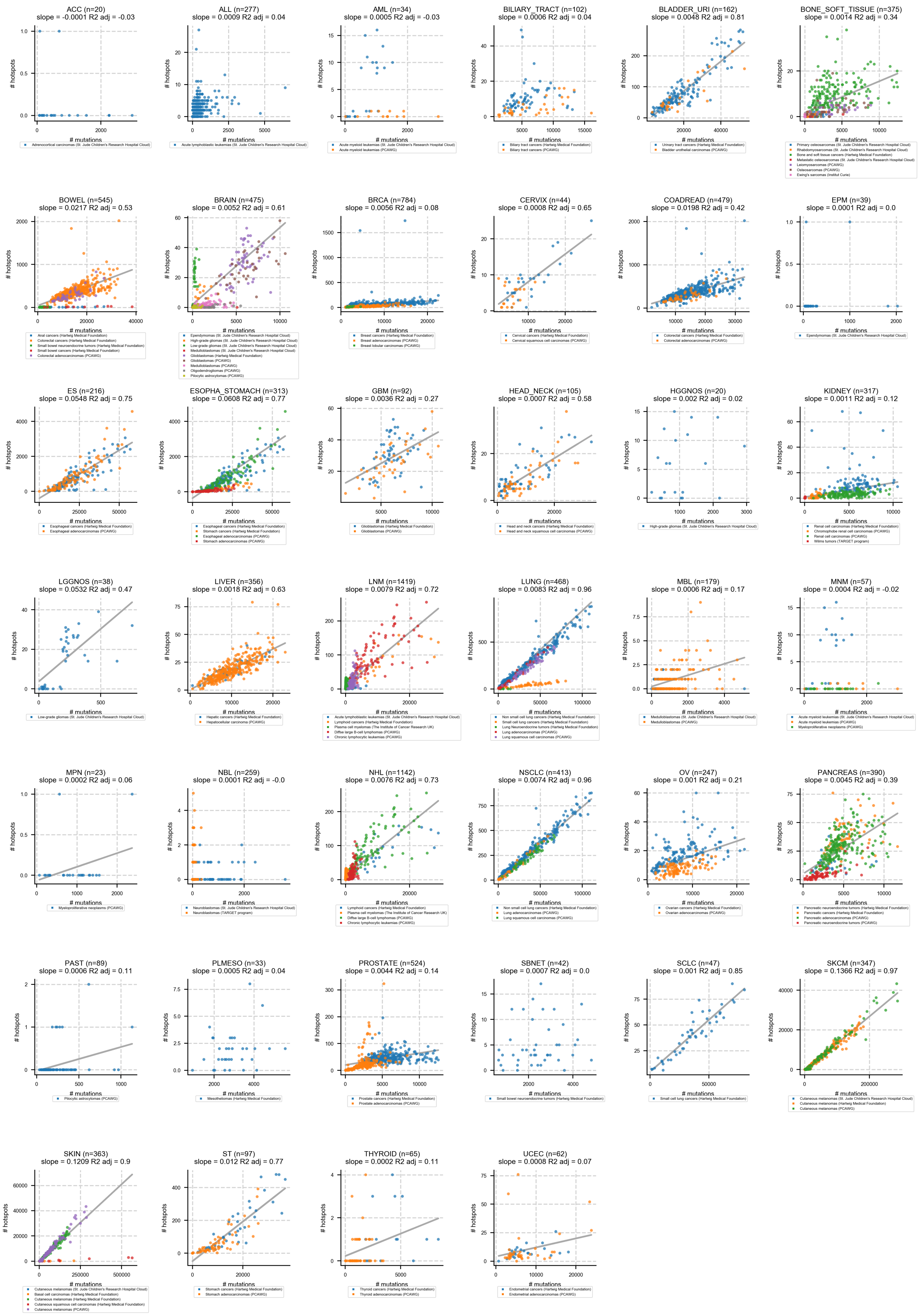
ACC (n=20)
slope = -0.0001 R2 adj = -0.03
ALL (n=277)
slope = 0.0009 R2 adj = 0.04
AML (n=34)
slope = 0.0005 R2 adj = -0.03
BILIARY_TRACT (n=102)
slope = 0.0006 R2 adj = 0.04
BLADDER_URI (n=162)
slope = 0.0048 R2 adj = 0.81
BONE_SOFT_TISSUE (n=375)
slope = 0.0014 R2 adj = 0.34
BOWEL (n=545)
slope = 0.0217 R2 adj = 0.53
BRAIN (n=475)
slope = 0.0052 R2 adj = 0.61
BRCA (n=784)
slope = 0.0056 R2 adj = 0.08
CERVIX (n=44)
slope = 0.0008 R2 adj = 0.65
COADREAD (n=479)
slope = 0.0198 R2 adj = 0.42
EPM (n=39)
slope = 0.0001 R2 adj = 0.0
ES (n=216)
slope = 0.0548 R2 adj = 0.75
ESOPHA_STOMACH (n=313)
slope = 0.0608 R2 adj = 0.77
GBM (n=92)
slope = 0.0036 R2 adj = 0.27
HEAD_NECK (n=105)
slope = 0.0007 R2 adj = 0.58
HGGNOS (n=20)
slope = 0.002 R2 adj = 0.02
KIDNEY (n=317)
slope = 0.0011 R2 adj = 0.12
LGGNOS (n=38)
slope = 0.0532 R2 adj = 0.47
LIVER (n=356)
slope = 0.0018 R2 adj = 0.63
LNM (n=1419)
slope = 0.0079 R2 adj = 0.72
LUNG (n=468)
slope = 0.0083 R2 adj = 0.96
MBL (n=179)
slope = 0.0006 R2 adj = 0.17
MNM (n=57)
slope = 0.0004 R2 adj = -0.02
MPN (n=23)
slope = 0.0002 R2 adj = 0.06
NBL (n=259)
slope = 0.0001 R2 adj = -0.0
NHL (n=1142)
slope = 0.0076 R2 adj = 0.73
NSCLC (n=413)
slope = 0.0074 R2 adj = 0.96
OV (n=247)
slope = 0.001 R2 adj = 0.21
PANCREAS (n=390)
slope = 0.0045 R2 adj = 0.39
PAST (n=89)
slope = 0.0006 R2 adj = 0.11
PLMESO (n=33)
slope = 0.0005 R2 adj = 0.04
PROSTATE (n=524)
slope = 0.0044 R2 adj = 0.14
SBNET (n=42)
slope = 0.0007 R2 adj = 0.0
SCLC (n=47)
slope = 0.001 R2 adj = 0.85
SKCM (n=347)
slope = 0.1366 R2 adj = 0.97
SKIN (n=363)
slope = 0.1209 R2 adj = 0.9
ST (n=97)
slope = 0.012 R2 adj = 0.77
THYROID (n=65)
slope = 0.0002 R2 adj = 0.11
UCEC (n=62)
slope = 0.0008 R2 adj = 0.07
### hotspots
### mutations

**Supp. Fig. 5. Relationship between observed hotspots and mutations.** Scatter plots showing number of SNV hotspots and mutations per sample across cancer types. Only datasets containing SNV hotspots and 20 or more samples are plotted. Slope and adjusted $R^2$ from linear regression models are shown on top of each plot. Regression lines are depicted for cancer types with model adjusted $R^2$ greater than 0.05. Samples colouring follows the sequencing cohort of origin, shown on the legend below.

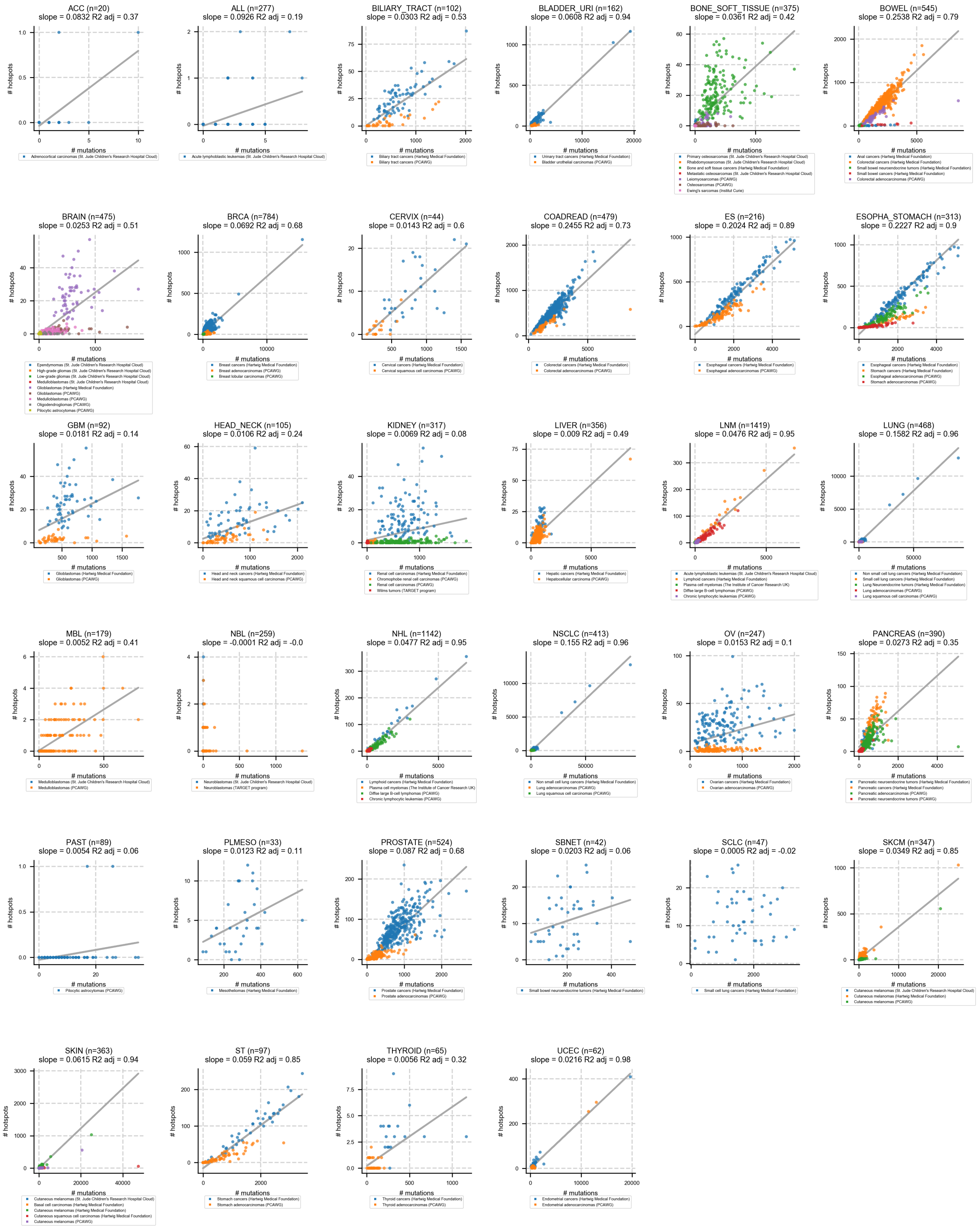
ACC (n=20)
slope = 0.0832 R2 adj = 0.37
ALL (n=277)
slope = 0.0926 R2 adj = 0.19
BILIARY_TRACT (n=102)
slope = 0.0303 R2 adj = 0.53
BLADDER_URI (n=162)
slope = 0.0608 R2 adj = 0.94
BONE_SOFT_TISSUE (n=375)
slope = 0.0361 R2 adj = 0.42
BOWEL (n=545)
slope = 0.2538 R2 adj = 0.79
BRAIN (n=475)
slope = 0.0253 R2 adj = 0.51
BRCA (n=784)
slope = 0.0692 R2 adj = 0.68
CERVIX (n=44)
slope = 0.0143 R2 adj = 0.6
COADREAD (n=479)
slope = 0.2455 R2 adj = 0.73
ES (n=216)
slope = 0.2024 R2 adj = 0.89
ESOPHA_STOMACH (n=313)
slope = 0.2227 R2 adj = 0.9
GBM (n=92)
slope = 0.0181 R2 adj = 0.14
HEAD_NECK (n=105)
slope = 0.0106 R2 adj = 0.24
KIDNEY (n=317)
slope = 0.0069 R2 adj = 0.08
LIVER (n=356)
slope = 0.009 R2 adj = 0.49
LNM (n=1419)
slope = 0.0476 R2 adj = 0.95
LUNG (n=468)
slope = 0.1582 R2 adj = 0.96
MBL (n=179)
slope = 0.0052 R2 adj = 0.41
NBL (n=259)
slope = -0.0001 R2 adj = -0.0
NHL (n=1142)
slope = 0.0477 R2 adj = 0.95
NSCLC (n=413)
slope = 0.155 R2 adj = 0.96
OV (n=247)
slope = 0.0153 R2 adj = 0.1
PANCREAS (n=390)
slope = 0.0273 R2 adj = 0.35
PAST (n=89)
slope = 0.0054 R2 adj = 0.06
PLMESO (n=33)
slope = 0.0123 R2 adj = 0.11
PROSTATE (n=524)
slope = 0.087 R2 adj = 0.68
SBNET (n=42)
slope = 0.0203 R2 adj = 0.06
SCLC (n=47)
slope = 0.0005 R2 adj = -0.02
SKCM (n=347)
slope = 0.0349 R2 adj = 0.85
SKIN (n=363)
slope = 0.0615 R2 adj = 0.94
ST (n=97)
slope = 0.059 R2 adj = 0.85
THYROID (n=65)
slope = 0.0056 R2 adj = 0.32
UCEC (n=62)
slope = 0.0216 R2 adj = 0.98
### hotspots
### mutations

**Supp. Fig. 6. Relationship between observed indel hotspots and mutations.** Scatter plots showing number of indel (merge of insertion and deletion) hotspots and mutations per sample across cancer types. Only datasets containing indel hotspots and 20 or more samples are plotted. Slope and adjusted $R^2$ from linear regression models are shown on top of each plot. Regression lines are depicted for cancer types with model adjusted $R^2$ greater than 0.05. Samples colouring follows the sequencing cohort of origin, shown on the legend below.

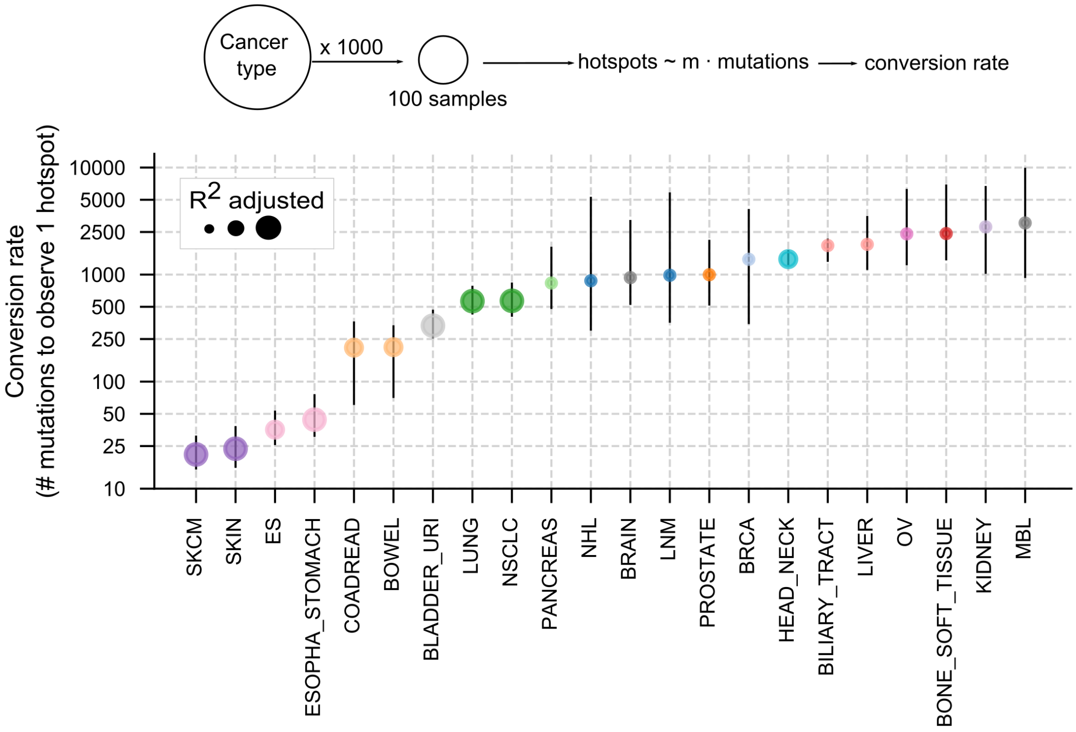


**Supp. Fig. 7. Conversion rates.** Median conversion rates (number of mutations to observe 1 hotspot) across cancer types with more than 100 samples and at least 750 significant linear models across random replicates (mean=978.9 models per cancer type, range=777-1000; Methods). Dot size represents the goodness of fit of the linear model (R2 adjusted). Error bars show the range of conversion rates across significant replicates.

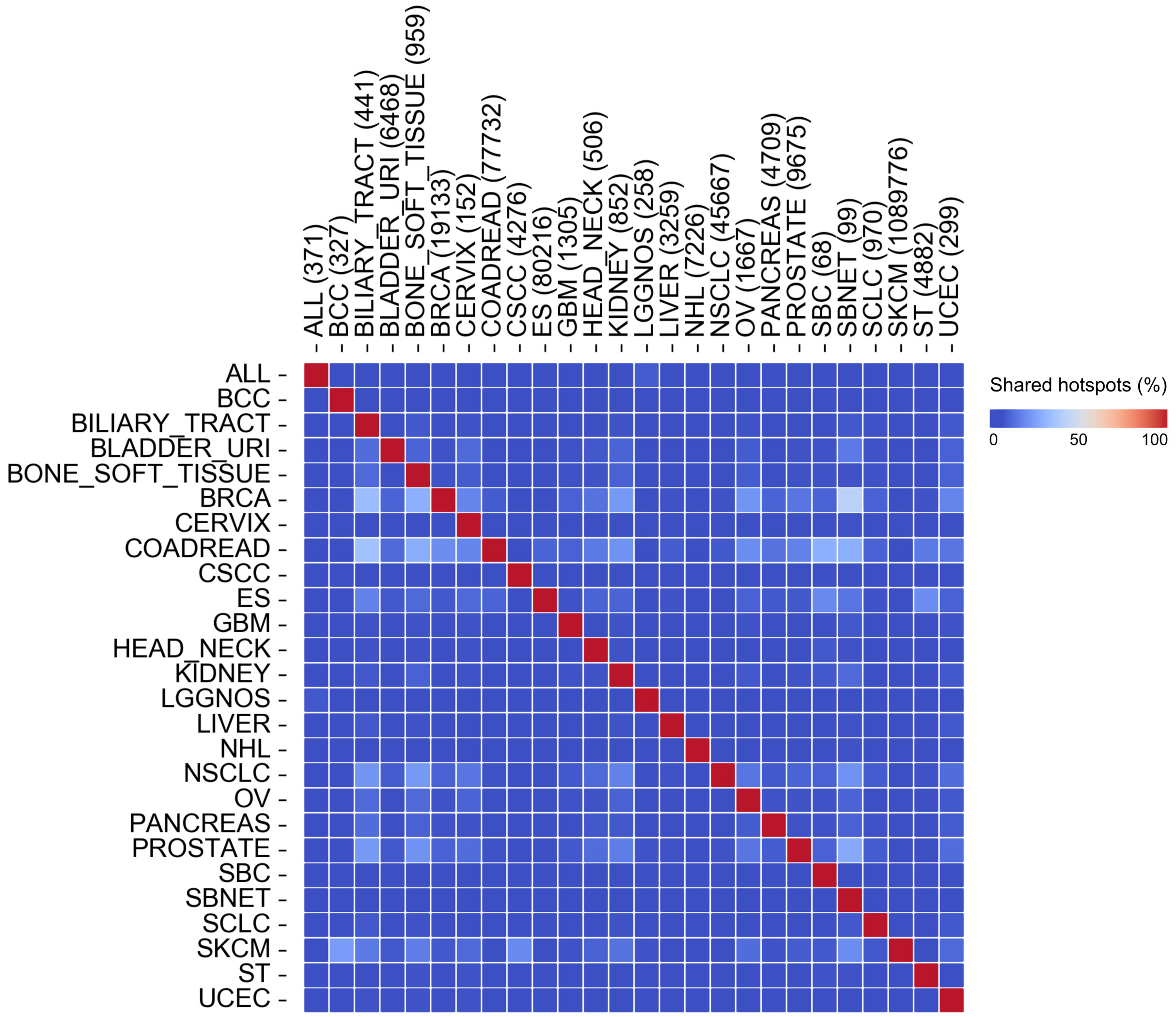


**Supp. Fig. 8. Hotspots shared between cancer types.** Heatmap depicting the fraction of hotspots shared between cancer types with at least 100 hotspots. For each cancer type pair, the colour of the box depicts the fraction of hotspots shared with respect to the cancer type shown in the columns. The number of hotspots per cancer type is shown on top.

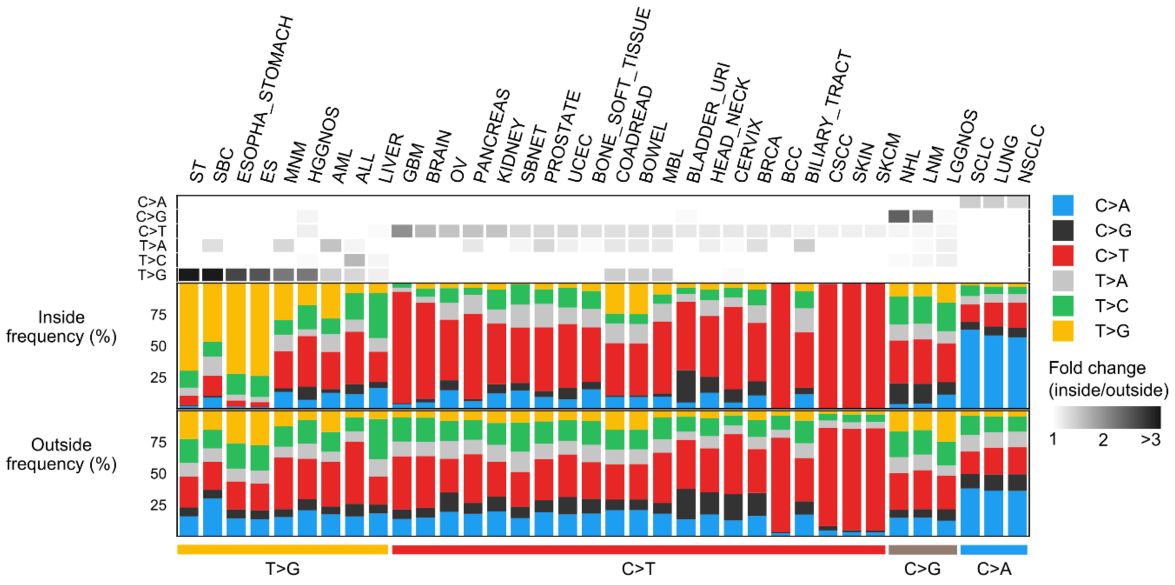


**Supp. Fig. 9. Hotspot enrichment across six pyrimidine-based substitution types.** Enrichments (fold change > 1) of different substitution types across mutations inside versus outside hotspots across cancer types bearing 100 or more hotspot mutations. The frequency of each substitution type inside and outside hotspots are shown at the bottom panels. Cancer types appear clustered through hierarchical clustering based on their fold-changes across substitution types. Cancer types within clusters are sorted following the descending magnitude of fold change.

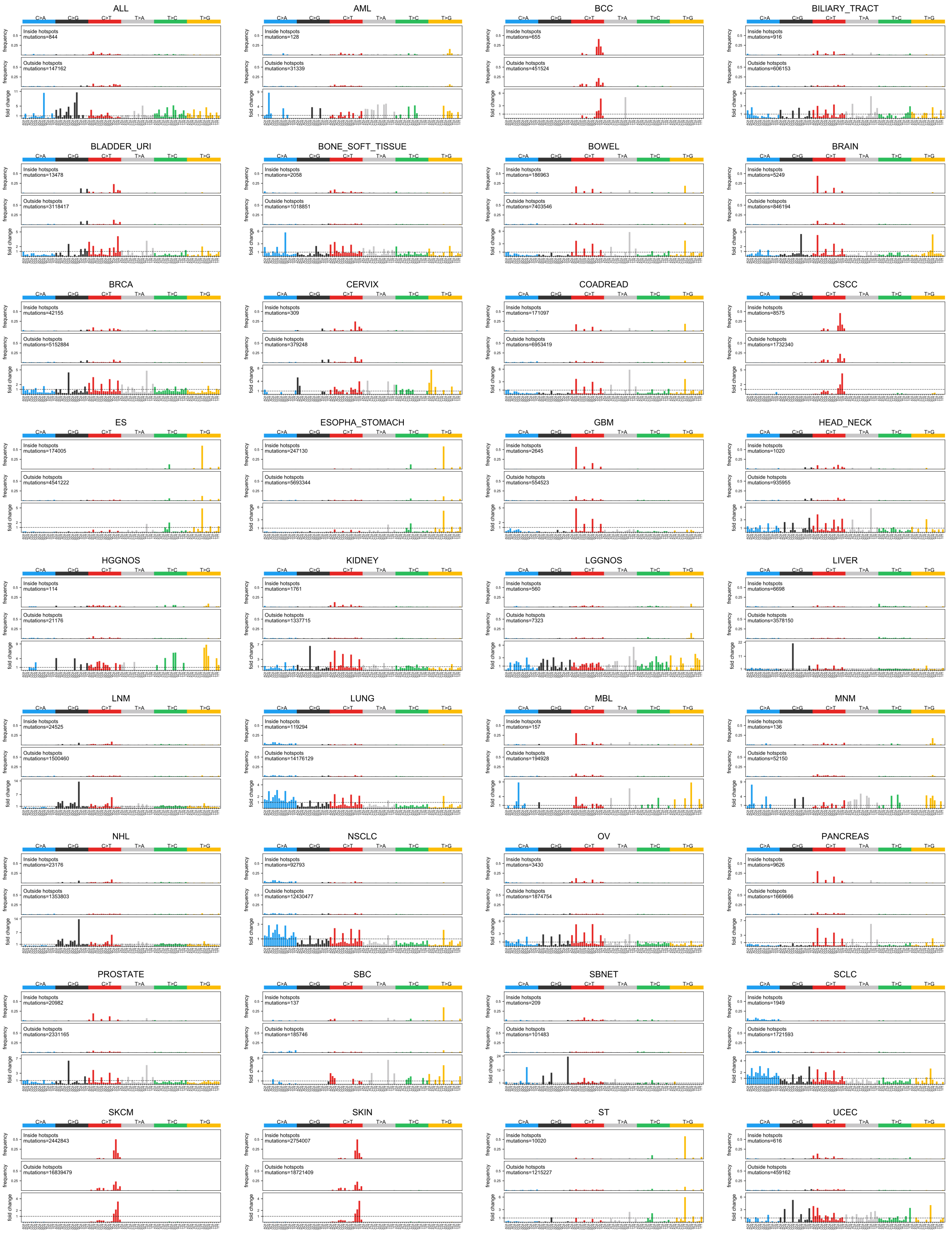
ALL
Inside hotspots mutations=844
Outside hotspots mutations=147162
AML
Inside hotspots mutations=128
Outside hotspots mutations=31339
BCC
Inside hotspots mutations=655
Outside hotspots mutations=451524
BILIARY_TRACT
Inside hotspots mutations=916
Outside hotspots mutations=606153
BLADDER_URI
Inside hotspots mutations=13478
Outside hotspots mutations=3118417
BONE_SOFT_TISSUE
Inside hotspots mutations=2058
Outside hotspots mutations=1018851
BOWEL
Inside hotspots mutations=186963
Outside hotspots mutations=7403546
BRAIN
Inside hotspots mutations=5249
Outside hotspots mutations=846194
BRCA
Inside hotspots mutations=42155
Outside hotspots mutations=5152884
CERVIX
Inside hotspots mutations=309
Outside hotspots mutations=379248
COADREAD
Inside hotspots mutations=171097
Outside hotspots mutations=6953419
CSCC
Inside hotspots mutations=8575
Outside hotspots mutations=1732340
ES
Inside hotspots mutations=174005
Outside hotspots mutations=4541222
ESOPHA_STOMACH
Inside hotspots mutations=247130
Outside hotspots mutations=5693344
GBM
Inside hotspots mutations=2645
Outside hotspots mutations=554523
HEAD_NECK
Inside hotspots mutations=1020
Outside hotspots mutations=935955
HGGNOS
Inside hotspots mutations=114
Outside hotspots mutations=21176
KIDNEY
Inside hotspots mutations=1761
Outside hotspots mutations=1337715
LGGNOS
Inside hotspots mutations=560
Outside hotspots mutations=7323
LIVER
Inside hotspots mutations=6698
Outside hotspots mutations=3578150
LNM
Inside hotspots mutations=24525
Outside hotspots mutations=1500460
LUNG
Inside hotspots mutations=119294
Outside hotspots mutations=14176129
MBL
Inside hotspots mutations=157
Outside hotspots mutations=194928
MNM
Inside hotspots mutations=136
Outside hotspots mutations=52150
NHL
Inside hotspots mutations=23176
Outside hotspots mutations=1353803
NSCLC
Inside hotspots mutations=92793
Outside hotspots mutations=12430477
OV
Inside hotspots mutations=3430
Outside hotspots mutations=1874754
PANCREAS
Inside hotspots mutations=9626
Outside hotspots mutations=1669666
PROSTATE
Inside hotspots mutations=20982
Outside hotspots mutations=2331165
SBC
Inside hotspots mutations=137
Outside hotspots mutations=185746
SBNET
Inside hotspots mutations=209
Outside hotspots mutations=101483
SCLC
Inside hotspots mutations=1949
Outside hotspots mutations=1721593
SKCM
Inside hotspots mutations=2442843
Outside hotspots mutations=16839479
SKIN
Inside hotspots mutations=2754007
Outside hotspots mutations=18721409
ST
Inside hotspots mutations=10020
Outside hotspots mutations=1215227
UCEC
Inside hotspots mutations=616
Outside hotspots mutations=459162
C>A C>G C>T T>A T>C T>G
frequency
fold change

**Supp. Fig. 10. Hotspot enrichment across 96-based trinucleotide single base substitutions.** Frequencies of hotspot (top) versus non-hotspot (middle) mutations across the 96 types of trinucleotide based substitutions (pyrimidine-based substitutions considering all possible combinations of nucleotides immediately upstream and downstream the mutated base). Inside-to-outside fold changes across trinucleotides are shown at the bottom panel of each plot. Fold changes above 1 (dashed grey line) denote an increase of mutations inside hotspots over the background (outside) mutation rate of the particular trinucleotide. Only cancer types with at least 100 hotspot mutations are shown.

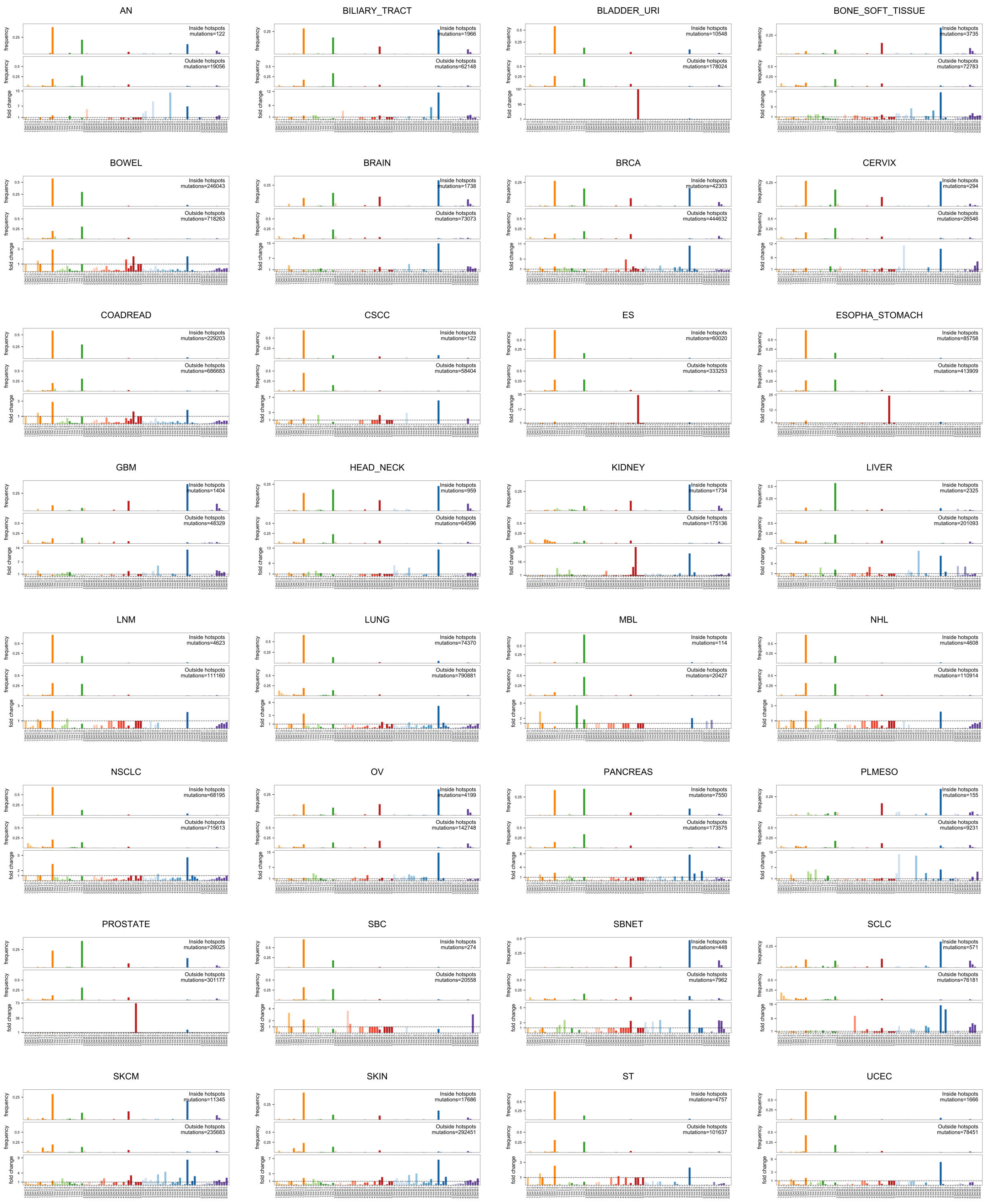
AN
Inside hotspots mutations=122
Outside hotspots mutations=19056
BILIARY_TRACT
Inside hotspots mutations=1966
Outside hotspots mutations=62148
BLADDER_URI
Inside hotspots mutations=10548
Outside hotspots mutations=178024
BONE_SOFT_TISSUE
Inside hotspots mutations=3735
Outside hotspots mutations=72783
BOWEL
Inside hotspots mutations=246043
Outside hotspots mutations=718263
BRAIN
Inside hotspots mutations=1738
Outside hotspots mutations=73073
BRCA
Inside hotspots mutations=42303
Outside hotspots mutations=444632
CERVIX
Inside hotspots mutations=294
Outside hotspots mutations=26546
COADREAD
Inside hotspots mutations=229203
Outside hotspots mutations=686683
CSCC
Inside hotspots mutations=122
Outside hotspots mutations=58404
ES
Inside hotspots mutations=60020
Outside hotspots mutations=333253
ESOPHA_STOMACH
Inside hotspots mutations=85758
Outside hotspots mutations=413909
GBM
Inside hotspots mutations=1404
Outside hotspots mutations=48329
HEAD_NECK
Inside hotspots mutations=959
Outside hotspots mutations=64596
KIDNEY
Inside hotspots mutations=1734
Outside hotspots mutations=175136
LIVER
Inside hotspots mutations=2325
Outside hotspots mutations=201093
LNM
Inside hotspots mutations=4623
Outside hotspots mutations=111160
LUNG
Inside hotspots mutations=74370
Outside hotspots mutations=790881
MBL
Inside hotspots mutations=114
Outside hotspots mutations=20427
NHL
Inside hotspots mutations=4608
Outside hotspots mutations=110914
NSCLC
Inside hotspots mutations=68195
Outside hotspots mutations=715613
OV
Inside hotspots mutations=4199
Outside hotspots mutations=142748
PANCREAS
Inside hotspots mutations=7550
Outside hotspots mutations=173575
PLMESO
Inside hotspots mutations=155
Outside hotspots mutations=9231
PROSTATE
Inside hotspots mutations=28025
Outside hotspots mutations=301177
SBC
Inside hotspots mutations=274
Outside hotspots mutations=20558
SBNET
Inside hotspots mutations=448
Outside hotspots mutations=7962
SCLC
Inside hotspots mutations=571
Outside hotspots mutations=76181
SKCM
Inside hotspots mutations=11345
Outside hotspots mutations=235683
SKIN
Inside hotspots mutations=17686
Outside hotspots mutations=292451
ST
Inside hotspots mutations=4757
Outside hotspots mutations=101637
UCEC
Inside hotspots mutations=1666
Outside hotspots mutations=78451
frequency
fold change

**Supp. Fig. 11. Hotspot enrichment across indel categories.** Frequencies of hotspot (top) versus non-hotspot (middle) mutations across the 83-based indel categories. Inside-to-outside fold changes across each class are shown at the bottom panel of each plot. Fold changes above 1 (dashed grey line) denote an increase of mutations inside hotspots over the background (outside) mutation rate of the particular trinucleotide. Only cancer types with at least 100 indel hotspot mutations are shown.

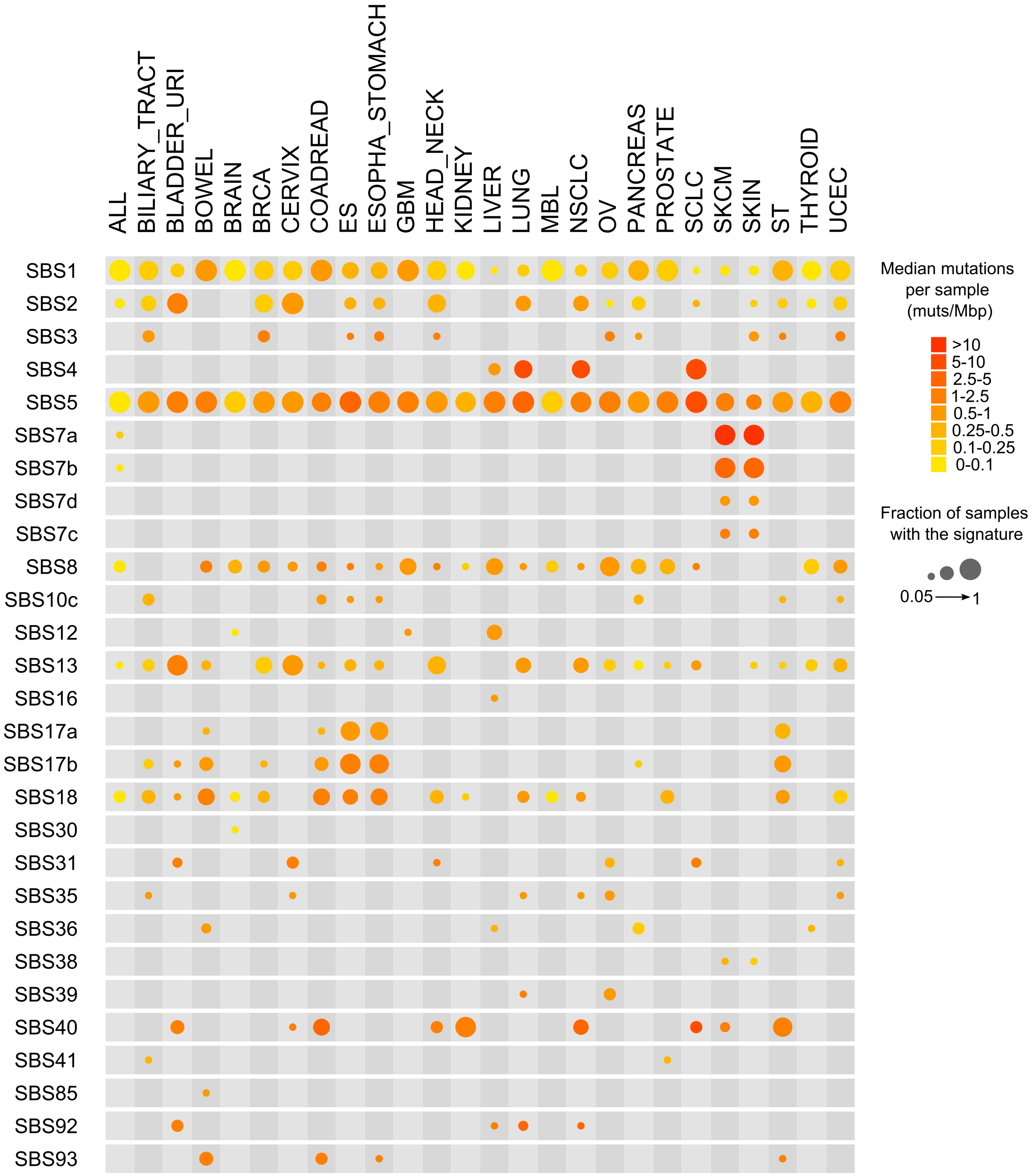


**Supp. Fig. 12. COSMIC reference signatures set decomposition of extracted SBS signatures.** Heatmap showing the decomposition of *de novo* extracted signatures into the COSMIC v3.2 reference set. Dots show the fraction of samples of the cancer type in which the signature is found to be active. Colour shows the median mutation burden per signature across samples where the signature is active (this is, 5% or more mutations within the sample are attributable to the signature). Only signatures active in 5% or more of the samples in at least one cancer type are shown.

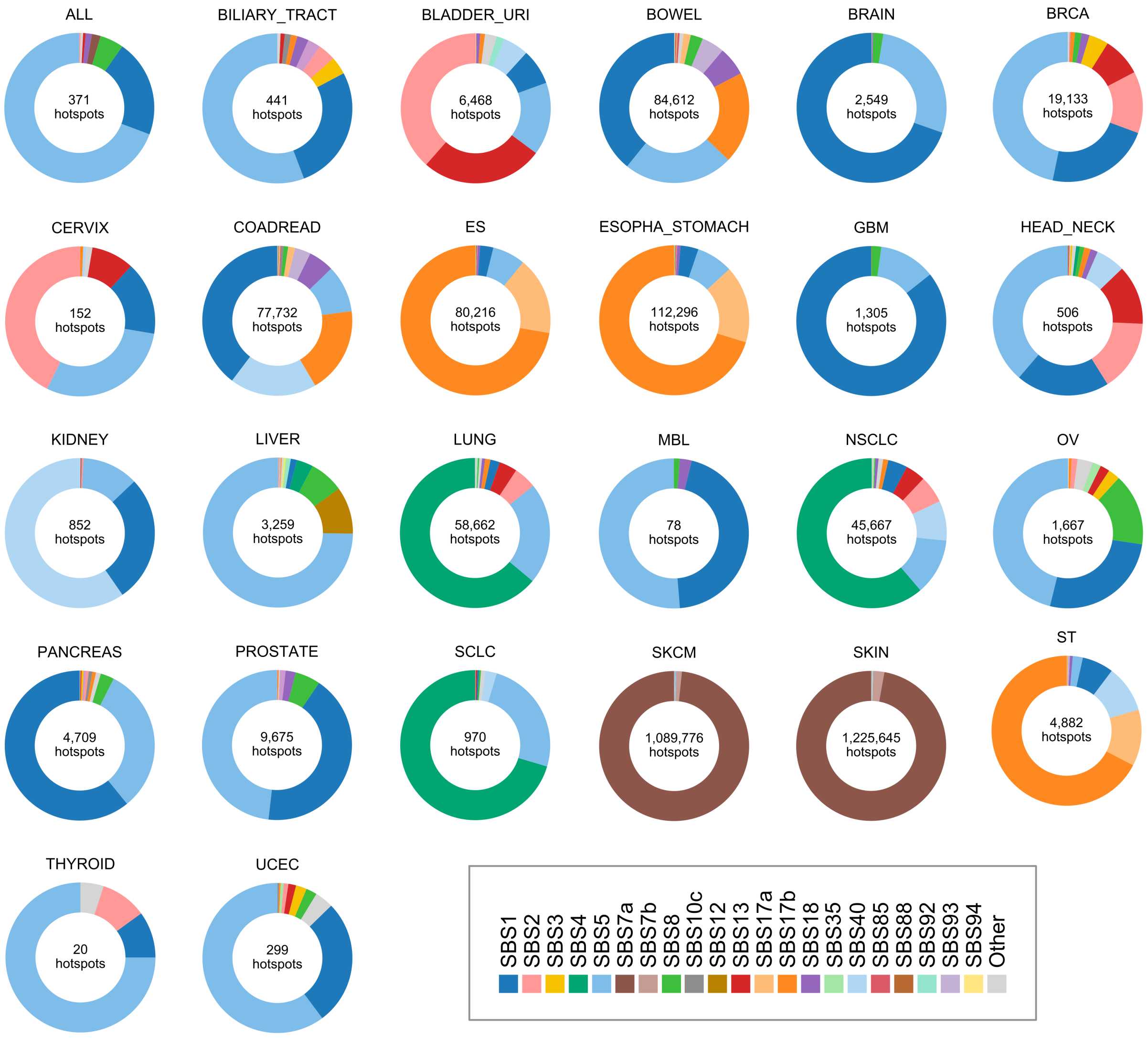


**Supp. Fig. 13. Proportion of hotspots per signature across cancers.** Pie charts showing the proportion of hotspots attributable to signatures active in each cancer type (Methods).

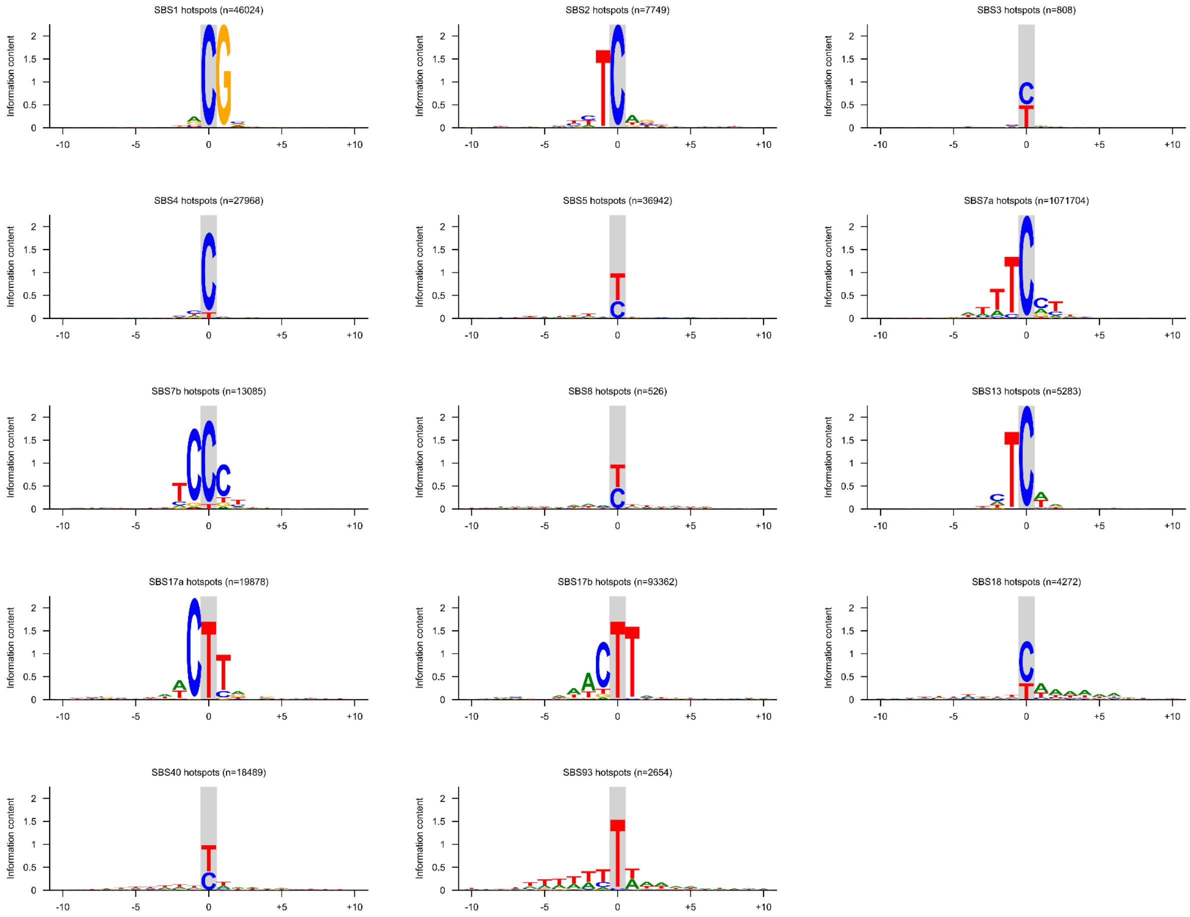


**Supp. Fig. 14. Hotspot logos across signatures.** Logos depicting the information content of the normalised nucleotide frequency per position in a 21 bp window centred at the nucleotide with the hotspot (grey vertical line). For each signature, hotspots were merged across cancer types (Methods).

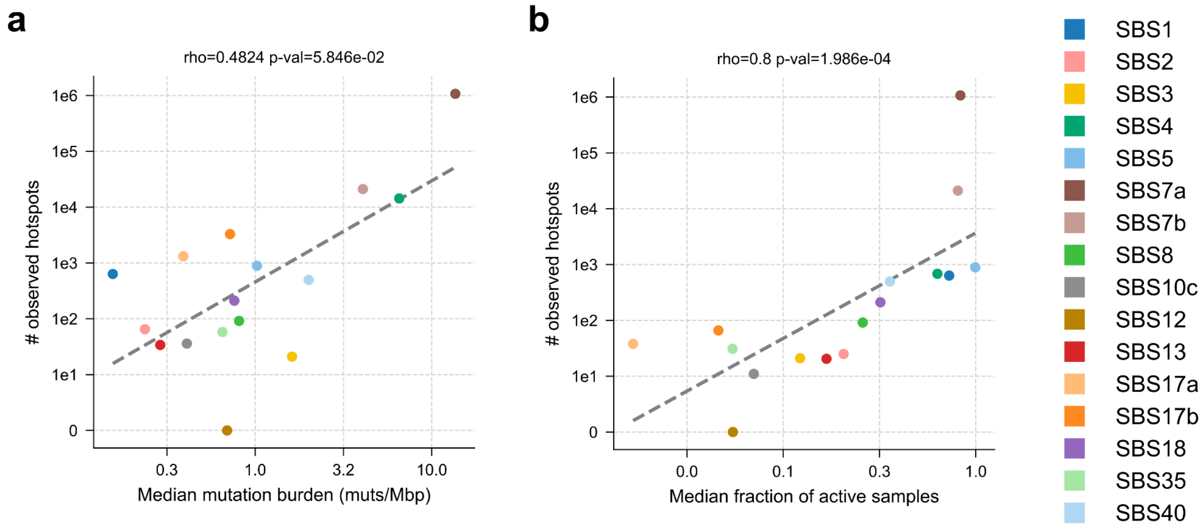


**Supp. Fig. 15. Relationship between hotspot burden, mutation burden and samples with activity of each signature. a**) Number of observed hotspots versus the median number of mutations attributable to the signature across cancer types where it is active. **b**) Number of observed hotspots versus the median fraction of active samples across cancer types where the signature is active. Dashed lines show the fitting of an linear regression (OLS) model on the data. Spearman's correlation coefficients (rho) are shown on top.

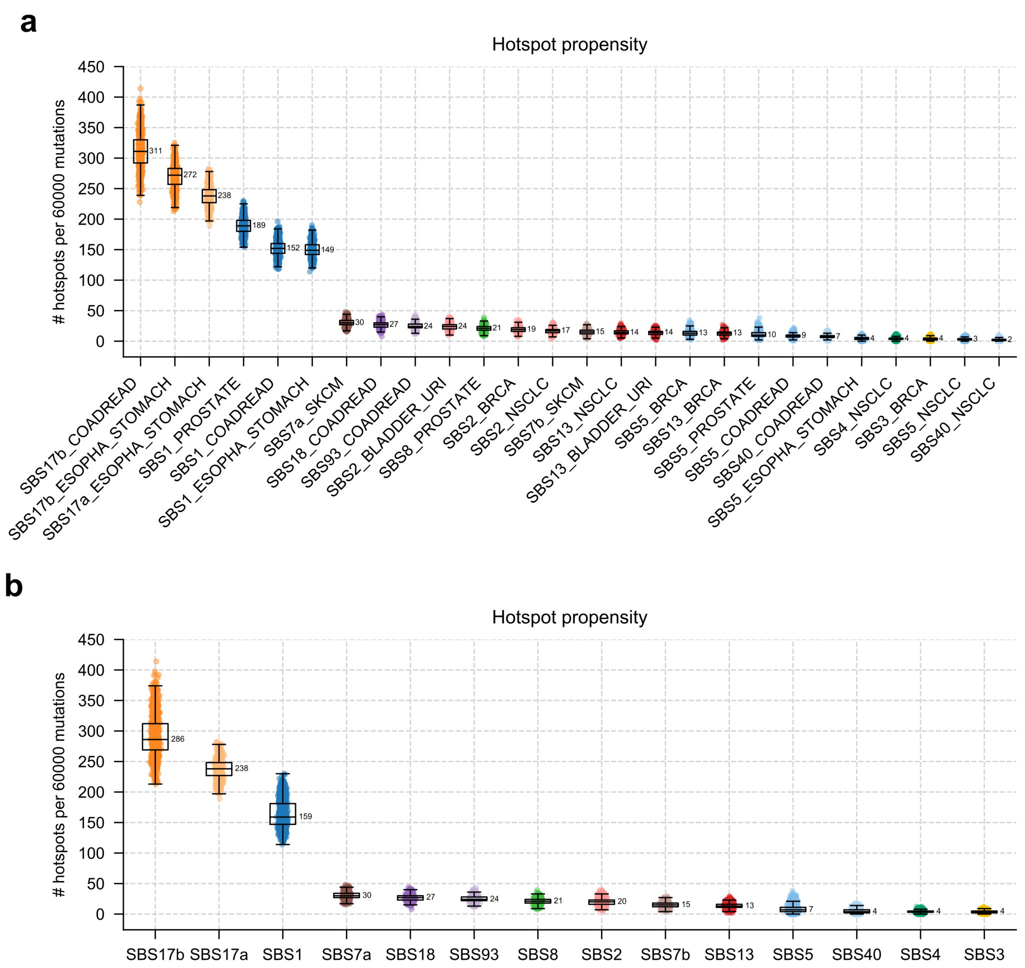


**Supp. Fig. 16. Hotspot propensity with increased mutation burden. a**) Number of hotspots per signature and cancer type observed by subsampling 60,000 total mutations (600 mutations/sample, 100 samples). **b**) Number of hotspots per signature observed within 60,000 subsampled mutations (600 mutations/sample, 100 samples) across cancer types merging data shown in a. Error bars in a and b show 1.5 times the IQR below and above 1st and 3rd quartiles, respectively. Signatures are sorted in descending order according to the mean number of observed hotspots. Note that the number of cancer types in this experiment differs from that shown in Fig. 2 where we used a subsampling of 30,000 total mutations (300 mutations/sample, 100 samples). Specifically, the following signature-cancer type pairs could not be analysed do to insufficient sampling size: SBS1-BLADDER_URI, SBS1-BRCA, SBS1-NSCLC, SBS5-BLADDER_URI, and SBS17a-COADREAD.

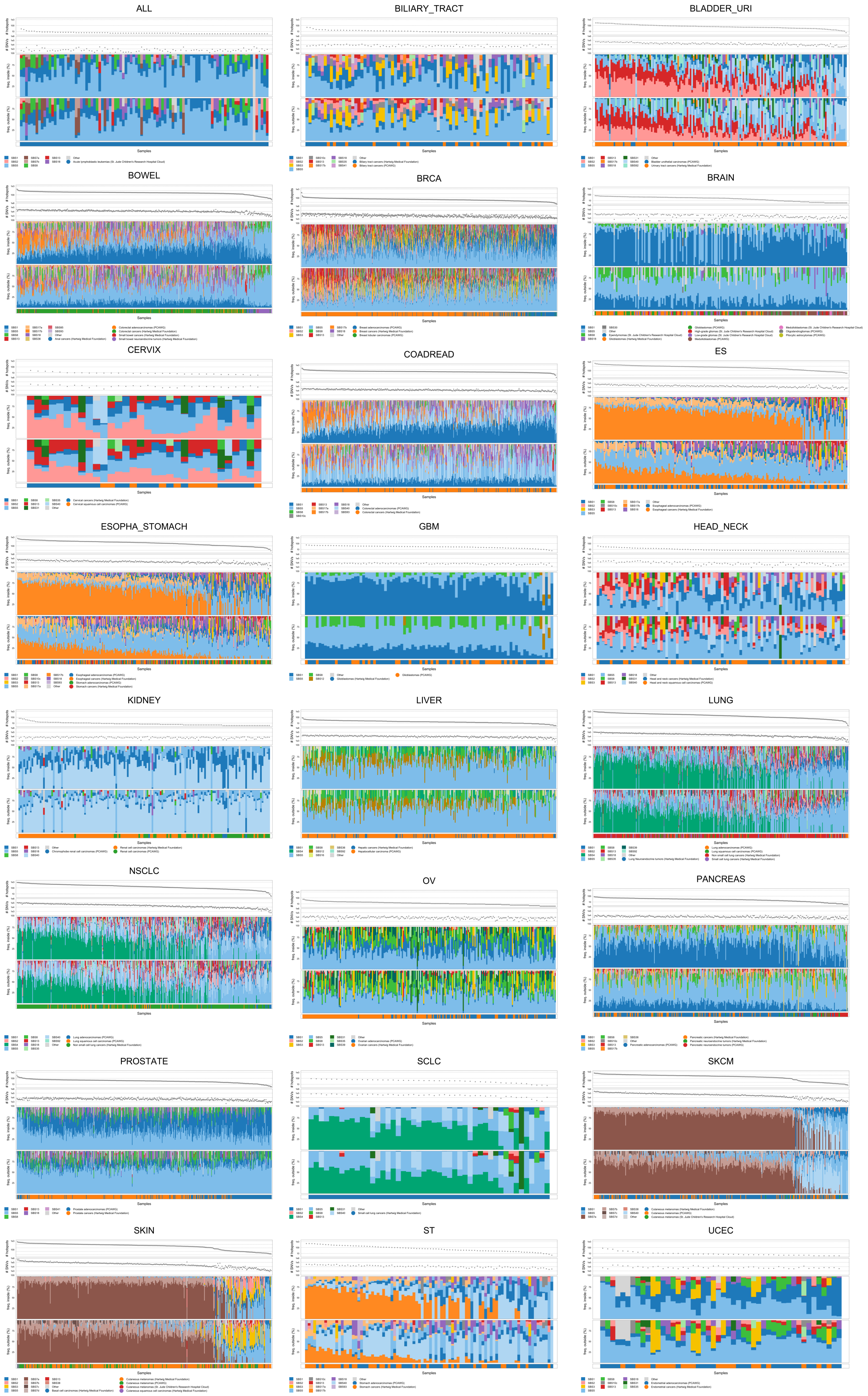
ALL
BILIARY_TRACT
BLADDER_URI
BOWEL
BRCA
BRAIN
CERVIX
COADREAD
ES
ESOPHA_STOMACH
GBM
HEAD_NECK
KIDNEY
LIVER
LUNG
NSCLC
OV
PANCREAS
PROSTATE
SCLC
SKCM
SKIN
ST
UCEC
Samples

**Supp. Fig. 17. Signature frequency inside and outside hotspots.** Barplots of the relative frequency of mutational signatures per sample within mutations inside (top) and outside hotspots (bottom) per cancer type. Signatures that are active in the cancer type (those that are present in 5% or more samples with a frequency larger than 5% of mutations per sample) are coloured following each legend, or labelled as "Other". Only samples with 3 or more hotspots are shown. MBL and THYROID cancer types, among which only 8 and 2 samples show 3 or more hotspots, respectively, are not included.

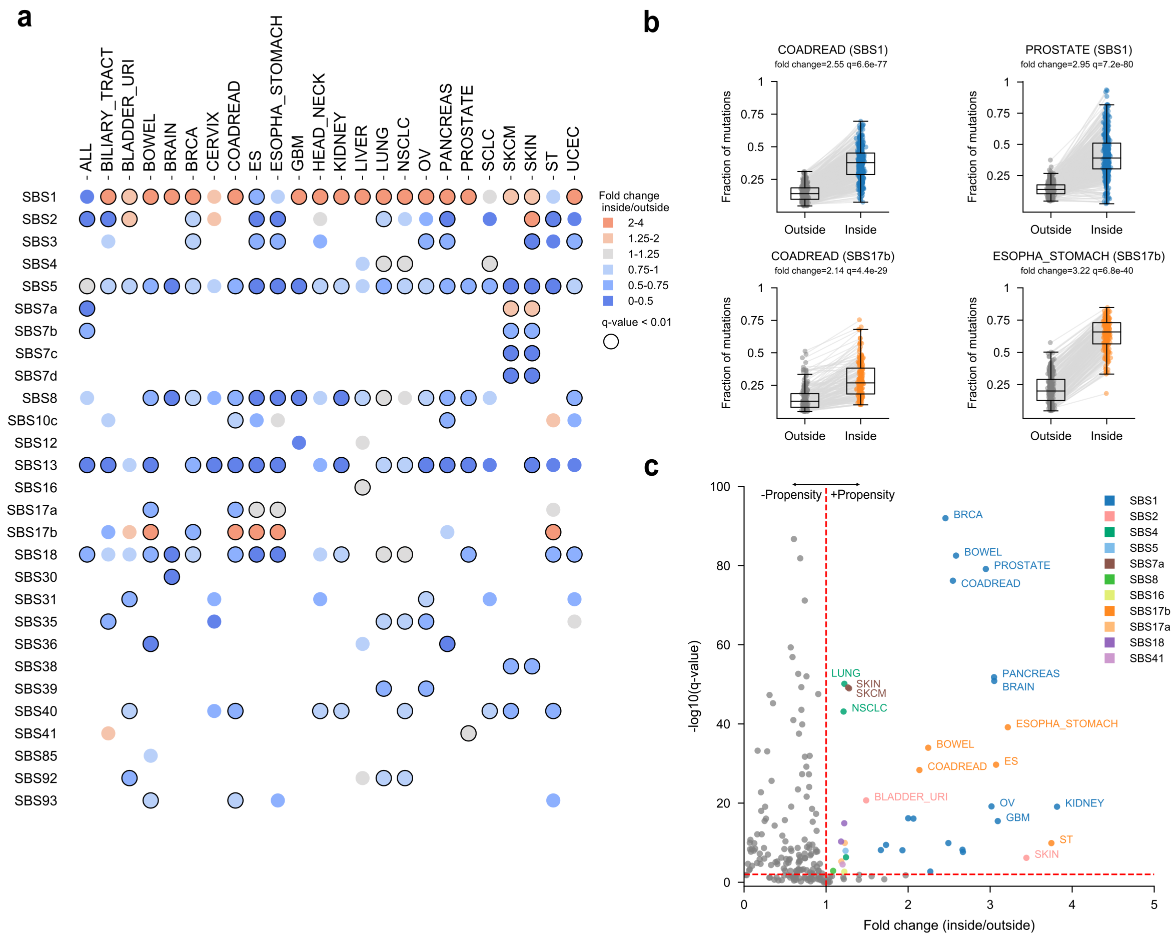


**Supp. Fig. 18. Propensity of signatures to form hotspots across cancer types measured through inside-to-outside fold-change of their activity. a**) Heatmap showing the inside-to-outside fold change across active signatures (present in 5% or more samples and contributing more than 5% of mutations of each sample). Signatures enriched in hotspots show fold-change greater than 1 (grey) or 1.25 (red); signatures that are depleted in hotspots show fold changes smaller than 1 (blue). The differences in signature frequency between inside and outside mutations were compared using Wilcoxon rank-sum test and were corrected for multiple testing (Methods). Significant fold-changes (after multiple test correction) are shown in bold. **b**) Illustration of the Wilcoxon rank-sum test of inside-to-outside signatures activity. Boxplots depicting signature frequencies per sample across mutations outside and inside hotspots. Whiskers extend 1.5 times the IQR below and above 1st and 3rd quartiles of the distribution. Grey lines connect data originating from the same sample. Fold changes and q-values computed from these comparisons are then shown in panels a and c. **c**) Scatter plot showing the propensity of signatures to form hotspots computed as the difference (fold-change, x axis and significance, y axis) between their activities outside and inside hotspots per cancer type. Signature-cancer type pairs showing significant differences (q<0.01) and fold change greater than 1 are shown in colour.

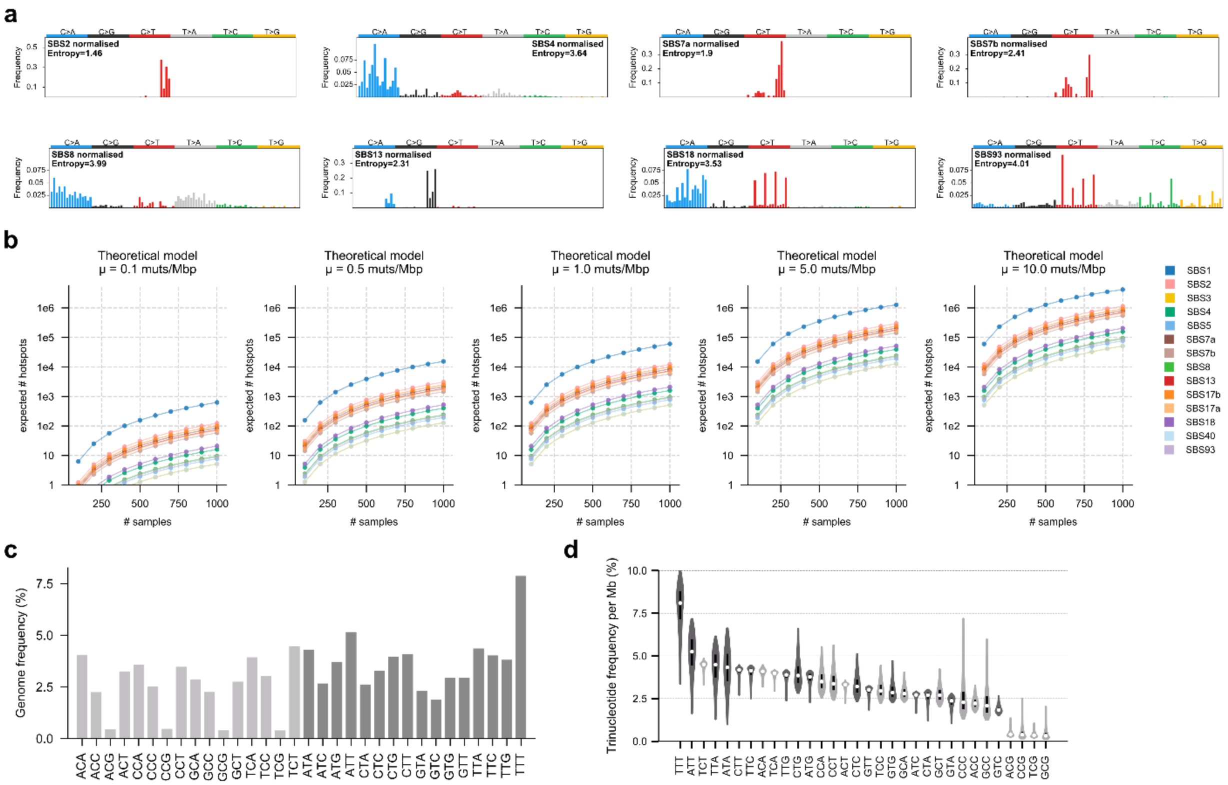


**Supp. Fig. 19. Influence of trinucleotide frequencies on hotspot propensity. a**) Normalised trinucleotide profiles of the 8 additional selected signatures under analysis (see Methods) and their entropies. **b**) Theoretical number of expected hotspots across different mutation rates (0.1-10 mutations per sample per megabase) and sample sizes (100-1,000 samples) per signature calculated using the model of homogeneous distribution of trinucleotide-specific mutation rates across the genome (Methods; Supplementary Note 5). Only positions within the mappable megabases were considered. The chosen mutation rates reflect a wide range of observed mutation rates across malignancies (Lawrence et al., 2013; Alexandrov et al., 2013). **c**) Frequency of the 32 pyrimidine-based trinucleotides across mappable genome megabases. **d**) Violin plots showing the frequency of trinucleotides across mappable megabases. White dots show the median trinucleotide frequency among all megabases.

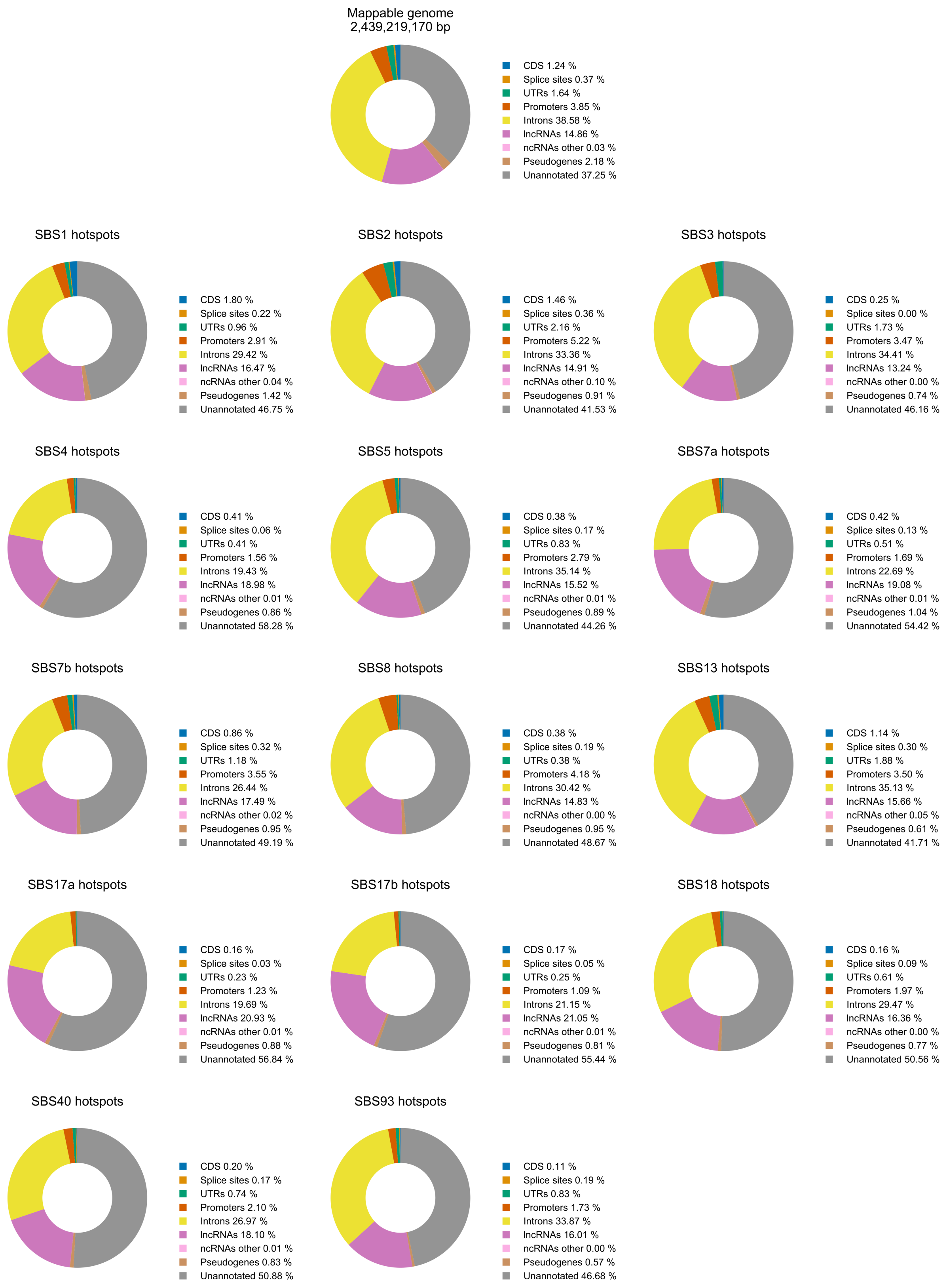
Mappable genome
2,439,219,170 bp
CDS 1.24 %
Splice sites 0.37 %
UTRs 1.64 %
Promoters 3.85 %
Introns 38.58 %
lncRNAs 14.86 %
ncRNAs other 0.03 %
Pseudogenes 2.18 %
Unannotated 37.25 %
SBS1 hotspots
CDS 1.80 %
Splice sites 0.22 %
UTRs 0.96 %
Promoters 2.91 %
Introns 29.42 %
lncRNAs 16.47 %
ncRNAs other 0.04 %
Pseudogenes 1.42 %
Unannotated 46.75 %
SBS2 hotspots
CDS 1.46 %
Splice sites 0.36 %
UTRs 2.16 %
Promoters 5.22 %
Introns 33.36 %
lncRNAs 14.91 %
ncRNAs other 0.10 %
Pseudogenes 0.91 %
Unannotated 41.53 %
SBS3 hotspots
CDS 0.25 %
Splice sites 0.00 %
UTRs 1.73 %
Promoters 3.47 %
Introns 34.41 %
lncRNAs 13.24 %
ncRNAs other 0.00 %
Pseudogenes 0.74 %
Unannotated 46.16 %
SBS4 hotspots
CDS 0.41 %
Splice sites 0.06 %
UTRs 0.41 %
Promoters 1.56 %
Introns 19.43 %
lncRNAs 18.98 %
ncRNAs other 0.01 %
Pseudogenes 0.86 %
Unannotated 58.28 %
SBS5 hotspots
CDS 0.38 %
Splice sites 0.17 %
UTRs 0.83 %
Promoters 2.79 %
Introns 35.14 %
lncRNAs 15.52 %
ncRNAs other 0.01 %
Pseudogenes 0.89 %
Unannotated 44.26 %
SBS7a hotspots
CDS 0.42 %
Splice sites 0.13 %
UTRs 0.51 %
Promoters 1.69 %
Introns 22.69 %
lncRNAs 19.08 %
ncRNAs other 0.01 %
Pseudogenes 1.04 %
Unannotated 54.42 %
SBS7b hotspots
CDS 0.86 %
Splice sites 0.32 %
UTRs 1.18 %
Promoters 3.55 %
Introns 26.44 %
lncRNAs 17.49 %
ncRNAs other 0.02 %
Pseudogenes 0.95 %
Unannotated 49.19 %
SBS8 hotspots
CDS 0.38 %
Splice sites 0.19 %
UTRs 0.38 %
Promoters 4.18 %
Introns 30.42 %
lncRNAs 14.83 %
ncRNAs other 0.00 %
Pseudogenes 0.95 %
Unannotated 48.67 %
SBS13 hotspots
CDS 1.14 %
Splice sites 0.30 %
UTRs 1.88 %
Promoters 3.50 %
Introns 35.13 %
lncRNAs 15.66 %
ncRNAs other 0.05 %
Pseudogenes 0.61 %
Unannotated 41.71 %
SBS17a hotspots
CDS 0.16 %
Splice sites 0.03 %
UTRs 0.23 %
Promoters 1.23 %
Introns 19.69 %
lncRNAs 20.93 %
ncRNAs other 0.01 %
Pseudogenes 0.88 %
Unannotated 56.84 %
SBS17b hotspots
CDS 0.17 %
Splice sites 0.05 %
UTRs 0.25 %
Promoters 1.09 %
Introns 21.15 %
lncRNAs 21.05 %
ncRNAs other 0.01 %
Pseudogenes 0.81 %
Unannotated 55.44 %
SBS18 hotspots
CDS 0.16 %
Splice sites 0.09 %
UTRs 0.61 %
Promoters 1.97 %
Introns 29.47 %
lncRNAs 16.36 %
ncRNAs other 0.00 %
Pseudogenes 0.77 %
Unannotated 50.56 %
SBS40 hotspots
CDS 0.20 %
Splice sites 0.17 %
UTRs 0.74 %
Promoters 2.10 %
Introns 26.97 %
lncRNAs 18.10 %
ncRNAs other 0.01 %
Pseudogenes 0.83 %
Unannotated 50.88 %
SBS93 hotspots
CDS 0.11 %
Splice sites 0.19 %
UTRs 0.83 %
Promoters 1.73 %
Introns 33.87 %
lncRNAs 16.01 %
ncRNAs other 0.00 %
Pseudogenes 0.57 %
Unannotated 46.68 %

**Supp. Fig. 20. Hotspots overlapping genomic elements.** Proportion of observed hotspots overlapping different categories of annotated genomic elements across signatures. The annotations consist of non-overlapping categories of genomic elements as defined in HotspotFinder (Supplementary Note 1) according to the following hierarchy, where CDS has the largest priority: CDS, splice sites, UTRs, promoters, introns, and lncRNAs/other ncRNAs/pseudogenes. lncRNAs, other ncRNAs and pseudogenes do not overlap any element of the former 5 classes, however they can be overlapping among each other. Splice sites, UTRs, promoters, and introns are extracted from protein-coding genes. lncRNAs annotations include both exons, introns and promoter sequences of lncRNA genes. The proportion of the mappable genome annotated with these genomic elements is shown on top.

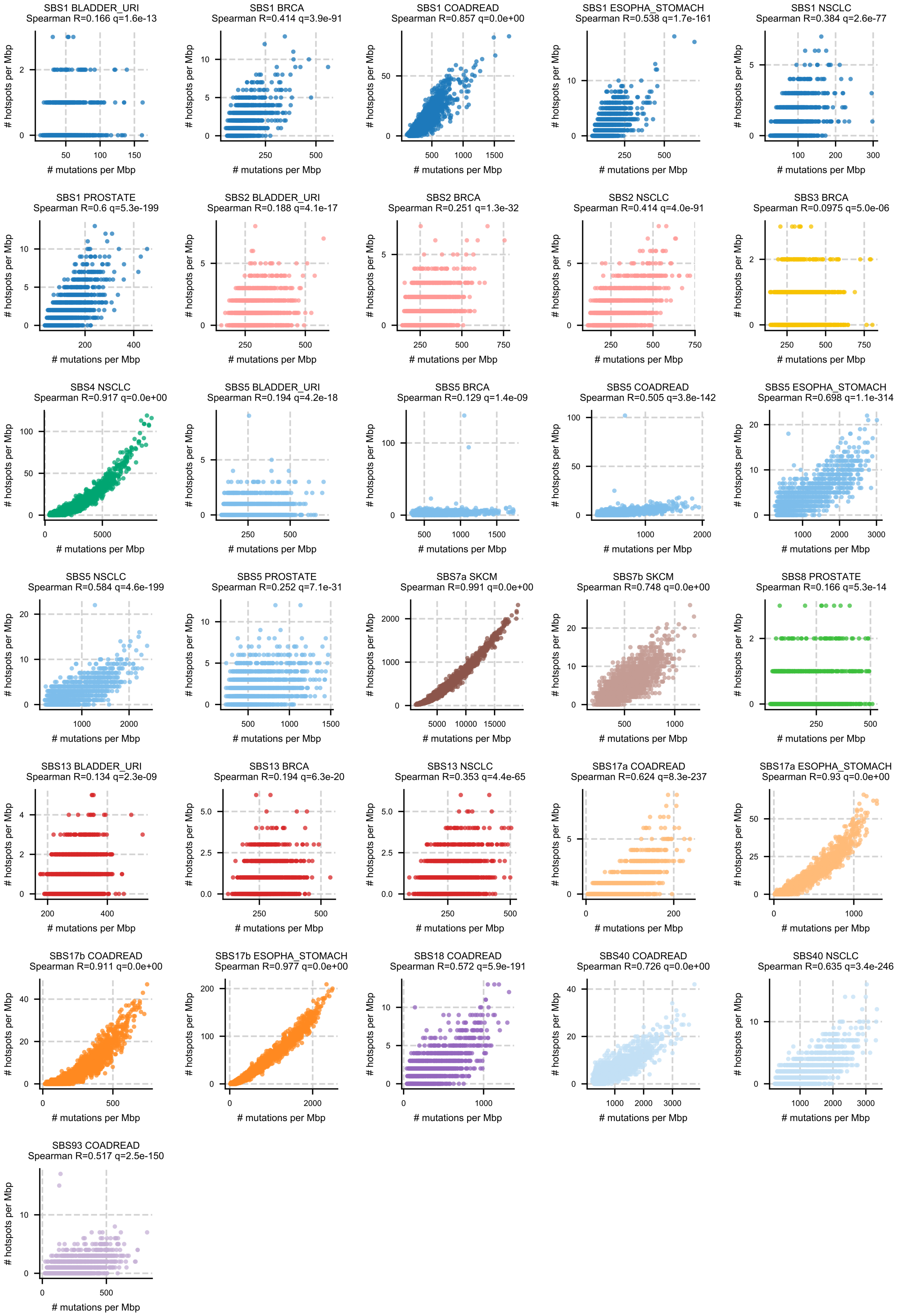
SBS1 BLADDER_URI
Spearman R=0.166 q=1.6e-13
SBS1 BRCA
Spearman R=0.414 q=3.9e-91
SBS1 COADREAD
Spearman R=0.857 q=0.0e+00
SBS1 ESOPHA_STOMACH
Spearman R=0.538 q=1.7e-161
SBS1 NSCLC
Spearman R=0.384 q=2.6e-77
SBS1 PROSTATE
Spearman R=0.6 q=5.3e-199
SBS2 BLADDER_URI
Spearman R=0.188 q=4.1e-17
SBS2 BRCA
Spearman R=0.251 q=1.3e-32
SBS2 NSCLC
Spearman R=0.414 q=4.0e-91
SBS3 BRCA
Spearman R=0.0975 q=5.0e-06
SBS4 NSCLC
Spearman R=0.917 q=0.0e+00
SBS5 BLADDER_URI
Spearman R=0.194 q=4.2e-18
SBS5 BRCA
Spearman R=0.129 q=1.4e-09
SBS5 COADREAD
Spearman R=0.505 q=3.8e-142
SBS5 ESOPHA_STOMACH
Spearman R=0.698 q=1.1e-314
SBS5 NSCLC
Spearman R=0.584 q=4.6e-199
SBS5 PROSTATE
Spearman R=0.252 q=7.1e-31
SBS7a SKCM
Spearman R=0.991 q=0.0e+00
SBS7b SKCM
Spearman R=0.748 q=0.0e+00
SBS8 PROSTATE
Spearman R=0.166 q=5.3e-14
SBS13 BLADDER_URI
Spearman R=0.134 q=2.3e-09
SBS13 BRCA
Spearman R=0.194 q=6.3e-20
SBS13 NSCLC
Spearman R=0.353 q=4.4e-65
SBS17a COADREAD
Spearman R=0.624 q=8.3e-237
SBS17a ESOPHA_STOMACH
Spearman R=0.93 q=0.0e+00
SBS17b COADREAD
Spearman R=0.911 q=0.0e+00
SBS17b ESOPHA_STOMACH
Spearman R=0.977 q=0.0e+00
SBS18 COADREAD
Spearman R=0.572 q=5.9e-191
SBS40 COADREAD
Spearman R=0.726 q=0.0e+00
SBS40 NSCLC
Spearman R=0.635 q=3.4e-246
SBS93 COADREAD
Spearman R=0.517 q=2.5e-150
### hotspots per Mbp
### mutations per Mbp

**Supp. Fig. 21. Relationship between hotspot burden and mutation burden across genome megabases.** Scatter plots showing the number of hotspots and mutations attributed to a signature in a cancer type across mappable genome megabases. Spearman's correlation between both variables is shown on top.

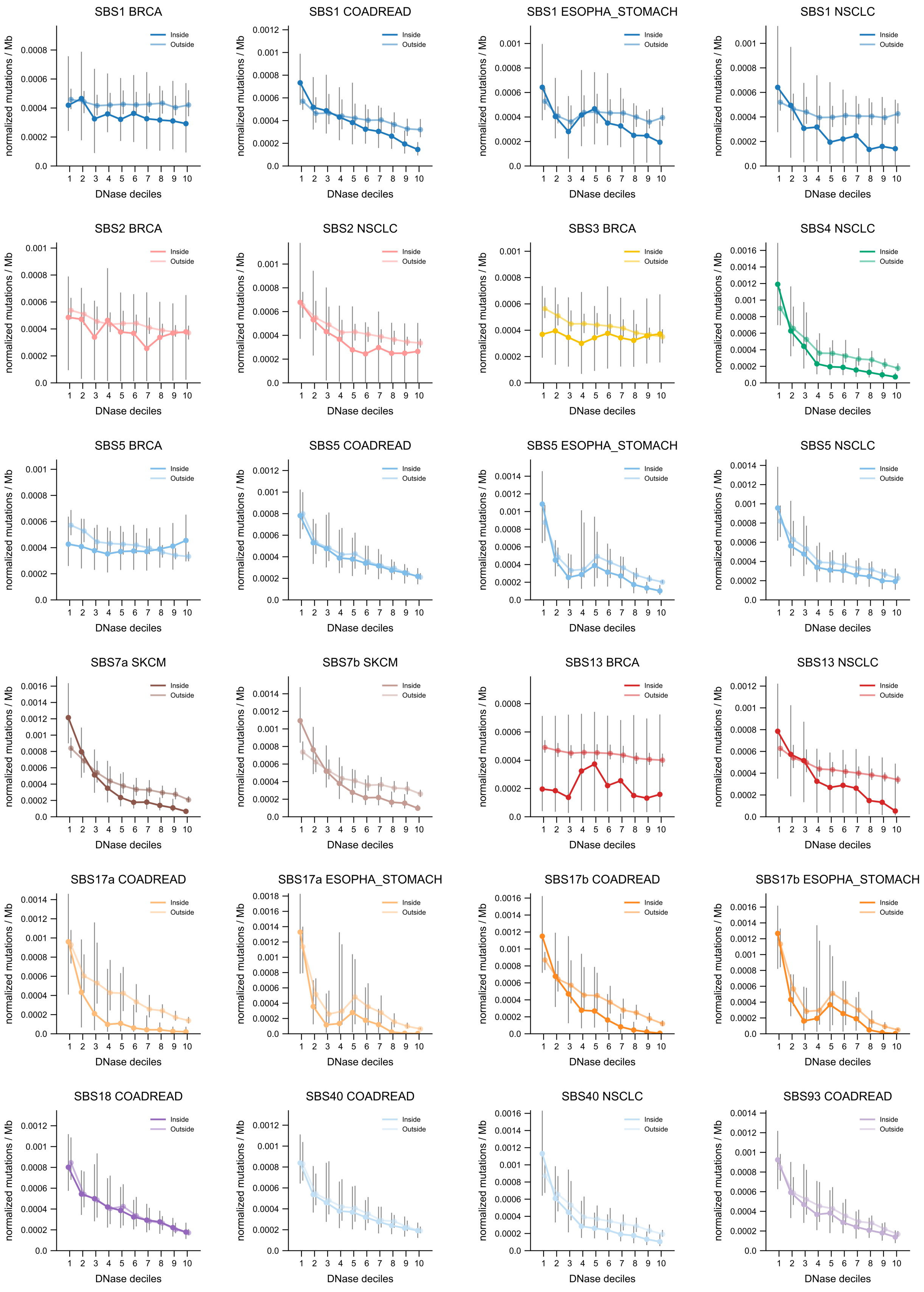
SBS1 BRCA
SBS1 COADREAD
SBS1 ESOPHA_STOMACH
SBS1 NSCLC
SBS2 BRCA
SBS2 NSCLC
SBS3 BRCA
SBS4 NSCLC
SBS5 BRCA
SBS5 COADREAD
SBS5 ESOPHA_STOMACH
SBS5 NSCLC
SBS7a SKCM
SBS7b SKCM
SBS13 BRCA
SBS13 NSCLC
SBS17a COADREAD
SBS17a ESOPHA_STOMACH
SBS17b COADREAD
SBS17b ESOPHA_STOMACH
SBS18 COADREAD
SBS40 COADREAD
SBS40 NSCLC
SBS93 COADREAD
Inside
Outside
normalized mutations / Mb
DNase deciles
1 2 3 4 5 6 7 8 9 10

**Supp. Fig. 22. Mutation rates across genomic regions with different chromatin accessibility.** Median normalised rates of observed mutations inside and outside hotspots (two lines) across megabases according to DNase signal of chromatin accessibility in tissue-matched epigenomes. Bins represent deciles of accessibility (measured as relative coverage by DNAse peaks) from low (1) to high (10). Error bars show the IQR of the distribution of mutations in each decile.

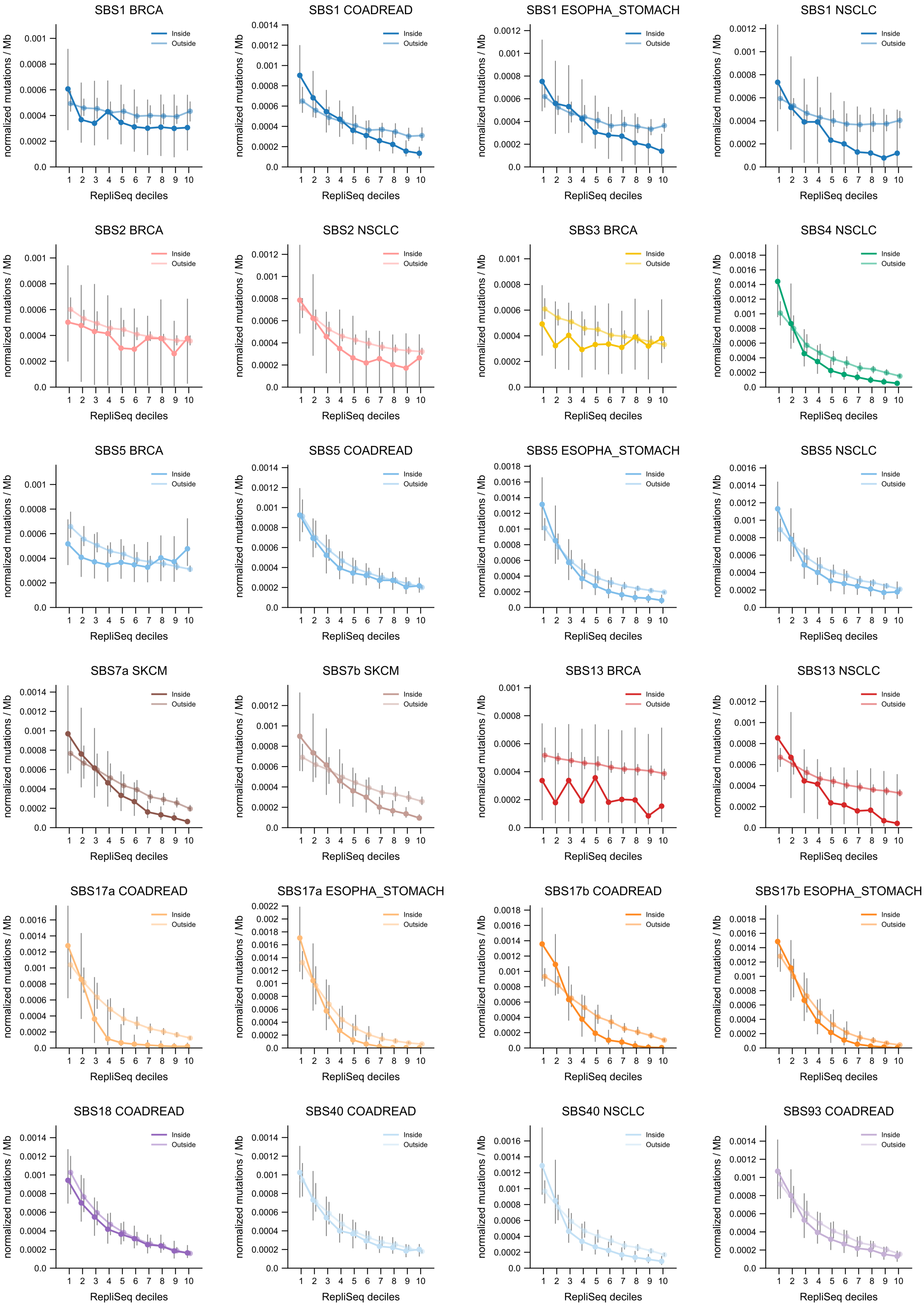
SBS1 BRCA
SBS1 COADREAD
SBS1 ESOPHA_STOMACH
SBS1 NSCLC
SBS2 BRCA
SBS2 NSCLC
SBS3 BRCA
SBS4 NSCLC
SBS5 BRCA
SBS5 COADREAD
SBS5 ESOPHA_STOMACH
SBS5 NSCLC
SBS7a SKCM
SBS7b SKCM
SBS13 BRCA
SBS13 NSCLC
SBS17a COADREAD
SBS17a ESOPHA_STOMACH
SBS17b COADREAD
SBS17b ESOPHA_STOMACH
SBS18 COADREAD
SBS40 COADREAD
SBS40 NSCLC
SBS93 COADREAD
Inside
Outside
normalized mutations / Mb
RepliSeq deciles

**Supp. Fig. 23. Mutation rate across genomic regions with different replication timing.** Median normalised rates of observed mutations inside and outside hotspots (two lines) across bins of Repli-Seq signal (across 7 cell lines of solid tissues). Bins represent deciles from late (1) to early (10) replication timing. Error bars show the IQR of the distribution of mutations per decile.

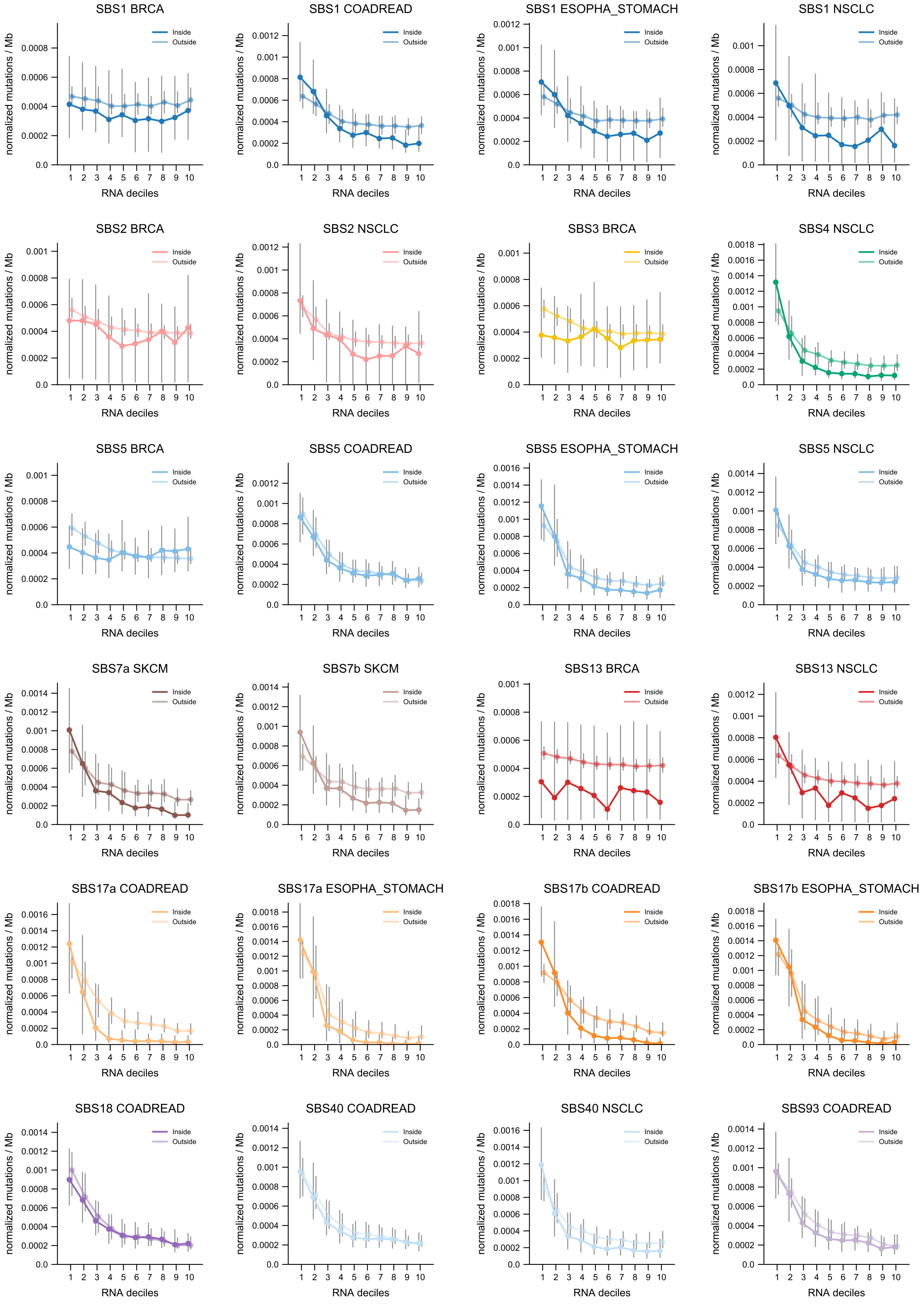
SBS1 BRCA
SBS1 COADREAD
SBS1 ESOPHA_STOMACH
SBS1 NSCLC
SBS2 BRCA
SBS2 NSCLC
SBS3 BRCA
SBS4 NSCLC
SBS5 BRCA
SBS5 COADREAD
SBS5 ESOPHA_STOMACH
SBS5 NSCLC
SBS7a SKCM
SBS7b SKCM
SBS13 BRCA
SBS13 NSCLC
SBS17a COADREAD
SBS17a ESOPHA_STOMACH
SBS17b COADREAD
SBS17b ESOPHA_STOMACH
SBS18 COADREAD
SBS40 COADREAD
SBS40 NSCLC
SBS93 COADREAD
Inside
Outside
normalized mutations / Mb
RNA deciles

**Supp. Fig. 24. Mutation rate across genomic regions with different transcriptional activity.** Median normalised rates of observed mutations inside and outside hotspots (two lines) across megabases binned by RNA-Seq signal in tissue-matched epigenomes. Bins represent deciles of RNA-Seq signal from low (1) to high (10). Error bars show the IQR of the distribution of mutations per decile.

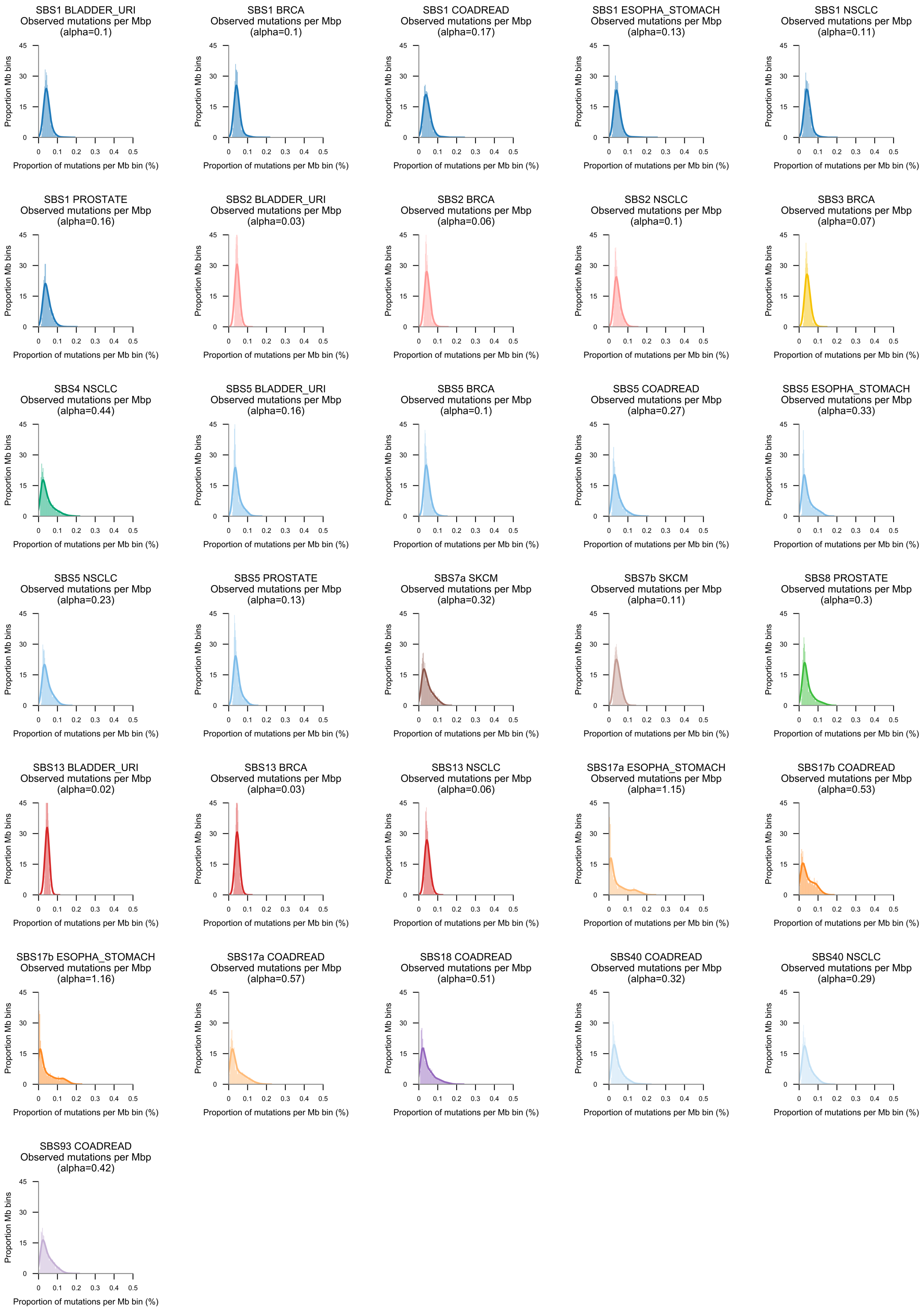
SBS1 BLADDER_URI
Observed mutations per Mbp
(alpha=0.1)
SBS1 BRCA
Observed mutations per Mbp
(alpha=0.1)
SBS1 COADREAD
Observed mutations per Mbp
(alpha=0.17)
SBS1 ESOPHA_STOMACH
Observed mutations per Mbp
(alpha=0.13)
SBS1 NSCLC
Observed mutations per Mbp
(alpha=0.11)
SBS1 PROSTATE
Observed mutations per Mbp
(alpha=0.16)
SBS2 BLADDER_URI
Observed mutations per Mbp
(alpha=0.03)
SBS2 BRCA
Observed mutations per Mbp
(alpha=0.06)
SBS2 NSCLC
Observed mutations per Mbp
(alpha=0.1)
SBS3 BRCA
Observed mutations per Mbp
(alpha=0.07)
SBS4 NSCLC
Observed mutations per Mbp
(alpha=0.44)
SBS5 BLADDER_URI
Observed mutations per Mbp
(alpha=0.16)
SBS5 BRCA
Observed mutations per Mbp
(alpha=0.1)
SBS5 COADREAD
Observed mutations per Mbp
(alpha=0.27)
SBS5 ESOPHA_STOMACH
Observed mutations per Mbp
(alpha=0.33)
SBS5 NSCLC
Observed mutations per Mbp
(alpha=0.23)
SBS5 PROSTATE
Observed mutations per Mbp
(alpha=0.13)
SBS7a SKCM
Observed mutations per Mbp
(alpha=0.32)
SBS7b SKCM
Observed mutations per Mbp
(alpha=0.11)
SBS8 PROSTATE
Observed mutations per Mbp
(alpha=0.3)
SBS13 BLADDER_URI
Observed mutations per Mbp
(alpha=0.02)
SBS13 BRCA
Observed mutations per Mbp
(alpha=0.03)
SBS13 NSCLC
Observed mutations per Mbp
(alpha=0.06)
SBS17a ESOPHA_STOMACH
Observed mutations per Mbp
(alpha=1.15)
SBS17b COADREAD
Observed mutations per Mbp
(alpha=0.53)
SBS17b ESOPHA_STOMACH
Observed mutations per Mbp
(alpha=1.16)
SBS17a COADREAD
Observed mutations per Mbp
(alpha=0.57)
SBS18 COADREAD
Observed mutations per Mbp
(alpha=0.51)
SBS40 COADREAD
Observed mutations per Mbp
(alpha=0.32)
SBS40 NSCLC
Observed mutations per Mbp
(alpha=0.29)
SBS93 COADREAD
Observed mutations per Mbp
(alpha=0.42)
Proportion Mb bins
0 15 30 45
0 0.1 0.2 0.3 0.4 0.5
Proportion of mutations per Mb bin (%)

**Supp. Fig. 25. Distribution of mutations across genome megabases.** Histograms show the fraction of mutations across mappable genome megabases. Lines represent a kernel density estimate (bandwidth=0.01) of the shape of the distribution. The overdispersion (alpha) of the underlying distribution of mutation counts across megabases is shown at the top of each plot (Methods).

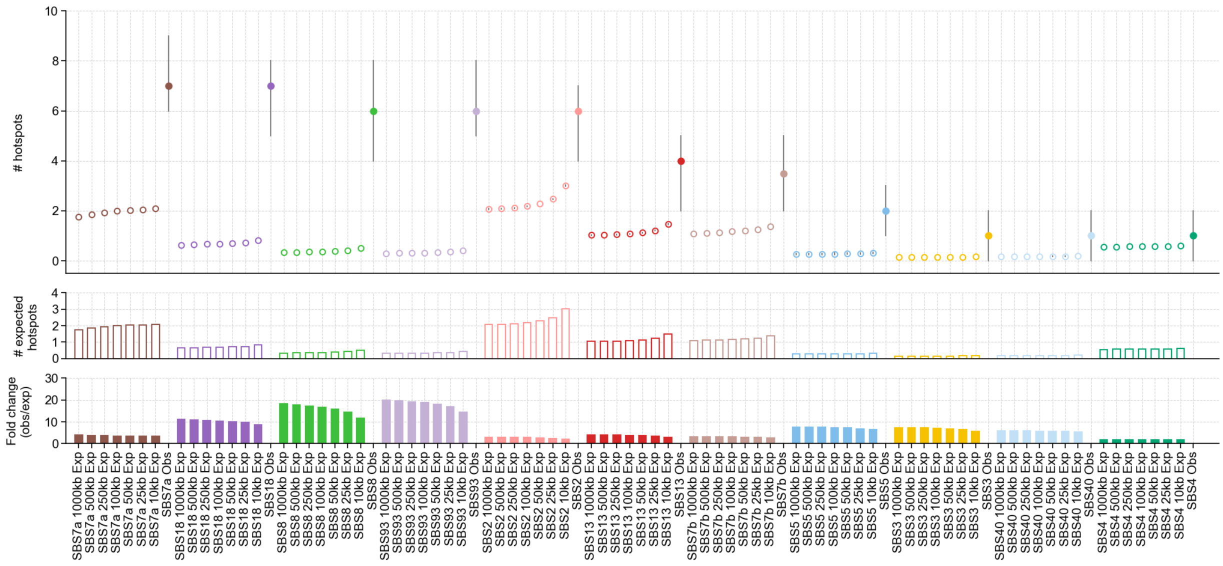


**Supp. Fig. 26. Expected hotspot propensity using submegabase models.** Number of expected and observed hotspots across signatures together with the expected number of hotspots computed using models accounting for trinucleotide composition and large-scale mutation rate variability for different genomic bin sizes (1 Mbp to 10 Kbp) (top). The number of observed hotspots per model is shown (middle). Fold change of the number of observed hotspots versus expected hotspot for each model (bottom). Observed and expected hotspot propensities have been calculated using 30,000 mutations (300 mutations/sample and 100 samples).

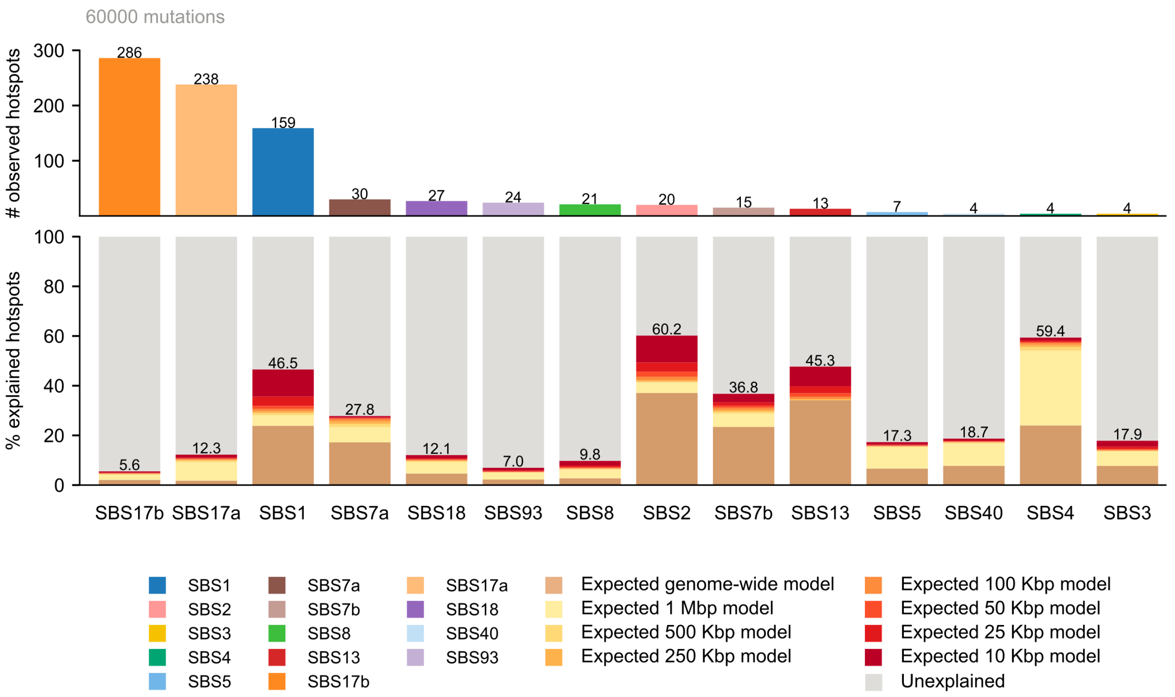


**Supp. Fig. 27. Expected hotspot propensity with larger subsampling.** Median number of hotspots per signature observed by subsampling 60,000 total mutations (600 mutations/sample, 100 samples) (top). Percentage of explained hotspots by the different models (bottom). Note that the number of cancer types in this experiment differs from that shown in Fig. 4 where a subsampling of 30,000 total mutations (300 mutations/sample, 100 samples) was used. Specifically, the following signature-cancer type pairs are not included in this experiment do to insufficient sampling size: SBS1-BLADDER_URI, SBS1-BRCA, SBS1-NSCLC, SBS5-BLADDER_URI, and SBS17a-COADREAD.

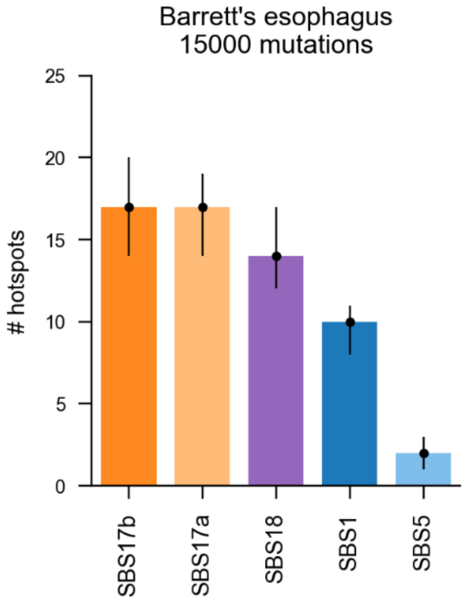


**Supp. Fig. 28. Hotspot propensity in Barrett's esophagus.** Hotspot propensity in Barrett's esophagus samples was computed using a total of 15,000 mutations per signature (50 samples and 300 mutations/sample).
