## Supplementary Notes for "Hotspot propensity across mutational processes"


##### Index


Supplementary Note 1: HotspotFinder

Supplementary Note 2: Mutational signatures extraction

Supplementary Note 3: Assignment of mutational signatures to mutations and hotspots

Supplementary Note 4: Comment on hotspot propensity

Supplementary Note 5: Theoretical models of hotspot propensity

Supplementary Notes References

### Supplementary Note 1
### HotspotFinder

HotspotFinder is an algorithm implemented in Python 3.6 to identify and annotate hotspots of somatic mutations across the hg38 human genome. HotspotFinder identifies recurrently mutated genomic positions (in two or more samples) across the genome. Then, it annotates hotspots with a set of genomic data to contribute to their interpretation.

HotspotFinder v1.0 freely available at: `bitbucket.org/bbglab/hotspotfinder`

#### 1. Input data

The minimum required HotspotFinder input consists of a file containing single nucleotide variants (SNVs), multiple nucleotide variants (MNVs) and/or small insertions and deletions (indels). The data to annotate hotspots is automatically downloaded within the package installation. Alternatively, users can provide their own paths to genomic annotations as described below.

**Mutations**

HotspotFinder's main input file is a TSV or CSV file containing SNVs, MNVs and short indels from a cancer cohort generated by whole genome (WGS), whole exome (WXS) or panel sequencing. In the current version, mutations are supported in the human genome build hg38. The mutations file requires the following information: chromosome, position, reference nucleotide(s), alternate nucleotide(s) and sample identifier. The header of the mutation file is 'CHROMOSOME', 'POSITION', 'REF', 'ALT', 'SAMPLE'.

By default, hotspots are identified using all samples in the mutations file. Alternatively, HotspotFinder can report hotspots independently in sub-groups or categories drawn from the input file. This feature requires adding an extra column with one of these headers: 'GROUP', 'GROUP_BY', 'COHORT', 'CANCER_TYPE', 'PLATFORM', and 'TYPE'. The new column name must be specified in the configuration file.

**Genomic annotations**

HotspotFinder incorporates genomic information to annotate hotspots and help in their interpretation. These annotations include: genome mappability, repeats and low complexity regions, population variants, immunoglobulin loci, and genomic elements (coding sequences from protein coding transcripts, splice sites, introns, 5' and 3' untranslated regions, promoters, non-coding RNAs, and pseudogenes). These files are automatically downloaded from bgdata package the first time that HotspotFinder is run. Users can also replace the automatic annotations with their own data by adding paths to the files within the configuration file:

- *Genome mappability.* Mappability was annotated from two different sources: regions of high mappability in the hg38 reference genome build and hg38 blacklisted regions of low mappability. In order to obtain high mappability regions, The GEnomic Multi-tool (GEM)[1] mappability software version 2013-04-06 was run on hg38 and those sequences with 100-mer pileup mappability equal or greater than 0.9 were kept. Blacklisted mappability regions were obtained from ENCODE[2] Unified GRCh38 Blacklist (downloaded from encodeproject.org/files/ENCFF356LFX on 16-06-2020).

- *Population variants*. Population variants were obtained from gnomAD[3] (downloaded from gnomad.broadinstitute.org on 25-06-2020) version 3.0. We kept variants with an allele frequency equal or greater than 1% (polymorphisms).

- *Repeats and low complexity regions.* Repeats across hg38 genome computed by RepeatMasker (RepeatMasker open-4.0.5, Repeat Library 20140131, hg38, Dec 2013) (developed by Smit, Hubley, Green at repeatmasker.org) were downloaded on 19-11-2020 at repeatmasker.org/species/hg.html

- *Immunoglobulin loci.* Immunoglobulin variable chain and T-cell receptor genes in hg38 build were obtained from Gencode[4] v35 comprehensive gene annotation file at ftp.ebi.ac.uk/pub/databases/gencode/Gencode_human/release_35/gencode.v35.annotation.gtf.gz (downloaded on 30-08-2020).

- *Genomic elements*. The following genomic elements from hg38 build were defined from Gencode[4] v35 comprehensive gene annotation file at ftp.ebi.ac.uk/pub/databases/gencode/Gencode_human/release_35/gencode.v35.annotation.gtf.gz (downloaded on 30-08-2020). Annotations were generated for all transcripts per gene, including the following genomic element categories.
    - Coding sequences (CDS): protein-coding sequences within protein-coding transcripts.
    - Splice sites: first and last 25 bp sequences within introns of protein-coding transcripts.
    - 5'UTRs: 5' untranslated sequences from protein-coding transcripts.
    - 3'UTRs: 3' untranslated sequences from protein-coding transcripts.
    - Proximal promoters: sequences 200 bp 5' and 200 bp 3' of transcription start sites (inclusive) of protein coding transcripts. These sequences may overlap 5'UTRs or CDS sequences.
    - Distal promoters: sequences 2,000 bp 5' of transcription start sites of protein coding transcripts.
    - Introns: intronic sequences of protein-coding transcripts.
    - lncRNA exons: exonic sequences of long non-coding RNA genes.
    - lncRNA splice sites: first and last 25 bp sequences within introns of long non-coding RNA genes.
    - lncRNA proximal promoters: sequences 200 bp 5' and 200 bp 3' of transcription start sites (inclusive) of long non-coding RNA genes.
    - lncRNA distal promoters: sequences 2,000 bp 5' of transcription start sites of long non-coding RNA genes.
    - lncRNA introns: intronic sequences of long non-coding RNAs.

- sncRNA: small non-coding RNAs of different types, including mt_rRNA, mt_tRNA, miRNA, misc_RNA, rRNA, scRNA, snRNA, and snoRNA. A complete description of each category is available at gencodegenes.org.
- Pseudogenes: all pseudogenes annotated at gencodegenes.org.

Considering that genomic elements may overlap each other, we set a hierarchical system of annotation priorities based on the potential functional relevance to protein-coding sequences. From highest to lowest priority, the overlap hierarchy was: CDS, splice sites, UTRs, promoters, introns, and lncRNAs (including their exons, introns and promoters) other ncRNAs and pseudogenes. No particular priority was assigned to the elements within the last three categories (their annotations are assigned equal priority and thus are allowed to overlap each other).

**Configuration file**

HotspotFinder allows a wide range of configurations to run. These can be specified through `hotspot.conf` file.

#### 2. Output data

HotspotFinder generates two output files: `results.tsv` and `warningpositions.txt`

**HotspotFinder hotspots**

The main output of HotspotFinder is saved in the TSV file `results.tsv`. Each row in the results file corresponds to a hotspot. Hotspots are independently identified in each of the four mutation types: single nucleotide variants (SNVs), multi-nucleotide variants (MNVs), short insertions and short deletions.

By default, `results.tsv` file contains the following columns:

- CHROMOSOME: chromosome
- POSITION: position
- CHR_POS: chromosome and position
- HOTSPOT_ID: unique hotspot identifier
- MUT_TYPE: mutation type
- COHORT: input cohort (file) name
- N_MUTATIONS: number of mutations in the hotspot
- N_MUTATED_SAMPLES: number of mutated samples in the hotspot
- FRAC_MUTATED_SAMPLES: fraction of mutated samples in the cohort
- REF: reference nucleotide
- ALT: alternate nucleotide(s)
- ALT_COUNTS: number of alternate nucleotide(s)
- FRAC_ALT: proportion of alternates among total alternates
- CONTEXT_3: trinucleotide context centred at the hotspot
- CONTEXT_5: pentanucleotide context centred at the hotspot
- N_COHORT_SAMPLES: number of samples in the cohort
- N_COHORT_MUTATIONS_TOTAL: number of total mutations in the cohort

**-** N_COHORT_MUTATIONS_SNV: number of single base substitutions in the cohort
**-** N_COHORT_MUTATIONS_MNV: number of multi-nucleotide substitutions in the cohort
**-** N_COHORT_MUTATIONS_INS: number of insertions in the cohort
**-** N_COHORT_MUTATIONS_DEL: number of deletions in the cohort
**-** MUTATED_SAMPLES: identifier of mutated samples
**-** MUTATED_SAMPLES_ALTS: identifier and corresponding alternate of mutated samples
**-** OVERLAP_WARNING_POSITION: True if the hotspot overlaps a warning position

If HotspotFinder is run with annotations, additional columns are included in `results.tsv` file:

**-** GENOMIC_ELEMENT: overlapping genomic element(s)
**-** SYMBOL: symbols off overlapping genomic element(s)
**-** GENE_ID: Ensembl gene identifiers of overlapping genomic elements
**-** TRANSCRIPT_ID: Ensembl transcript identifiers of overlapping genomic elements
**-** GENOMIC_REGION: class of overlapping genomic element(s)
**-** GENOMIC_REGION_PRIORITY: genomic element class with highest priority
**-** CODING_NONCODING: overlapping coding or non-coding genomic elements
**-** MAPPABILITY: high or low according to GEM mappability regions overlap
**-** MAPPABILITY_BLACKLIST: pass if no overlap with blacklisted mappability regions
**-** VARIATION_AF: overlapping population variants
**-** HOTSPOTFINDER_FILTERS: flag if hotspot is mappable and does not overlap blacklisted mappability regions or population variants.
**-** REPEATS: overlapping repeats
**-** REPEATS_OVERLAP: flag if hotspot overlaps repeats
**-** IG_TR: overlapping immunoglobulin regions
**-** IG_TR_OVERLAP: flag if hotspot overlaps immunoglobulin regions

**HotspotFinder warning positions**
During the process of mutation parsing some unexpected data can be observed and are reported in a separate TSV file, `warningpositions.txt`, with the following header: 'CHROMOSOME', 'POSITION', 'REF', 'ALT', 'SAMPLE', 'WARNING', 'SKIP' (check “Mutation parsing” for further information).

#### 3. Implementation

**Mutation parsing**
Parsing of somatic mutations is performed in two steps as explained below:

In the first step, the package **bgparsers** reads the mutation file and categorises mutations into 3 mutation types: SNVs, MNVs and indels. Indels are further classified into simple insertions (e.g., G>GA or G>GAAA) or simple deletions (e.g, GT>G, GTG>G). Complex indels that are a mixture of both previous categories (e.g., GTG>GA, GTG>GAAA) are skipped. At this level, some filters are applied: 1) mutations falling outside of autosomal or sexual chromosomes are skipped; 2) mutations having equal reference and alternate nucleotides are removed; 3) mutations with annotated reference nucleotide(s) that do not correspond to the reference genome build (in this case, only for SNVs, MNVs and deletions) are discarded; 4) mutations that contain an unknown nucleotide (a nucleotide not corresponding to A, C, G, T) are removed; 5) mutations with an unknown nucleotide in their pentamer sequence context, according to their start position, are skipped from the analysis.

In the second step, the algorithm checks how many alternates a given sample (patient) has for each mutated position of the genome. It is expected that homozygous sample for a locus will have a single alternate (e.g., chr1:23798511_G>A), whereas heterozygous samples can have 2 (e.g., chr1:23798511_G>A and chr1:23798511_G>T ). However, other scenarios may arise such as homozygous samples annotated with two equal alternates separately in the input file,heterozygous samples bearing3 different alternates in the case of SNVs or more than 3 in the case of other mutations. Given that the algorithm is agnostic of how the mutation calling has been performed and additional information that might be of interest to interpret these situations such as tumour ploidy or copy number alterations, it parses mutations using the following criteria:

a) If there is only one alternate count for a given patient, genome position and mutation type, this mutation is kept for the analysis.

b) If there is more than one alternate count for a given patient, genome position and mutation type, three situations can occur and are categorised in a warning level as follows: 1) a unique alternate is found more than one time (e.g., chr1:23798511_G>A and chr1:23798511_G>A); 2) there are two different alternates (e.g., chr1:23798511_G>A and chr1:23798511_G>C); 3) there are three or more different alternates (e.g., chr1:23798511_G>A, chr1:23798511_G>C and chr1:23798511_G>T). Given that mutations in scenario 1 and 2 are likely to reflect *bona fide* biological information (homozygous or heterozygous mutants for a locus, respectively) all mutations are kept for analysis. This implies that the number of mutations and the number of mutated samples in a given position in the genome will not match. Mutations in scenario 3 are discarded because their biological interpretation is unclear considering the data available within HotspotFinder. Any mutation falling in these three categories is reported in the output file `warningpositions.txt`. This information is also highlighted in the 'OVERLAP_WARNING_POSITION' column in the hotspots output file `results.tsv`.

**Hotspots identification**

Hotspots are defined as single genomic positions bearing 2 or more somatic mutations from 2 or more different samples. Hotspots are independently identified for 4 types of mutations: single nucleotide variants (SNVs), multi-nucleotide variants (MNVs), short insertions and short deletions.

By default, mutations are merged together according to their reference start position, irrespective of the length of the reference or alternate nucleotides in the case of MNVs, insertions and deletions. For instance, the variants chr1:123456_A>T mutation, chr1:123456_A>T, and chr1:123456_A>C mutation will be identified as a hotspot of SNVs in the position chr1:123456 with alternates T,T,A. In the case of MNVs, variants are merged into hotspots if they start in the same genomic position. For example: if the variants chr17:7571726_GTG>CCC, chr17:7571726_GT>CA, chr17:7571726_GTG>CCA appear in three independent samples, they will be merged into a hotspot. In the case of insertions, chr1:123456_A>AT, chr1:123456_A>ATT and chr1:123456_A>AGG variants in three independent patients will result in an insertion hotspot at chr1:123456 with alternates T,TT,GG. Similarly, in the case of deletions, chr1:123456_AG>A, chr1:123456_AGT>A and chr1:123456_AGTA>A mutation will result in a hotspot at chr1:123456 with reference sequences G, GT and GTA.

In the current version, it is also possible to restrict the identification of hotspots to those mutations sharing the same nucleotide change in the case of SNVs and MNVs, equal inserted sequence in the case of insertions or equal deleted sequence in the case of deletions. This behaviour can be specified in the configuration file. All results shown within the current manuscript are based on alternate-specific hotspots. This option was chosen to increase the signal-to-noise ratio to further analyse the mutational processes causing hotspots.

**Hotspots annotation**

Once hotspots are identified, by default HotspotFinder intersects their coordinates with genomic annotations. One can disable this option in the configuration file and the running time will be shortened. Hotspots that do not overlap high mappability regions or do overlap mappability blacklisted regions are considered artefact hotspots and are flagged accordingly. Similarly, hotspots that overlap a polymorphic site, irrespective of the mutation type or alternate, are considered artefacts of germline filtering. Importantly, these hotspots are kept in the output file `results.tsv`. By default, hotspots that do not overlap genomic elements are removed from the output file. This behaviour can be changed in the configuration file.

### Supplementary Note 2
### Extraction of mutational signatures

##### Mutational signatures extraction using total SNVs

We extracted *de novo* trinucleotide-context single base substitution signatures (96-mutation types using pyrimidines as reference) using SigProfiler framework[5–7] over the 31 cancer types, excluding Pancancer, with at least 30 samples and 100,000 total SNVs (hotspot and non-hotspot mutations): ALL, BILIARY_TRACT, BLADDER_URI, BONE_SOFT_TISSUE, BOWEL, BRAIN, BRCA, CERVIX, COADREAD, ES, ESOPHA_STOMACH, GBM, HEAD_NECK, KIDNEY, LIVER, LNM, LUNG, MBL, NHL, NSCLC, OV, PANCREAS, PLMESO, PROSTATE, SBNET, SCLC, SKCM, SKIN, ST, THYROID, UCEC.

For each cancer type, the GRCh38 96-mutational catalogues of total SNVs were calculated using SigProfilerMatrixGenerator[7] version 1.1.26. Mutational signatures were extracted with SigProfilerExtractor[6] version 1.1.0 testing the optimal solution for minimum of 1 and a maximum of 20 signatures (minimum_signatures, maximum_signatures parameters) with 1,024 NMF replicates (nmf_replicates parameter) and GRCh38 reference genome. All other NMF parameters were set as default. Solutions with average signature stability above 0.8, minimum signature stability above 0.2 and mean cosine distances (error) below 0.15 were pre-selected for manual review. We then assessed the cosine similarity between them and COSMIC v3.2 reference signatures by computing the complementary of the distance among each pair of vectors of mutational probabilities using scipy.spatial.distance:

$$cosine\ similarity = 1 - cosine\ distance$$

*De novo* signatures were assigned to the best matching (largest cosine similarity) COSMIC reference signature. After reviewing the assigned signature sets among all solutions, the best solution was selected based on the biological plausibility of the extracted signatures. When different solutions showed equally plausible assignments by cosine similarity, the solution with smallest mean cosine distances and largest cosine similarity was selected (Supp. Note 2 Fig. 1).

In order to compare signatures activity across cancer types, the best solution was finally decomposed into COSMIC v3.2 reference signatures using the default decomposition function within SigProfilerExtractor (refit_denovo_signatures=True). Signatures that could not be decomposed, and therefore were not equivalent among cancer types, were discarded from further analysis. Cancer types whose signature extractions had no solution matching the above-mentioned quality criteria were discarded for further analysis, including BONE_SOFT_TISSUE, LNM, NBL, and NHL. Likewise, the following known artefact signatures[8] were excluded from the analysis: SBS27, SBS43, SBS45, SBS46, SBS47, SBS48, SBS49, SBS50, SBS51, SBS52, SBS53, SBS54, SBS55, SBS56, SBS57, SBS58, SBS59, SBS60.

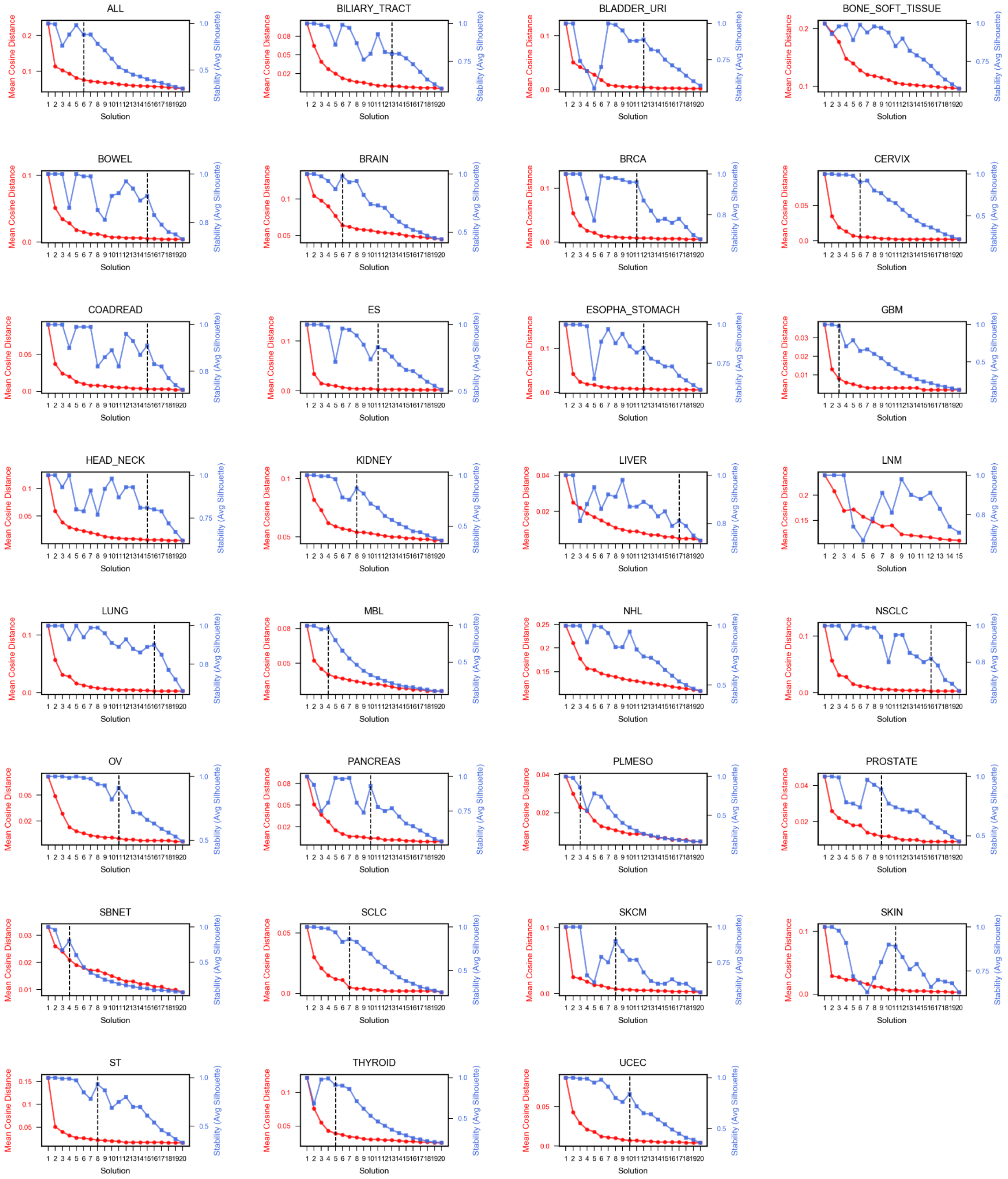


**Supp. Note 2 Fig. 1. Selection plots for extracted signatures using total SNVs**. Vertical dashed lines show the best extracted solution that fulfilled the quality criteria explained above.

**Extractions of SBS17a and SBS17b**

SBS17a and SBS17b are two signatures with high hotspot propensity in our analysis, both of which are of unknown aetiology[8]. SBS17a is characterised by frequent CpTpN>C transitions, while SBS17b preferentially shows NpTpT>G transversions. CpTpT trinucleotides are the most frequently mutated in both of them[8]. First identified as a joint signature, SBS17 [9], these signatures have been found in primary tumours, notably in esophageal cancers[10], and, in the case of SBS17b, also in metastases of patients exposed to 5-fluorouracil (5-FU) or capecitabine during the treatment or their primary tumours[11,12].

In our study, SBS17a and SBS17b were found across esophageal-stomach and colorectal cancers (Supp. Fig. 12). Three *de novo* signatures contributed to SBS17a and SBS17b in each cancer type (Supp. Note 2 Fig. 2). Among both sets, the signatures contributing to more mutations (SBS96-A in esophageal-stomach and SBS96-F in colorectal cancers) showed both the T>G and T>C components of SBS17b and SBS17a, with predominance of T>G transversions that explain their higher cosine similarity to COSMIC v3.2 SBS17b). Additionally, two signatures (SBS96-B in esophageal-stomach and SBS96-I in colorectal cancers) were clearly identified as SBS17b (cosine similarity of 1 and 0.97 to SBS17b, respectively). Among the 194 esophageal-stomach tumours with SBS96-B activity (defined by a signature contribution of at least 5% of mutations of the total mutation burden), 112 were metastatic and 37 (33%) of these had been exposed to either 5-FU or capecitabine. In colorectal cancers, 146/150 tumours bearing SBS96-B were metastases, the majority of which (n=123, 84%) showed 5-FU or capecitabine exposure. This is consistent with previous studies on SBS17b [11,12]. Altogether, these data explain the larger mutation burden of SBS17b compared to SBS17a in our dataset.

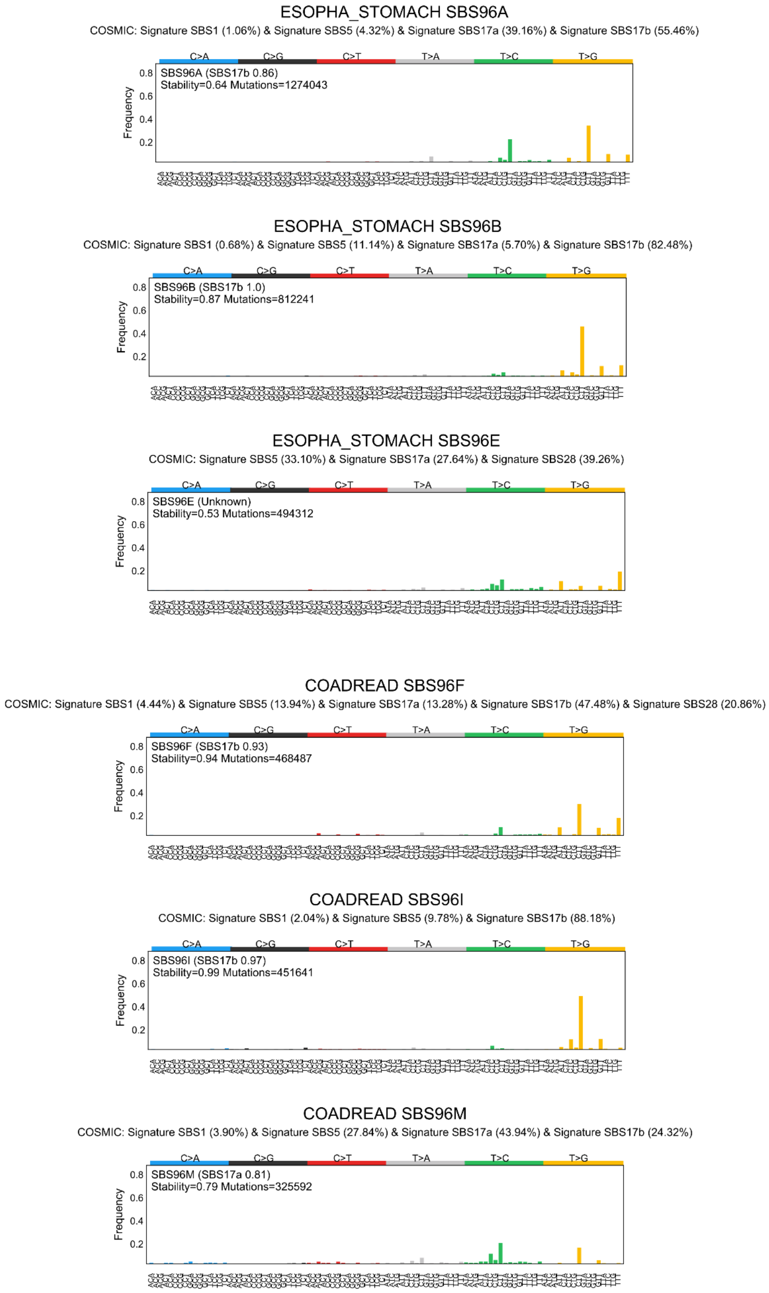


**Supp. Note 2 Fig. 2. Extracted signatures contributing to SBS17a and SBS17b.** *De novo* extracted signatures partially decomposed into COSMIC v3.2 SBS17a and SBS17b signatures. The signature decomposition attribution to these and other signatures is shown.

#### Mutational signatures extraction using SNVs inside hotspots

A second extraction of *de novo* trinucleotide-context single base substitution signatures (96-mutation types using pyrimidines as reference) was carried out using exclusively the set of SNVs inside hotspots for the top 4 cancer types with largest number of hotspots: COADREAD; ESOPHA_STOMACH (instead of ES), NSCLC, SKCM. GRCh38 96-mutational catalogues of SNVs overlapping hotspots were calculated using SigProfilerMatrixGenerator[7] version 1.1.26. Mutational signatures were extracted with SigProfilerExtractor[6] version 1.1.0 running for a minimum of 1 and a maximum of 15 signatures (minimum_signatures, maximum_signatures parameters) with 1,024 NMF replicates (nmf_replicates parameters). All other NMF parameters were set as default. The best solution was selected following the same approach as in the total SNVs extraction (Supp. Note 2 Fig. 3). The resulting *de novo* signatures inside hotspots listing the COSMIC v3.2 signature with largest cosine similarity above 0.8 are shown below (Supp. Note 2 Fig. 4)

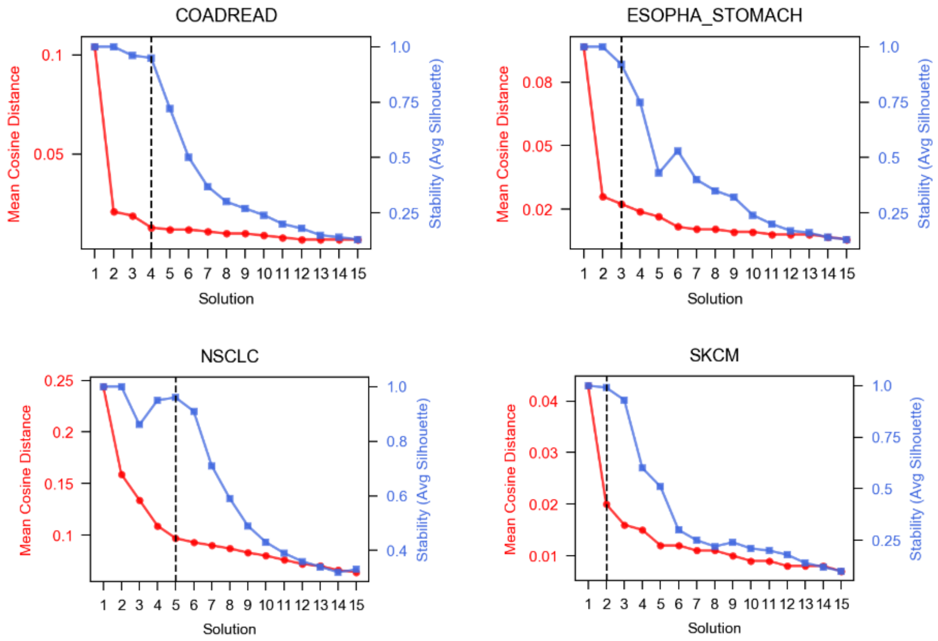


**Supp. Note 2 Fig. 3. Selection plots for extracted signatures using SNVs inside hotspots**. Vertical dashed lines show the best extracted solution that fulfilled the quality criteria explained above.

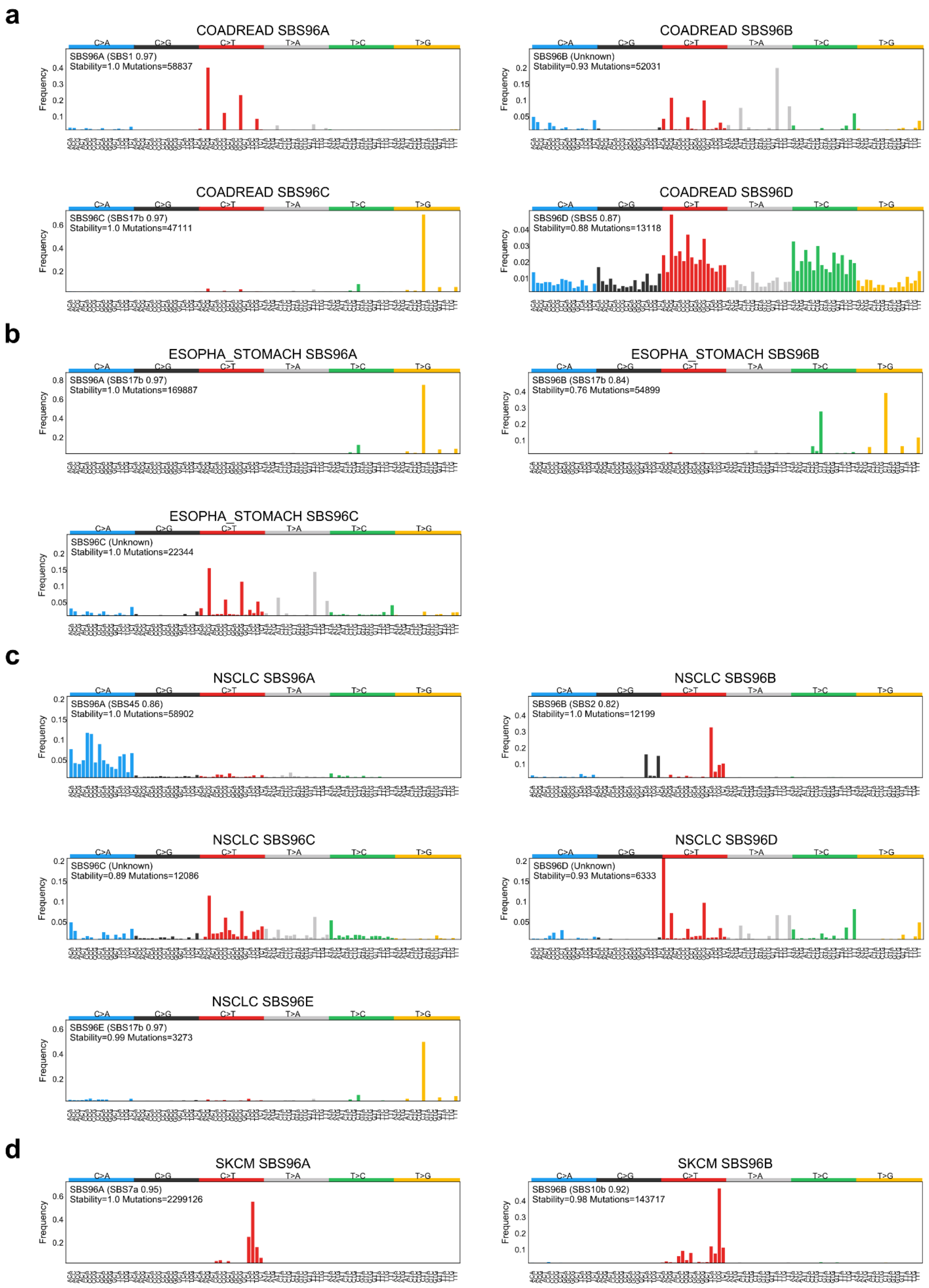


**Supp. Note 2 Fig. 4. Extracted mutational signatures from SNVs inside hotspots**. Signature profiles showing the *de novo* extracted signatures for **a**) colorectal cancers, **b**) esophagus-stomach cancers, **c**) non-small cell lung cancers and **d**) skin melanomas. The COSMIC signature with largest cosine similarity above 0.8 to the profile is stated, otherwise it is labelled as unknown.

### Supplementary Note 3
### Assignment of mutational signatures to mutations and hotspots

##### Cancer datasets

As part of our analysis in cancer samples, we assigned or attributed mutations and hotspots to the mutational signature that most likely generated them. This attribution was carried out as follows.

As detailed in Supplementary Note 2, we first extracted *de novo* mutational signatures from cancer types using SigProfilerExtractor[6]. In order to obtain an homogeneous set of reference signatures and their exposures across cancer types, we then used SigProfilerExtractor decomposition of *de novo* signatures into the COSMIC v3.2 signatures reference set. Specifically, SigProfilerExtractor uses a nonnegative least squares (NNLS) algorithm to identify the coefficients (contributions) of COSMIC signatures that best recapitulate each of the *de novo* signatures in a cancer type (for details see [6]). Once the contribution of a set of COSMIC signatures has been obtained for each *de novo* signature, an additional NNLS algorithm is applied to identify the activity of COSMIC signatures to each sample in the dataset. Then, the probability of a particular mutation within a sample (attributed to a trinucleotide-context substitution, e.g., A[C>A]A) to arise from a COSMIC decomposed signature is calculated based on the mutation trinucleotide-context, the sample-specific signature exposures and the signature profile (mutational probabilities per trinucleotide-context).

Specifically, the likelihood that signature $S$ contributes to a mutation observed in triplet context $t$ and sample $k$ is computed as follows[13]:

$$P(S;t,k)=\frac{w_{km}S(t)}{\sum_{i=1}^{M} w_{ki}S_i(t)}$$

where the vector of total exposures of the sample is given by the following weighted sum:

$$e_k=\sum_{i=1}^{M} w_{ik}S_i$$

$S(t)$ denotes the frequency of context $t$ in the profile of signature $S$ and $i$ indexes all the mutational signatures contributing mutations to the cohort.

This results in a vector of mutational probabilities per sample and trinucleotide context across the active signatures in the cancer type. From this matrix of vectors, we then assign individual mutations within a sample to a mutational signature using maximum likelihood, if the probability of maximum likelihood signature is above a probability of 0.5 (Supp. Note 3 Fig. 1). The mutational profiles resulting from mutations assigned by maximum likelihood to a signature showed a high similarity to the corresponding COSMIC signature profile (median cosine similarity = 0.98, range(0.63, 1); Supp. Note 3 Fig. 2).

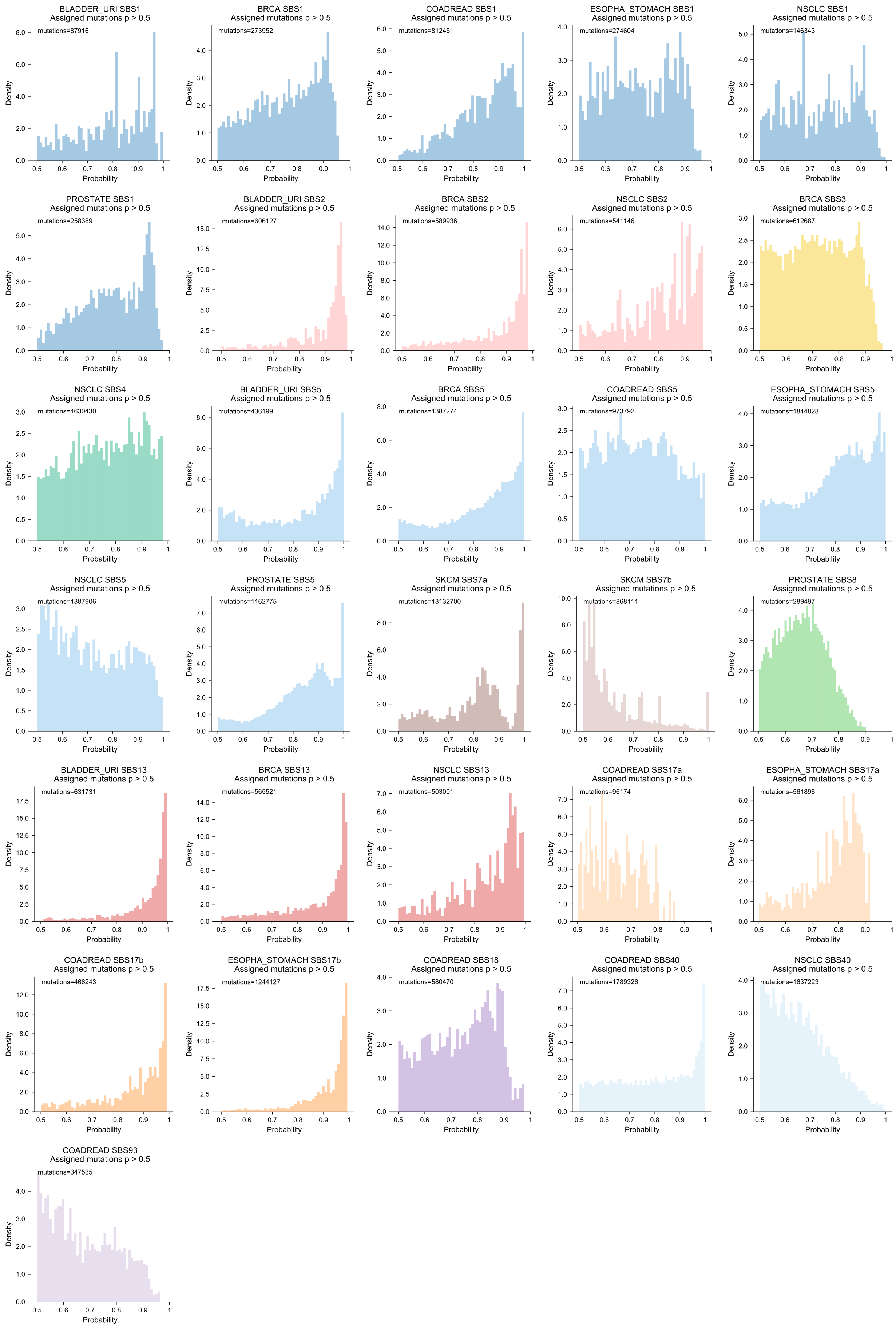
BLADDER_URI SBS1
Assigned mutations p > 0.5
mutations=87916
BRCA SBS1
Assigned mutations p > 0.5
mutations=273952
COADREAD SBS1
Assigned mutations p > 0.5
mutations=812451
ESOPHA_STOMACH SBS1
Assigned mutations p > 0.5
mutations=274604
NSCLC SBS1
Assigned mutations p > 0.5
mutations=146343
PROSTATE SBS1
Assigned mutations p > 0.5
mutations=258389
BLADDER_URI SBS2
Assigned mutations p > 0.5
mutations=606127
BRCA SBS2
Assigned mutations p > 0.5
mutations=589936
NSCLC SBS2
Assigned mutations p > 0.5
mutations=541146
BRCA SBS3
Assigned mutations p > 0.5
mutations=612687
NSCLC SBS4
Assigned mutations p > 0.5
mutations=4630430
BLADDER_URI SBS5
Assigned mutations p > 0.5
mutations=436199
BRCA SBS5
Assigned mutations p > 0.5
mutations=1387274
COADREAD SBS5
Assigned mutations p > 0.5
mutations=973792
ESOPHA_STOMACH SBS5
Assigned mutations p > 0.5
mutations=1844828
NSCLC SBS5
Assigned mutations p > 0.5
mutations=1387906
PROSTATE SBS5
Assigned mutations p > 0.5
mutations=1162775
SKCM SBS7a
Assigned mutations p > 0.5
mutations=13132700
SKCM SBS7b
Assigned mutations p > 0.5
mutations=868111
PROSTATE SBS8
Assigned mutations p > 0.5
mutations=289497
BLADDER_URI SBS13
Assigned mutations p > 0.5
mutations=631731
BRCA SBS13
Assigned mutations p > 0.5
mutations=565521
NSCLC SBS13
Assigned mutations p > 0.5
mutations=503001
COADREAD SBS17a
Assigned mutations p > 0.5
mutations=96174
ESOPHA_STOMACH SBS17a
Assigned mutations p > 0.5
mutations=561896
COADREAD SBS17b
Assigned mutations p > 0.5
mutations=466243
ESOPHA_STOMACH SBS17b
Assigned mutations p > 0.5
mutations=1244127
COADREAD SBS18
Assigned mutations p > 0.5
mutations=580470
COADREAD SBS40
Assigned mutations p > 0.5
mutations=1789326
NSCLC SBS40
Assigned mutations p > 0.5
mutations=1637223
COADREAD SBS93
Assigned mutations p > 0.5
mutations=347535
Density
Probability

**Supp. Note 3 Fig. 1. Distribution of mutational probabilities across mutations attributed to a signature**. Histograms showing the distribution of mutational probabilities for mutations attributed to COSMIC mutational signatures by maximum likelihood (p > 0.5) across cancer types.

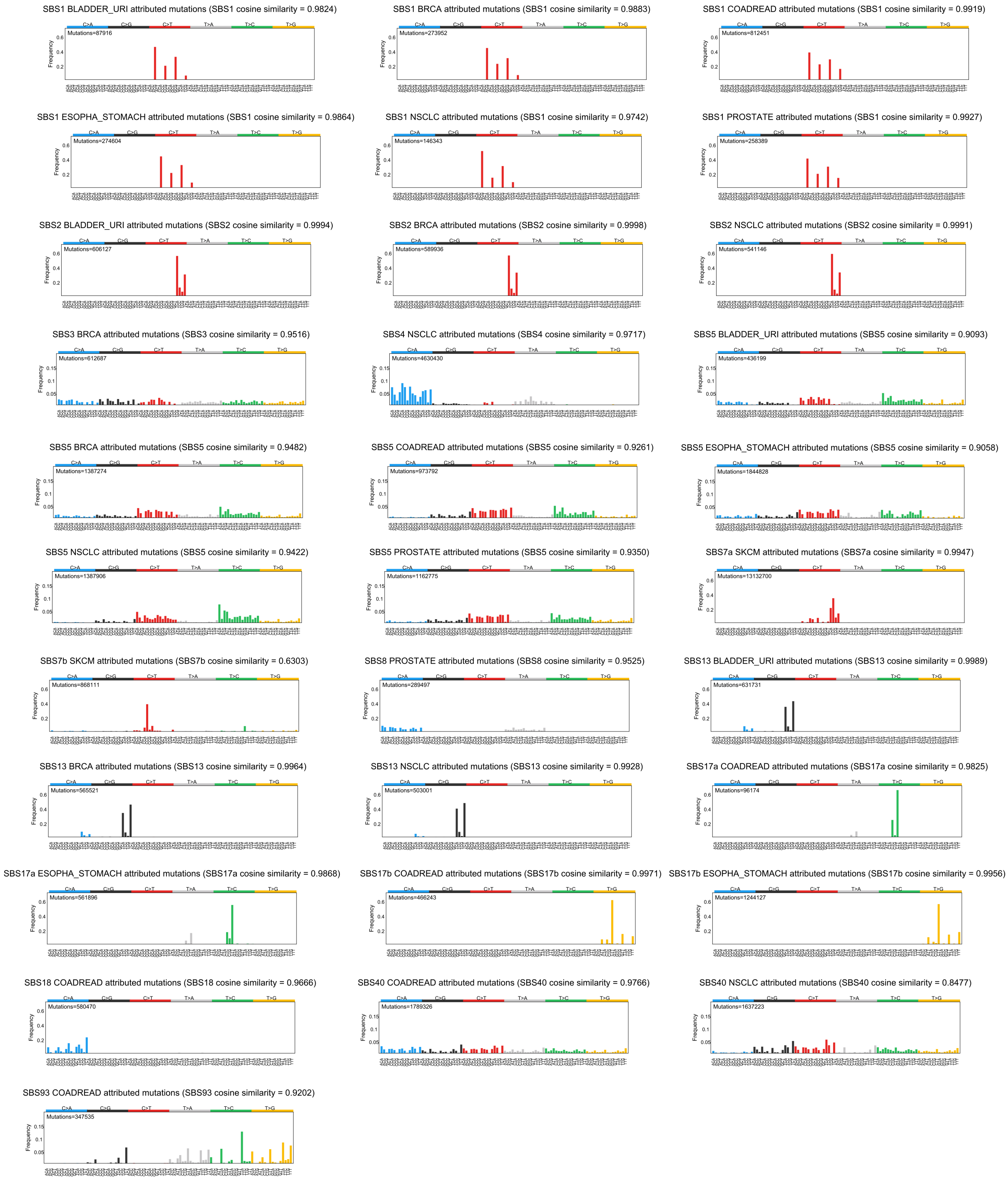


**Supp. Note 3 Fig. 2. Mutational profiles of mutations attributed to signatures across cancers**. Mutational profiles for mutations attributed to COSMIC mutational signatures by maximum likelihood ($p > 0.5$) across cancer types. The cosine similarity to the corresponding COSMIC signature is shown on top.

#### Normal tissue datasets

Similarly as in the cancer datasets, we attributed mutations in colonic crypts[14], Barrett's esophagus[15] and germline datasets[16–21] to mutational signatures active in the respective tissue by maximum likelihood with a probability above 0.5. The mutational profiles of the resulting attributed mutations per signature showed a high cosine similarity (>0.9) to their reference COSMIC signature (Supp. Note 3 Fig. 3).

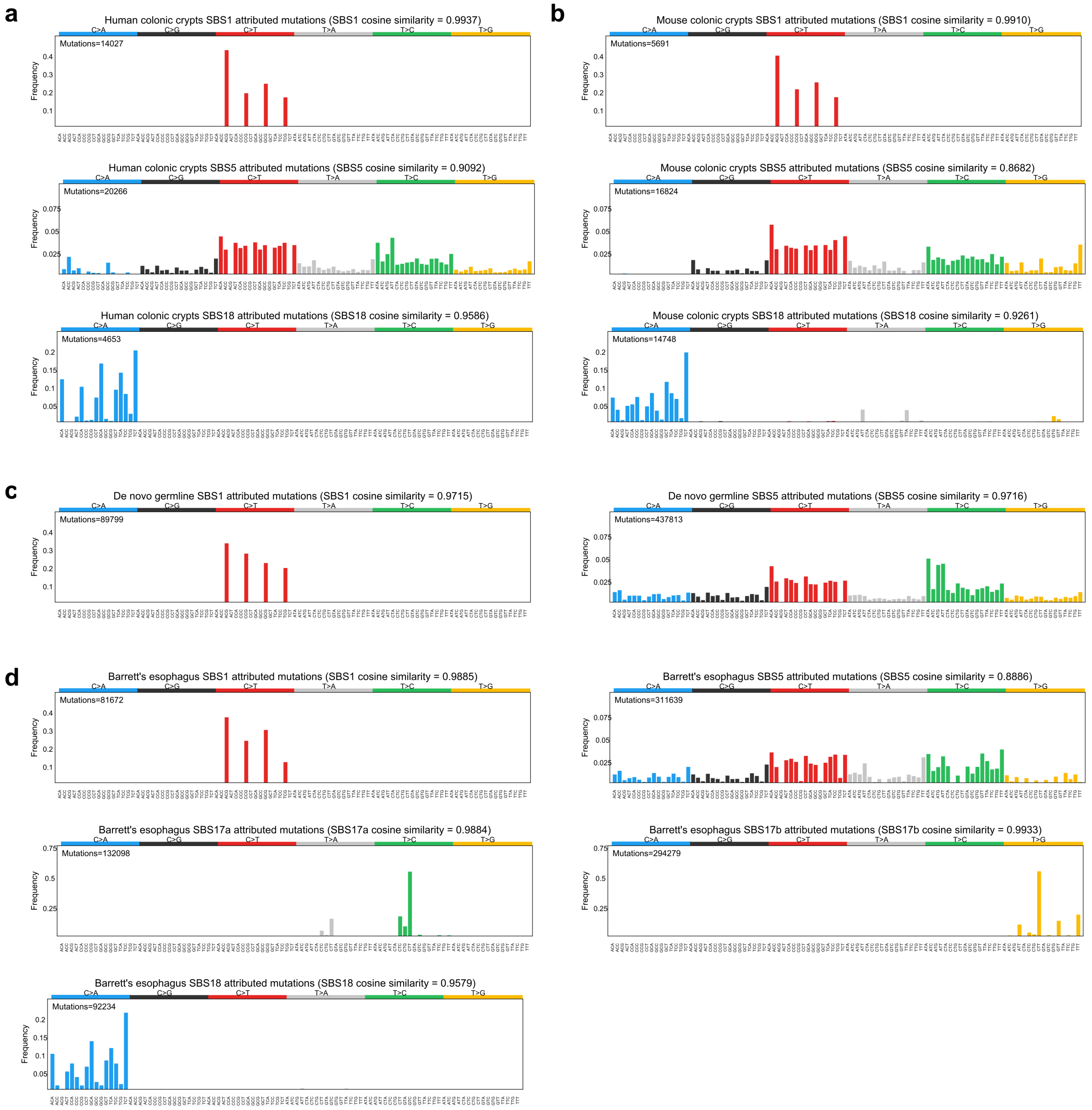


**Supp. Note 3 Fig. 3. Mutational profiles of mutations attributed to signatures in normal tissues.** Mutational profiles for mutations attributed to COSMIC mutational signatures by maximum likelihood (p >

0.5) in **a**) human colonic crypts, **b**) mouse colonic crypts, **c**) de novo germline mutations and **d**) Barrett's esophagus. The cosine similarity to the corresponding COSMIC signature in the matched reference genome is shown on top.

### Supplementary Note 4
### Comment on hotspot propensity

We used two metrics to calculate the propensity of mutational processes to form hotspots. First, we calculated the number of newly observed hotspots through subsampling of mutations contributed by a signature under equal mutation and sample sizes across a cohort. Briefly, for each cancer type, we assigned mutations to the signature with maximum probability of having generated them. To increase the accuracy of this allocation, we only attributed a mutation to a signature if its probability was above 0.5. Then, we randomly selected (subsampled) 1,000 sets of N mutations (e.g., 30,000 mutations) from each signature across 100 patients (all of which contributed with equal mutation burden) and computed the number of hotspots observed in each subsample. Under equal mutation burden, we expect to observe more hotspots contributed by signatures with higher propensity to form them. Given that we leverage separate sets of mutations per signature, this estimate of a signature's hotspot propensity is independent of all other signatures in the cancer type. This is particularly important when evaluating mutational processes active in the same samples with overlapping trinucleotide preference, which may interact in the formation of hotspots. For example, SBS17a and SBS17b, which are active in esophageal and colorectal cancers, partially share their target trinucleotides (specially the CpTpT trinucleotide[8]), but show different mutation burden in our dataset. By separating the sets of mutations attributed to each of them (Supp. Note 4 Fig. 1), their hotspot propensities can be assessed independently from each other. This metric, however, assumes that mutations contributed by a signature are equally distributed across the tumours in a cohort. Since this is not necessarily true, we developed a metric that is orthogonal to the one described above.

In this second metric, we leveraged the total number of observed hotspots and mutations (classified as being outside or inside hotspots) contributed by all signatures and computed the fold change of the activity of each of them inside and outside hotspot mutations (Supp. Fig. 18). We reasoned that signatures with higher propensity to form hotspots would be more frequently found in the set of mutations inside hotspots than in the set of mutations outside hotspots (Supp. Fig. 17). Therefore, if we computed the ratio (fold change) of a signature activity inside versus outside hotspots, those with propensity to form hotspots would show a fold change larger than 1. Conversely, signatures with no particular propensity for hotspots formation would show similar frequencies in both sets of mutations and consequently a fold change of 1. The fold change for the top three signatures with highest hotspot propensity based on subsampling, SBS17b, SBS17a and SBS1, was larger than 1 across a wide variety of cancers (Supp. Fig. 18), thus validating our first observations. However, this measure showed some caveats.

Although applicable to most signature-cancer type pairs in our dataset without the need of subsampling samples and mutations, the fold change is computed using hotspots that have been contributed by all signatures present in the cancer type. That is, the hotspots according to which mutations are classified as inside or outside are assumed to be generated by a combination of mutational signatures in the cancer type, given the trinucleotide context and the signatures activities in the sample. Each mutation is treated as a vector of probabilities of the mutation arising from the catalogue of signatures in the cancer type. Then, in order to compute the frequency inside hotspots of a given signature in a cancer type, the vector of mutational probabilities across all mutations inside hotspots is normalised to 1. Likewise, the same step is carried out for outside mutations. This allows the comparison of signature frequencies inside and outside hotspots (otherwise the ratio of the sum of probabilities would be smaller than 1, assuming the number of mutations outside hotspots outnumbers that of mutations inside). The fact that the fold change is computed using hotspots contributed by different signatures poses a limitation to accurately identifying the hotspot propensity of different signatures with similar propensities but different activities in a cancer type.

As introduced before, a clear example of this are SBS17a and SBS17b in esophageal-stomach and colorectal cancers. SBS17b showed bigger hotspot fold change than SBS17a (median 3.1 and 1.2 fold change across cancer types, respectively). However, when we calculated hotspot propensity through subsampling of mutations independently assigned to any of both signatures by maximum likelihood (Supp. Note 4 Fig. 1), SBS17a and SBS17b showed very similar high propensity to form hotspots, generating, respectively, 79 and 73 hotspots across 30,000 mutations (Fig. 3f). The fact that SBS17b contributed more mutations than SBS17a in both cancer types and that both signatures target the same trinucleotides, particularly CpTpT sites, increases the probability of identifying SBS17b inside hotspots in detriment of SBS17a when the fold change enrichments are computed. For all these reasons, we chose the subsampling strategy as the main method to estimate hotspot propensity across signatures.

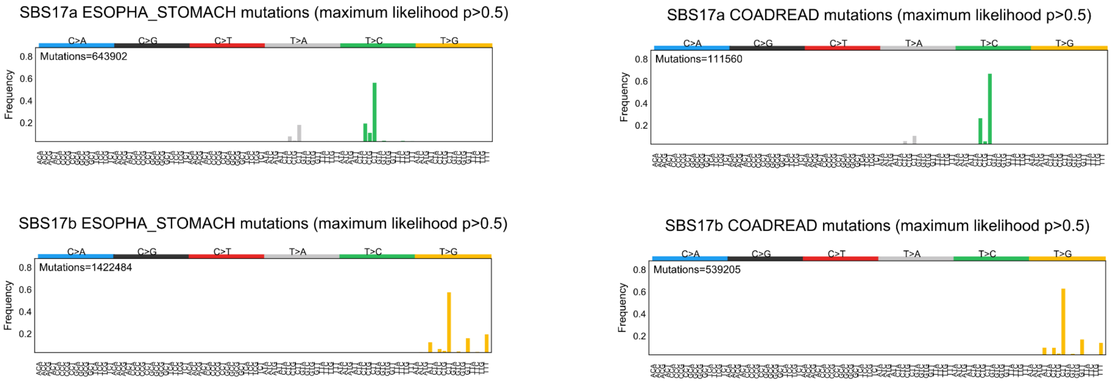


**Supp. Note 4 Fig. 1**. **Mutations assigned to SBS17a and SBS17b**. Frequencies of trinucleotide-based substitutions (either inside or outside hotspots) attributed to SBS17a and SBS17b by maximum likelihood above 0.5.

### Supplementary Note 5
### Theoretical models of hotspot propensity

**Overview**

Given a genomic chunk or region of DNA (e.g. the whole mappable reference genome or a specific covering of the genome in chunks of size 1Mbps, 500, 250, 100, 50, 25 or 10 Kbps) we provide a theoretical calculation of the expected number of hotspots produced in it by a synthetic cohort of samples that have identical mutation rates at that region. We name this expected number of hotspots “hotspot propensity” when referring to the expected count at genome-wide level, which can be obtained by adding up the expected number of hotspots across genomic chunks of the given size.

We assume that mutations are generated in accordance with the same mutational process, which in this analysis will be characterised by the relative frequency of each nucleotide substitution taking into account the flanking nucleotides (trinucleotide context). Furthermore, in our computation we will make the simplifying assumption that mutations are generated independently across samples and genomic positions. Using this fact, note that if we partition a region into disjoint subregions, the expected number of hotspots should be equal to the sum of expectations across subregions. In particular, we can compute the expected number of hotspots at a single genomic position, then sum the expected number of hotspots across all genomic positions of the region of interest. Note that in our framework all positions with the same reference trinucleotide will bear the same expected number of hotspots if we assume that the mutation rate is homogeneous across positions.

Formally, given a genomic region, an homogeneous mutation rate and a mutational profile operative in the region, we can compute how many hotspots are expected at each position, which will depend on the reference trinucleotide of the position, then sum the expectations across positions to get a total expectation for the whole region.

In order to compute the expected number of hotspots at a single position, we first need to transform the overall mutation rate of the region into mutation probabilities at single positions. In the following sections we will explain in detail how to derive this mutation probabilities at single positions (section 1), how with this information we can derive the expected number of hotspots at a given position when stacking a cohort with several samples (section 2) and how we use this information to derive expected number of hotspots genome-wide (section 3) in mappable genomic segments (section 4) and how we include DNA methylation covariates (section 5). Finally, in section 6 we showcase a stochastic generative model of hotspots counts that is compatible with our calculation of the expected hotspot rate.

**1. Mutation probability at a single position**

Consider a genomic region where a mutational process is operative with overall mutation rate $\mu$, meaning that the expected number of mutations in the region is $\mu$. Let $\underline{m}=\left(m_{t>a}\right)$ be the vector of normalised mutation rates – indexed by pairs of reference triplet $t$ and alternate allele $a$ – that defines the mutational process. Here the term "normalised" implies that the vector already reflects the relative mutation rates across trinucleotide contexts after correcting for the trinucleotide content of the region where the mutational frequencies were sampled from. In other words, it reflects the relative likelihood to observe some trinucleotide contexts compared to others. For a position with reference triplet $t$ we compute the probabilities to undergo either of the three possible alternate alleles (for simplicity denoted $A$, $B$ and $C$) as follows:

$$p_{t>A} = \frac{1}{M}\mu m_{t>A}, \quad p_{t>B} = \frac{1}{M}\mu m_{t>B}, \quad p_{t>C} = \frac{1}{M}\mu m_{t>C}, \quad p = p_{t>A} + p_{t>B} + p_{t>C}$$

$$\text{where} \quad M = \sum_{s=1}^{32} n_s \left(m_{s>A} + m_{s>B} + m_{s>C}\right)$$

and $n_t$ is the number of positions in the region with reference triplet $t$. Note that for each triplet we can take $A$, $B$, $C$ as e.g. the three possible alternate alleles with respect to the reference in lexicographic order (Supp. Note 5 Fig. 1a).

**2. Expected hotspot rate at a single position**

Let's consider a stacked cohort of samples, each with identical mutation rates, and let's compute the expected number of hotspots at a specific genomic position. Let $N$ denote the number of samples and $p$ the probability that there is a mutation at a given sample. We will denote the possible alternate types at the position of interest as either $A$, $B$ and $C$, and the respective per sample probabilities as $p_A$, $p_B$ and $p_C$. Clearly, $p = p_A + p_B + p_C$.

A hotspot is called whenever the same alternate is found at the same position in at least two samples. We can think of the single position across samples as an $N$-word of base pairs of either the reference type (denoted $R$) or alternate types $\{A, B, C\}$. Following this representation, the probability $H$ of calling a hotspot in the position is the same as the probability to draw two instances of the same alternate type across samples. By the pigeonhole principle, this is equivalent to the probability of not having any of these four outcomes: i) all letters are $R$; ii) there is exactly one alternate; iii) there are exactly two distinct alternates; iv) there are exactly three distinct alternates (Supp. Note 5 Fig. 1b). Bearing this in mind, the probability to draw at least one hotspot in the position is calculated as follows:

$$H = 1 - (1-p)^N - N(1-p)^{N-1}p - 2\binom{N}{2}\left(\sum_{i\neq j} p_i p_j\right)(1-p)^{N-2} - 3!\binom{N}{3} p_A p_B p_C (1-p)^{N-3}$$

Note that this way of computing the expected number of hotspots per position yields a slight underestimation of the expected number of hotspots as per the HotspotFinder definition of hotspot, whereby at most three hotspots, corresponding to the three possible alternate alleles in the position, are possible within a single position. To account for this, only a single observed hotspot is considered per position in all calculations of observed hotspot propensities and their comparisons to the theoretical models.

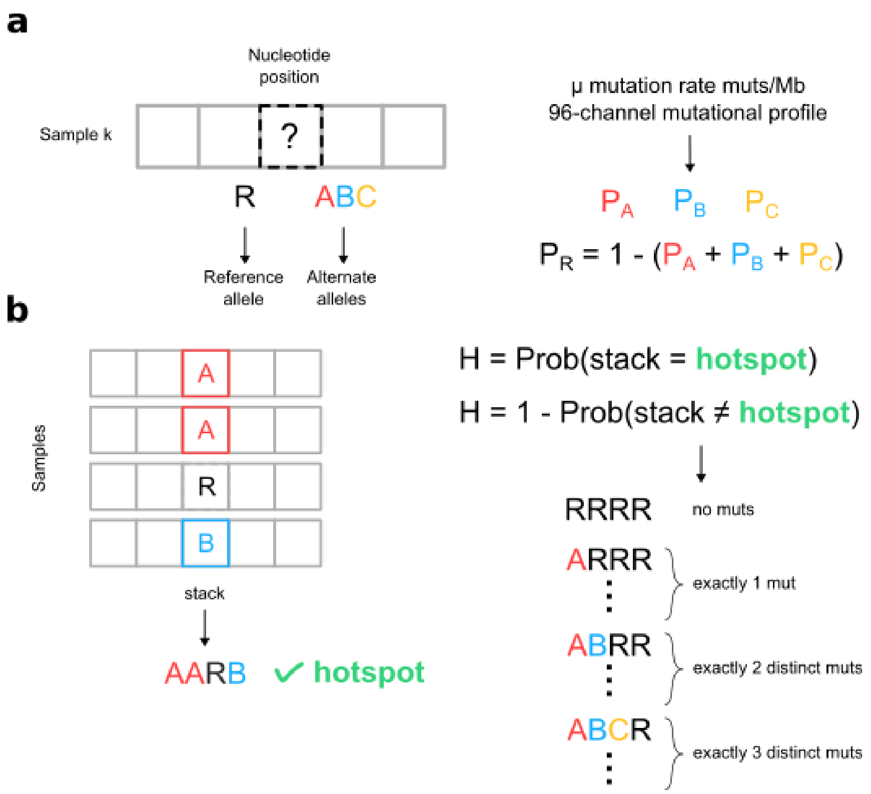


**Supp. Note 5 Fig. 1. Rationale behind the calculation of the expected number of hotspots in a single position**. **a**) First, using the mutation rate and the profile of the mutational process operative in the samples, we infer a probability for either the reference or alternate alleles to occur in a position. **b**) With this information we can then compute the probability that the position harbours a hotspot by looking at the cases when there is not a hotspot.

##### 3. Expected hotspot rate genome-wide

Making the simplifying assumption of a homogeneous mutation rate along the genome, we can compute the expected number of hotspots genome-wide --hotspot propensity-- generated by a mutational process $\underline{m} = (m_{t>a})$ with mutation rate $\mu$. By the result of the previous section, note that we will only need the triplet content of the genome, i.e. the number of positions with a given reference triplet for the 32 possible (pyrimidine-centred) reference triplets. Following the computations of the previous section, if $H(t; \underline{m}; \mu)$ denotes the expected number of hotspots at a position with reference triplet $t$ induced by the mutational process $\underline{m}$ operative with mutation rate $\mu$, then the expected number of hotspots is given by the following expression:

$$H = \sum_t n_t H(t; \underline{m}; \mu)$$

#### 4. Expected hotspot rate across mappable genomic chunks

Despite giving us a fair theoretical estimate, the previous calculation makes a strong assumption that is clearly violated by real data, namely, that the mutation rate genome-wide is homogeneous. However, we want to address a question that arises in our analyses: whether the expected hotspot rate diverges from the observed when we factor in the mutation rate variability across the genome.

To address this we computed the theoretical hotspot rate per megabase applying the same ideas depicted above with the precaution of keeping track of the relative mutation rate estimates of a mutational process per genomic chunk, i.e. how more/less mutable are regional chunks when compared with each other.

We proceed by inferring an empirical estimate of the relative mutability per genomic chunk and signature, i.e. using the mutations sequenced in real tumors. By means of signature deconstruction, for each sample in our cohort we can infer how confidently the mutations found in each of the 96 trinucleotide contexts are attributable to a specific mutational process (Supplementary Note 3). With this measure of confidence we can then count how many mutations in a cohort match a specific mutational process (maximum likelihood[13]; Methods; Supplementary Note 3), then map those mutation counts to each genomic chunk out of a set with high mappability (Methods). For estimation of the relative mutation rate of zero-count chunks we introduced a pseudocount consistent across chunk sizes whereby the pseudocount added to the chunks of size $s$ is equal to $s/1\,Mbp$: in doing so we are not changing the total mutation burden across different chunk-size analyses. With this information we can then infer the relative mutation rate attributable to a specific mutational process across chunks and apply the above described methodology to compute hotspot rates per chunk. These estimates can be finally merged to obtain the genome-wide number of expected hotspots or hotspot propensity.

#### 5. SBS1-induced hotspot rate with a DNA methylation covariate

To study the expected hotspot propensity of signature SBS1, we built models restricted to the NpCpG>T tri-nucleotide contexts (ACG>T, CCG>T, GCG>T, TCG>T). For the three cancer types analysed (COADREAD, ESOPHA_STOMACH, NSCLC) we also retrieved DNA methylation sequencing data (Methods) from which we could derive the fold-change mutation rate between methylated and unmethylated cytosines for each NpCpG>T context. Using the relative mutagenicities of the NpCpG>T contexts derived from the SBS1 96-channel profile as a baseline (4 channels), we defined three additional SBS1 models, one for each cancer type, where two relative mutagenicity values are given for each NpCpG>T context, accounting for methylated and unmethylated cytosines (8 channels).

Just as in the previous analyses, we calculated the hotspot propensity per chunk accounting for the NpCpG sites only. Since not all the bases in the mappable genome chunks were completely annotated for their DNA methylation status, for those chunks with missing values we imputed the same proportion of methylated CpGs as the proportion found genome-wide for the same cancer type. Upon estimation of the relative mutation rate per chunk, for each cancer type and chunk size we computed the expected hotspot rate with 300 mutations/sample across 100 samples.

##### 6. Numerical validation of the theoretical hotspot rate

To rule out implementation flaws and provide a stochastic hotspot generator consistent with the simplifying assumptions of the theoretical model, we conducted a round of numerical experiments with toy examples. Briefly, we generated cohorts of random synthetic samples for which the triplet content, mutation rate per sample, number of samples and mutational process (signature) generating the mutations were specified and we directly computed the number of hotspots for each cohort. To keep the computational cost low we resorted to small synthetic samples ~ 1 Kbp in size. For five different configurations characterised by the triplet content of the samples (Supp. Note 5 Fig. 2a) we generated 20 random cohorts with 50 samples and 10 mutations per sample, and compared the number of hotspots in each cohort against the theoretical hotspot rate. The proportion of triplets used to define each triplet content set-up were drawn randomly from the collection of mappable 1 Mbp chunks used elsewhere in this paper. We conducted this experiment three times with signatures SBS1, SBS17b and SBS5, respectively. In all cases, the empirical hotspot rate matched closely the expected hotspot rate provided by the theoretical model of hotspot propensity (Supp. Note 5 Fig. 2b) as intended.

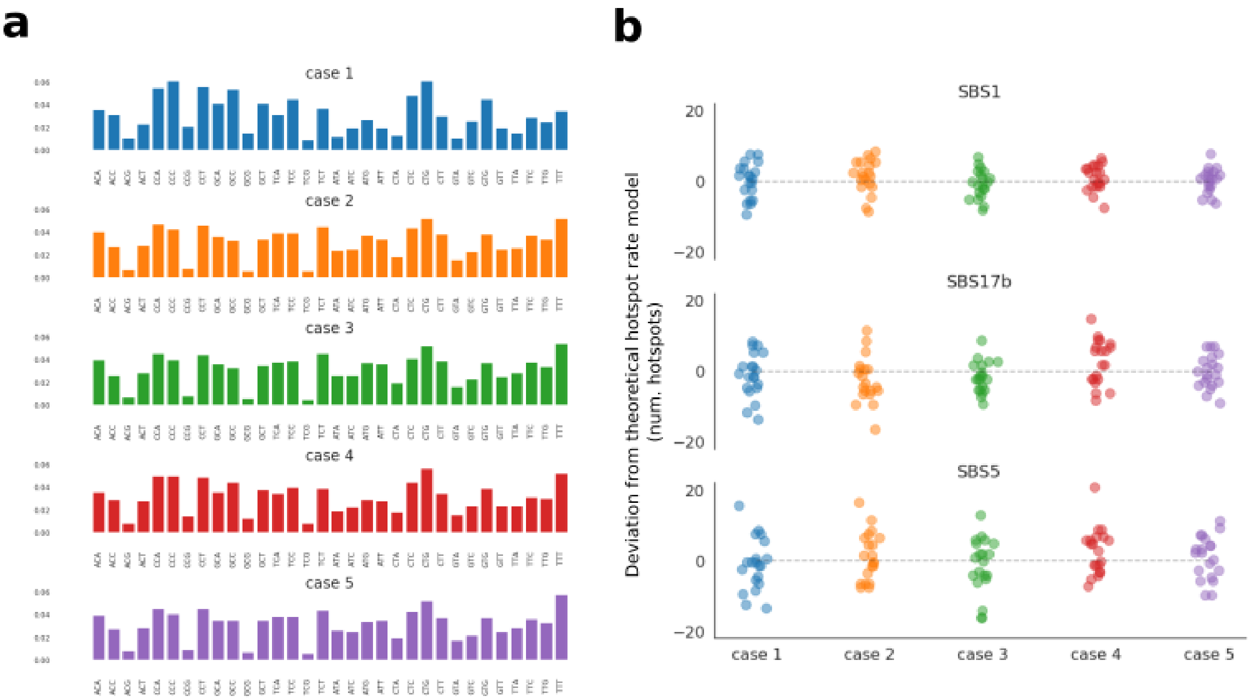


**Supp. Note 5 Fig. 2. a**) Trinucleotide content frequencies of the five case scenarios considered for the random numerical experiments and **b**) centred values after stochastic simulation with respect to the expected hotspot propensity cast by the theoretical model for the three signatures SBS1, SBS17b and SBS5. In each case we produced 20 random cohorts (points in the scatter plot) using 50 samples ~ 1 Kbp in size and 10 mutations per sample.

### Supplementary Notes References


1. Marco-Sola, S., Sammeth, M., Guigó, R. & Ribeca, P. The GEM mapper: fast, accurate and versatile alignment by filtration. *Nat. Methods* **9**, 1185–1188 (2012).
2. ENCODE Project Consortium. An integrated encyclopedia of DNA elements in the human genome. *Nature* **489**, 57–74 (2012).
3. Karczewski, K. J. *et al.* The mutational constraint spectrum quantified from variation in 141,456 humans. *Nature* **581**, 434–443 (2020).
4. Frankish, A. *et al.* GENCODE reference annotation for the human and mouse genomes. *Nucleic Acids Res.* **47**, D766–D773 (2019).
5. Alexandrov, L. B., Nik-Zainal, S., Wedge, D. C., Campbell, P. J. & Stratton, M. R. Deciphering signatures of mutational processes operative in human cancer. *Cell Rep.* **3**, 246–259 (2013).
6. Islam, S. M. A. *et al.* Uncovering novel mutational signatures by de novo extraction with SigProfilerExtractor. *Cell Genomics* **2**, None (2022).
7. Bergstrom, E. N. *et al.* SigProfilerMatrixGenerator: a tool for visualizing and exploring patterns of small mutational events. *BMC Genomics* **20**, 685 (2019).
8. Alexandrov, L. B. *et al.* The repertoire of mutational signatures in human cancer. *Nature* **578**, 94–101 (2020).
9. Alexandrov, L. B. *et al.* Signatures of mutational processes in human cancer. *Nature* **500**, 415–421 (2013).
10. Secrier, M. *et al.* Mutational signatures in esophageal adenocarcinoma define etiologically distinct subgroups with therapeutic relevance. *Nat. Genet.* **48**, 1131–1141 (2016).
11. Pich, O. *et al.* The mutational footprints of cancer therapies. *Nat. Genet.* **51**, 1732–1740 (2019).
12. Christensen, S. *et al.* 5-Fluorouracil treatment induces characteristic T>G mutations in human cancer. *Nat. Commun.* **10**, 4571 (2019).
13. Morganella, S. *et al.* The topography of mutational processes in breast cancer genomes. *Nat. Commun.* **7**, 11383 (2016).
14. Cagan, A. *et al.* Somatic mutation rates scale with lifespan across mammals. *Nature* **604**, 517–524

(2022).

15. Paulson, T. G. *et al.* Somatic whole genome dynamics of precancer in Barrett's esophagus reveals features associated with disease progression. *Nat. Commun.* **13**, 2300 (2022).
16. Rahbari, R. *et al.* Timing, rates and spectra of human germline mutation. *Nat. Genet.* **48**, 126–133 (2016).
17. An, J.-Y. *et al.* Genome-wide de novo risk score implicates promoter variation in autism spectrum disorder. *Science* **362**, (2018).
18. C Yuen, R. K. *et al.* Whole genome sequencing resource identifies 18 new candidate genes for autism spectrum disorder. *Nat. Neurosci.* **20**, 602–611 (2017).
19. Sasani, T. A. *et al.* Large, three-generation human families reveal post-zygotic mosaicism and variability in germline mutation accumulation. *eLife* **8**, (2019).
20. Halldorsson, B. V. *et al.* Characterizing mutagenic effects of recombination through a sequence-level genetic map. *Science* **363**, (2019).
21. Goldmann, J. M. *et al.* Parent-of-origin-specific signatures of de novo mutations. *Nat. Genet.* **48**, 935–939 (2016).
